# Cell heterogeneity and fate bistability drive tissue patterning during intestinal regeneration

**DOI:** 10.1101/2025.01.14.632683

**Authors:** C. Schwayer, S. Barbiero, D. B. Brückner, C. Baader, N. A. Repina, O. E. Diaz, L. Challet Meylan, V. Kalck, S. Suppinger, Q. Yang, J. Schnabl, U. Kilik, J. G. Camp, B. Stockinger, M. Bühler, M. B. Stadler, E. Hannezo, P. Liberali

## Abstract

Tissue regeneration relies on the ability of cells to undergo *de novo* patterning. While tissue patterning has been viewed as the transition from initially identical un-patterned cells to an arrangement of different cell types, recent evidence suggests that initial heterogeneities between cells modulate tissue-scale pattern formation. Yet, how such heterogeneities arise and, thereafter, regulate cell type emergence in a population of cells is poorly understood. Using *in vivo* and *in vitro* mouse regenerative systems, we identify a critical tissue density that is required to induce heterogeneous inactivation of the mechanosensor YAP1. Experimental and biophysical approaches demonstrate that YAP1 cell-to-cell heterogeneity pre-patterns the first cell fate decision, via both chromatin remodelling and a supracellular feedback between FOXA1 and Delta-Notch signalling. This feedback motif induces cell fate bistability endowing memory to the system and the maintenance of patterns during homeostasis. These findings reveal a generalisable framework in which transient cell-to-cell heterogeneity, regulated by tissue-scale properties, serves as a critical control parameter for the emergence of cell fate and stable patterning during regeneration.

## Introduction

The emergence of patterns is crucial for the spatial organization and function of tissues in development and regeneration (*1–5*). Seminal work in developmental biology has shown that initially uniform assemblies of cells are patterned through morphogen gradients (*6–14*) and direct cell-cell interactions (*15–21*). These mechanisms guide the specification of cell fates and their spatial arrangement, forming the basis for tissue organization. Similarly, during regeneration, cells undergo reprogramming and de-differentiation (*22*, *23*), losing their original fate and assuming a foetal-like identity (*24–28*). Subsequently, patterns re-emerge *de novo*, restoring spatial order and functional organisation (*3*, *4*). This process highlights striking mechanisms wherein cells must transition from an undifferentiated homogenous population of cells to a patterned structure composed of different cell types.

Classical theoretical work has considered patterning as arising from a blank canvas of identical cells (*6–8*), where stochastic fluctuations enable any cell to adopt any fate. Yet, recent evidence has uncovered that even at the earliest stages of pattern formation - when cells lack defined fates and spatial organisation - there remains substantial heterogeneity in cellular states, including mechanical, biochemical, or metabolic variability (*29–32*). Interestingly, such cell-to-cell variability is often actively regulated (*33*), begging the question of whether it serves a functional role in multicellular systems. Over recent years, studies have increasingly suggested that specific forms of heterogeneities can influence cellular behaviours and guide the initiation of patterns (*30*, *34–42*). For instance, cellular diversity in bacterial colonies underlies their ability to adapt to fluctuating resources (*31*, *32*). Similarly, variability in cortical tension can promote selective cell delamination, as observed in cardiac trabeculation (*29*), and variability in cell division timing can enhance robust patterning in mammalian embryogenesis (*43*). Further, cell-to-cell heterogeneity of the mechanosensor YAP1 (Yes-associated protein 1) regulates Delta-Notch lateral inhibition and the first cell fate decision giving rise to a proper crypt-villus axis formation in intestinal organoids (*30*).

Together, these previous observations raise two central questions: first, how are these heterogeneities established, or encoded, at the tissue-scale and second, how are tissue-scale heterogeneities subsequently read out, or decoded, by individual cells to orchestrate stable patterns in a dynamic environment?

### Tissue density controls YAP1 cell-to-cell variability and patterning during *in vivo* regeneration

To investigate how tissue heterogeneity controls pattern emergence during regeneration, we use mouse intestinal organoids, a system that enables high spatiotemporal resolution studies and that has been shown to recapitulate *in vivo* regeneration (*30*). In this model, an un-patterned organoid, composed of regenerative cells, gives rise to a subset of secretory cells named Paneth cells, a critical step for establishing the stem cell niche and the crypt-villus axis (*30*). This process requires the heterogeneous YAP1 activity across cells in the same organoid to initiate Delta-Notch lateral inhibition, and the emergence of the first DLL1 (Delta-like 1) positive (DLL1^+^) cells (*30*). To complement this *in vitro* system, we employed an *in vivo* mouse model of irradiation-induced intestinal regeneration (10Gy) (*44*, *45*) and we tracked the process: at the onset (24h post irradiation), cells exhibited YAP1 activation and a loss of canonical epithelial lineage markers (Fig. 1A, fig. S1A). Following this YAP1-mediated reprogramming phase (*22*), secretory progenitors were the first cell type to emerge, peaking at 48h post irradiation. Stem cells and absorptive cells subsequently appeared, reaching 20% and 50% of their homeostatic cell type abundance by 48h post irradiation, respectively (Fig. 1A, fig. S1A). The same spatiotemporal sequence of events was observed in *in vitro* organoid growth, where an initial increase of YAP1 activity (increased nuclear translocation of YAP1, read out by mean intensity - total intensity divided by area - of active YAP1 antibody) precedes the emergence of secretory DLL1^+^ progenitors (*30*), setting the stage for the re-establishment of homeostasis (Fig. 1B, fig. S1D). These parallels between *in vitro* and *in vivo* systems underscore the conserved role of YAP1 in driving pattern emergence during intestinal regeneration (*30*, *46*) and how intestinal organoid formation mimics the spatiotemporal sequence of regenerative events of the epithelium.

**Figure 1:**
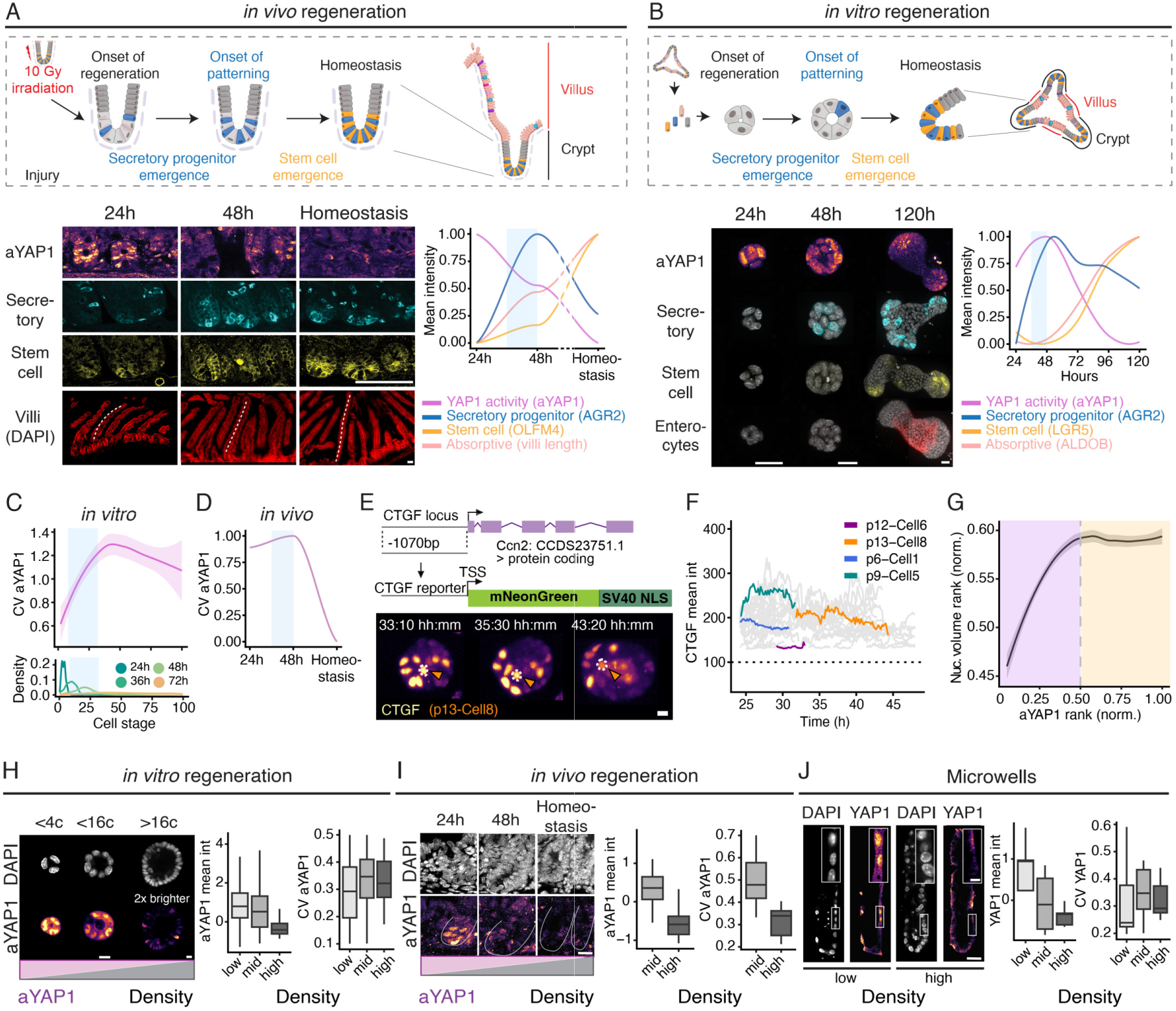
Tissue density controls YAP1 cell-to-cell variability and patterning during *in vivo* regeneration (A) Scheme of *in vivo* regeneration (top). On the left, z-planes of active YAP1, AGR2 (secretory progenitor marker) and OLFM4 (stem cell marker) immunostainings, and Maximum intensity projection (MIPs) of nuclear labelled DAPI show cross section of crypt-villus axis in small intestine, Duodenum (white lines indicate length of villi). For better display, 24h DAPI stain in villi is shown 6x brighter and at 48h 2x brighter compared to homeostasis. On the right, line plot shows mean intensities of cell type markers and YAP1 activity. Blue rectangle indicates the time of secretory progenitor emergence (36-48h). Scale bars, 100µm. Dashed lines separate regenerating timepoints from homeostasis. N = 3 mice, n = 3,118 cells (between 170 and 300 cells per time point per staining). For each time point and each staining 9 to 20 crypts have been analysed. For villi length each time point counts 25 to 30 villi. Unsmoothed data in fig. S1A. (B) Scheme of *in vitro* regeneration (top). On the left, MIPs of active YAP1, AGR2 (secretory progenitor marker) staining, Lgr5::DTR-EGFP (stem cell marker) in organoids are shown. On the right, line plot shows mean intensities of the cell type markers and YAP1 activity. Blue rectangle indicates the time of secretory progenitor emergence (36-48h). Scale bar, 10µm. n = 8,954 organoids. (C, D) Line plots show coefficient of variation (CV) of classified aYAP1 as a function of cell stage *in vitro* (C) or timepoint *in vivo* (D). Blue rectangle indicates the time of secretory progenitor emergence (36-48h). Lower plot in (C) shows density distribution of cell stage per time point. N = 3 experiments, n = 1,727 organoids, N = 9 mice from three separate timepoints (24h, 48h, homeostasis), n = 19 crypts from 24h, 9 crypts from 48h, and 15 crypts from homeostasis. (E) Scheme of CTGF reporter line design. MIPs of CTGF-mNeonGreen-NLS signal during light sheet imaging. (F) Line plot of CTGF mean intensity over time. Lines indicate tracks of single cells. Single cells were tracked for a period of one cell division. Dotted line indicates background threshold. Scale bar, 10µm. n = 89 single cells tracked across 4 organoids. (G) Plot shows nuclear volume ranked from lowest to highest values per organoid as a function of active YAP1 rank per organoid. Organoids from 8-20 cell stage were analysed and ranks were normalized for the exact cell stage. N = 3 experiments, n = 3,371 organoids. (H) Z-plane of DAPI and aYAP1 staining from 4-cell (4c) to more than 16-cell stage (> 16c). Boxplots show z-scored mean intensity and coefficient of variation of aYAP1 per different cell density bins (low < 0.3, mid < 0.7, high > 0.7 number of nuclei per 100µm^2^). aYAP1 z-plane at > 16c is displayed 2x brighter compared to other cell stages for better visualisation. N = 3 experiments, n = 5,065 organoids. Scale bar, 10µm. (I) Z-plane of DAPI and aYAP1 staining of *in vivo* tissue sections of intestinal crypts (white lines outline each crypt). Boxplots show z-scored mean intensity and coefficient of variation of aYAP1 per different cell density bins. N = 9 mice from three separate timepoints (24h, 48h, homeostasis), n = 27 crypts for mid-density bin, and 16 crypts for high density bin. Scale bar, 20µm. (J) Z-plane of DAPI and total YAP1 staining of engineered microwells with different amounts of cells and densities. Inset shows 4x zoom in. Plots show z-scored mean intensity of total Yap1 staining in the nucleus and CV of nuclear YAP1 per different cell density bins. N = 3 experiments, total number of microwells analysed, n = 27 (5 in low density bin, 15 in mid density bin, 7 in high density bin). Scale bar, 10µm, 2.5µm (zoom-in).

To analyse the cell-to-cell heterogeneity of the regenerative response in more detail, we developed an automated analysis pipeline capable of segmenting single cells and extracting both cellular- and tissue-level features from large numbers of samples across multiple time points (*47*, *48*) (fig. S1B, >50K single cells analysed). Secretory cells emerged consistently at the 8- to 16-cell stages, corresponding to the 36-48h window in organoids (Fig. 1B, fig. S1C, (*30*)), and at 48h *in vivo* (Fig. 1A). In this time window, YAP1 becomes heterogeneously inactivated both *in vitro* and *in vivo*, before returning to homogeneously low levels during homeostasis (Fig. 1A-D). Moreover, cell-to-cell heterogeneity of YAP1 activity, quantified as the coefficient of variation of classified YAP1 activity per organoid or crypt, shows a steep increase in concomitance with secretory cell emergence between 24-48h *in vitro and in vivo* (Fig. 1C,D, blue rectangle represents the time of secretory progenitor emergence).

To gain further insights into the heterogeneous inactivation dynamics of YAP1, we engineered a fluorescent reporter based on the YAP1 target gene CTGF and validated its functionality (fig. S1H,I). Using this reporter with light sheet imaging and single cell tracking, we observed that cells do not oscillate in the YAP1 activity, but every cell loses YAP1 activity over time. We therefore conclude that cell-to-cell heterogeneity in YAP1 activity arises from differences in deactivation rates among individual cells (Fig. 1E,F). To determine whether YAP1 activity correlates with cell features, we conducted high-throughput imaging analysis of YAP1 activity in organoids assessing single cells at the 8-32 cell stage. Heterogeneity in nuclear volume and elongation and, the nuclear stiffness marker Lamin A/C (*49*), increases with cell stage mimicking the behavior of YAP1 heterogeneity (fig. S1E-G). At the single cell level, ranking cells according to YAP1 activity and related intrinsic features revealed that the cells with larger nuclei, greater cell volume, and more elongated nuclei (higher aspect ratio) tend to show higher YAP1 activity within the same organoid (Fig. 1G, fig. S1K).

Nuclear and cell size decrease as epithelial volume and tissue density increase (fig. S1E-G, J). This prompted us to explore whether tissue density regulates YAP1 cell-to-cell variability. Remarkably, YAP1 is homogeneously active at low density, heterogeneously inactivated at intermediate density, and homogeneously inactive at high density both in organoids and in *in vivo* regeneration (Fig. 1H,I, fig. S1L-N,P). This suggests that YAP1 heterogeneity is established or encoded at a specific tissue density giving a permissive time window to restore homeostasis. To test this, we controlled tissue density at constant tissue curvature, using bioengineered scaffolds at varying cell numbers (*50*). Indeed, YAP1 activity was inversely correlated with tissue density, with maximum cell-to-cell heterogeneity observed at intermediate density levels (Fig. 1J, fig. S1O). Importantly, YAP1 heterogeneity is established at a regime of intermediate density that coincides with the time window at which tissue integrity is restored just prior to the establishment of the first secretory cell, both during *in vitro* and *in vivo* regeneration (Fig. 1A-D). This specific density regime is important for patterning, as not patterned organoids exhibit a lower tissue density compared to patterned organoids (fig. S1Q,R). Functionally perturbing tissue density either via inhibiting cell division (Aphidicolin, Taxol) (fig. S1S,U) or by inducing lumen inflation in organoids (Forskolin (*51*) or Prostaglandin E2 (PGE) (*52*) (fig. S1V,W) prevented DLL1 emergence at low tissue density (fig. S1S-W). DLL1 emergence could be partially rescued after removing the cell division inhibition (fig. S1T,U).

Taken together, these findings reveal that tissue density controls YAP1 (*53*, *54*) heterogeneous inactivation (Fig. 1J) as well as DLL1 emergence (fig. S1S-W), suggesting that tissue-scale properties control the timing of tissue repair, defining a window of opportunity for patterning.

### YAP1 heterogeneity and DLL1 bistability drive patterning

Having established that YAP1 heterogeneity is encoded at an intermediate tissue density, we next explored how this YAP1 heterogeneity is decoded at the single-cell level to give rise to a stereotypic patterning outcome. In organoids, YAP1 heterogeneity emerges at early cell stages concomitantly with the increase in DLL1^+^ cell fraction (Fig. 2A,B). To test the functional role of YAP1 heterogeneity, we manipulated YAP1 activity using temporally controlled perturbations: YAP1 inhibition with Verteporfin (from 36h) or YAP1 activation (from 0h) with a selective LATS kinase inhibitor (*46*). These perturbations resulted in either homogeneously low or homogeneously high tissue-wide YAP1 activity, respectively (Fig. 2D,E, fig. S2A). Both treatments resulted in a reduction of DLL1^+^ cells and disrupted the patterning (Fig. 2D,F, fig. S2B). However, while YAP1 inhibition fully abolished DLL1 expression, YAP1 activation induced a homogenous intermediate expression of DLL1, with no cell fully specified into DLL1^+^ fate (Fig. 2D, fig. S2G). Notably, in wild-type organoids, single-cell YAP1 activity correlates with DLL1 expression (Fig. 2C), further supporting a functional link between YAP1 and DLL1.

**Figure 2:**
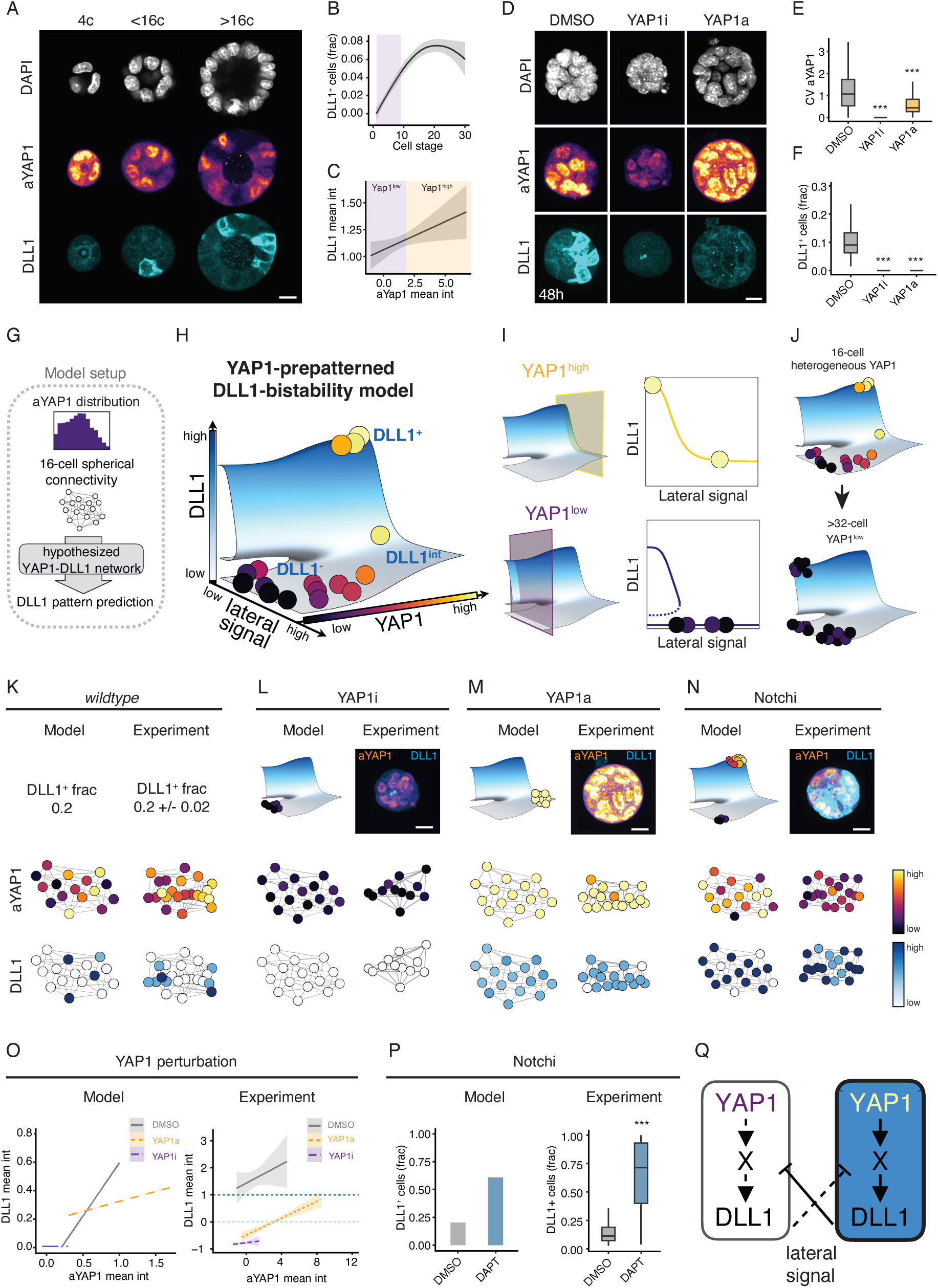
YAP1 heterogeneity and DLL1 bistability drive patterning (A) Z-planes of DAPI, aYAP1, DLL1 staining at different cell stages. Scale bar, 10µm. (B) Line plot of the mean DLL1^+^ cell fraction as a function of cell stage. Mean intensities +/− 90% confidence interval. Purple rectangle indicates emergence of YAP1 heterogeneity. N = 3 experiments, n = 5,210 organoids. (C) Line plot of z-scored DLL1 and aYAP1 mean intensities per organoid at 8 to 20-cell stage. Mean intensities +/− 90% confidence interval. N = 3 experiments, n = 340 DLL1 patterned organoids. (D) MIPs of DAPI, aYAP1, DLL1 of DMSO control, YAP1i (inhibitor, Verteporfin), YAP1a (activator, LATSi) at 48h. Scale bar, 10µm. (E) Boxplot of CV of classified aYAP1 in different YAP1 perturbations at 48h. N = 3 experiments, n(DMSO) = 166, n(YAP1i) = 891, n(YAP1a) = 579 organoids. (F) Boxplot of DLL1^+^ cell fraction per organoid in different YAP1 perturbations at 48h. N = 3 experiments, n(DMSO) = 166, n(YAP1i) = 891, n(YAP1a) = 579 organoids. (G) Illustration of our model setup: the experimentally measured distribution of YAP1 activity of patterned organoids with 8-20 cells is used to generate heterogeneous networks with the connectivity of a 16-cell sphere based on which DLL1 patterning is predicted. Throughout, network plots are generated by projection of spherical coordinates on a 2D plane, and nearest neighbour connections are indicated with grey lines. (H) YAP1-prepatterned DLL1 bistability model: a plane of stable DLL1 attractor states is defined within each single cell as a function of this cell’s YAP1 activity and the lateral inhibition signal sent from neighbouring cells. Stable cellular states of DLL1 are fixed points of the dynamics, conceptually equivalent to attractors in Waddington’s landscape. Blue colouring indicates the DLL1 state and is proportional to the vertical axis of the plot (bottom left). The locations of points of a simulated 16-cell organoid are indicated on this plot and correspond to the same organoid as shown in panel (K). Point colours indicate YAP1 level. The gap in the surface along the DLL1 axis for YAP1^low^ and low lateral signal indicates the activation barrier separating the bistable states. (I) Cuts through the attractor surface at high and low YAP1 activity, respectively, with locations of low and high YAP1 cells from panel (H) indicated. (J) Equivalent attractor surface plots visualised for the time-course of organoid development, in which 16-cell organoids exhibit heterogeneous YAP1 levels leading to patterning, and organoids beyond the 32-cell stage exhibit low YAP1. In the latter case, the progression of initially high YAP1, and thus DLL1^+^ cells towards the bistable DLL1^+^ attractor is indicated. (K) Network plots of simulated (left) and experimental (right) wildtype organoids with fraction of DLL1^+^ cells per organoid. Mean +/− SEM. N, n for experiment same as in Fig. 2C. (L, M, N) DLL1 attractor surface and network plots for YAP1 inhibition (L), YAP1 activation (M) and Notch perturbation (N) on the left and merged MIPs of aYAP1 and DLL1 and experimental network plots on the right. Perturbation of lateral inhibition signalling in the model is implemented as cells being insensitive to any laterally received signal. (O) Correlation of single-cell z-scored mean intensity of DLL1 and aYAP1, shown for organoids from 8-20 cell stage for control, YAP1 activation, and YAP1 inhibition experiments. Model output (left) and data from one representative experiment (right). n(DMSO) = 18, n(YAP1i) = 79, n(YAP1a) = 44 organoids. (P) Bar plot shows the fraction of DLL1^+^ cells for model (left) and boxplot shows the fraction of DLL1^+^ cells for experiment (right). N = 3 experiments, n = 351(DMSO), n(DAPT) = 838 organoids. (Q) Schematic of the YAP1-prepatterned DLL1 bistability model. Statistical significance is indicated with *** (p < 0.001).

To uncover the mechanisms linking YAP1 heterogeneity to DLL1 patterning we employed our quantitative organoid multidimensional single-cell dataset to constrain a mathematical model of tissue patterning. Classical models have explained DLL1 patterning as arising from lateral inhibitory signals sent from neighbours as a function of their DLL1 levels (*16–18*, *55*). Here, we aim to define the minimal regulatory network linking tissue-level features (DLL1-dependent lateral signals, defined as the average neighbouring DLL1 levels (fig. S2E), and organoid connectivity) to cell-to-cell variability (YAP1 activity) (*16–18*, *56*). Focusing on the 16-cell stage, when the initial patterning event emerges (Fig. 2A,B), we generated heterogeneous networks of cells (nodes in the network) connected with their direct neighbouring cells (links in the network) with YAP1 activities distributed according to the experimentally measured YAP1 distribution (Fig. 2G, SI Theory section 1). Using these networks as input to simulations of hypothesised YAP1-DLL1 regulatory interactions, we predicted DLL1 patterns as output of the model (Fig. 2G). We tested two initial hypotheses combining YAP1 regulation of DLL1 with classical lateral inhibition (*16–18*), in each case systematically screening a broad space of relevant model parameters (SI Theory section 2-3). In the first scenario, YAP1 directly pre-patterns DLL1 autonomously within cells (fig. S2C). While this model successfully explained the YAP1 activation experiment, it incorrectly predicted the emergence of DLL1^+^ cells under YAP1 inhibition (Fig. 2D-F). In the second model, YAP1 pre-patterns lateral interactions, meaning that YAP1 activity modulates how much DLL1 is produced for a given amount of lateral signal (fig. S2D). This model correctly predicted outcomes of YAP1 inhibition but failed to reproduce the YAP1 activation experiment (Fig. 2D-F). These discrepancies suggested that neither simple cell-autonomous nor purely interaction-based mechanisms sufficed to explain the experimental data for all parameters set.

We next proposed a more complex model where YAP1 activates cell-autonomous DLL1 production and regulates lateral interactions simultaneously. With this assumption, we indeed captured both YAP1 inhibition and activation experiments across a broad parameter space (SI Theory section 3). Importantly, under YAP1 activation, the model predicted the loss of DLL1 patterning as for the inhibition, but with the key difference that it displays uniform intermediate DLL1 levels, because cells mutually block patterning through signalling to one another. Notably, imaging confirmed this prediction showing a higher fraction of un-patterned organoids (fig. S2B) and increased number of organoids with intermediate DLL1 levels (fig. S2G). However, a limit of this model is that, for any YAP1 activity, there is a single equilibrium between DLL1 levels and signals from the neighbours. Consequently, this model alone could not account for the observed maintenance of DLL1 expression after YAP1 downregulation during homeostasis in late organoids, suggesting a “memory” effect of prior YAP1 heterogeneity. The simplest theoretical interpretation of this “memory” effect is that there are additional regulatory interactions ensuring that YAP1^low^ cells must exhibit a second stable DLL1^+^ state, i.e. a cell-intrinsic bistability in DLL1 levels (*18*). Adding this feature to our model (SI Theory section 3) results in the stable cellular states of DLL1 depending in a non-trivial manner on both the YAP1 activity and the lateral signal sent by neighbours (Fig. 2H,I). Specifically, cells with low YAP1 activity exhibit two stable states: a DLL1^−^ state and a DLL1^+^ state, separated by a barrier (Fig. 2H,I). YAP1^high^ cells, in contrast, exhibit a single stable DLL1^+^ state at low lateral signal, and an intermediate DLL1 state at high lateral signal, showing that YAP1 activity is the control parameter of the system and enables a transition between different bistable states. In order to reach the DLL1^+^ state at low lateral signal and low YAP1 activity (e.g. at > 32-cell stage), cells must transition through a YAP1^high^ state (e.g. at 16-cell stage) (Fig. 2J).

This model quantitatively captures the wild-type fraction of DLL1^+^ cells (Fig. 2K), both YAP1 perturbations (Fig. 2L,M, fig. S2F), and single-cell correlations between YAP1 and DLL1 (Fig. 2O). Importantly, this analysis also demonstrates that the YAP1 activity of cells in the YAP1 activator experiment is within physiological, yet uniformly high levels (Fig. 2E,O). To experimentally validate the model without altering the cell intrinsic YAP1 activity, we perturbed lateral inhibition. In particular, an intuitive prediction of the model is that the DLL1^+^ fraction should increase if the signal exchange between neighbouring cells is perturbed. Indeed, this prediction was confirmed by using a Notch signalling inhibitor (gamma-secretase inhibitor, DAPT (*57*)) (Fig. 2N,P). Upon Notch inhibition, the spatially distinct patterning of Delta-Notch is perturbed changing from a salt-and-pepper pattern of DLL1^+^ cells neighbouring HES1^+^ cells to an increase in DLL1^+^ cells (fig. S2H,I). These findings underscore how bistability locks the pre-patterned heterogeneity into stable fates, tightly linking tissue-level YAP1 heterogeneity to cell-intrinsic patterning mechanisms. The model raises the intriguing question of the mechanistic implementation of this YAP1-mediated pre-patterning. From a theoretical perspective, an intermediate species (denoted X in Fig. 2Q), which links YAP1 to DLL1 and is inhibited by DLL1 in neighbouring cells, could provide a mechanistic implementation of our heterogeneity-driven bistability model (SI Theory Section 4).

### YAP1 primes cells for secretory fate via chromatin accessibility

To uncover the molecular mechanisms decoding the heterogeneity in single-cell YAP1 activity and potentially integrating pre-patterning during intestinal organoid regeneration, we performed single-cell Assay for Transposase-Accessible Chromatin using sequencing (scATAC-seq) profiling 66,976 cells across 252,049 peaks (Fig. 3A,B). This approach was motivated by YAP1’s known role as a transcriptional activator and modulator of the chromatin state critical for cellular identity and differentiation (*58*). We used intestinal organoids as they provide the temporal resolution necessary to identify the diverse cell states involved in the transient regeneration process, to a level of resolution that would be impossible to achieve *in vivo*. Comparing chromatin accessibility in regenerating organoids with published *in vivo* data from intestinal regeneration post stem cell ablation, we found that our *in vitro* system recapitulated well the chromatin accessibility observed *in vivo* (fig. S3A,B). We then annotated the major intestinal lineages based on the predicted expression of known marker genes (Fig. 3C, and fig. S3C,D). At early stages (24h to 36h, prior to DLL1 emergence), we identified two main clusters. The first cluster, termed reprogramming cells, exhibited the coexistence of intestinal epithelial markers (e.g. *Alpi*, *Aldob*, *Apoa1*) and classical foetal and regenerating markers (e.g. *Yap1*, *Areg*, *Anxa3*, *Ajuba*) (*22*, *30*, *44*, *59*), while lacking canonical stem cell markers (e.g. *Lgr5*, *Olfm4*, *Smoc2*, *Ascl2*). In contrast, the second cluster, termed regenerative cells, showed a loss of epithelial signatures, maintaining accessibility of foetal and regenerating markers accompanied by accessibility gain in stem cell markers (fig. S3C,D).

**Figure 3:**
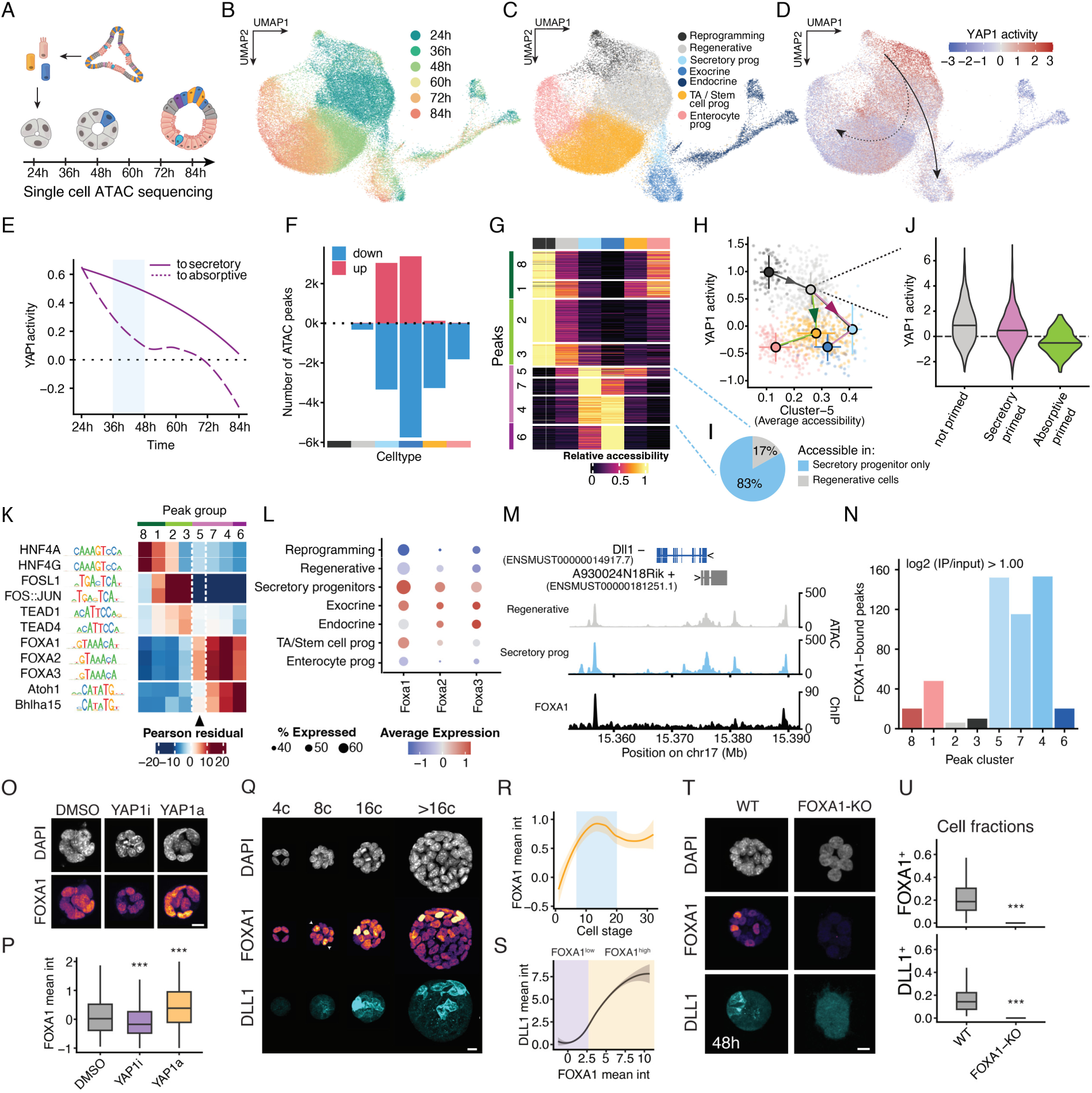
YAP1 primes cells for secretory fate via chromatin accessibility (A) Schematic drawing of single cell ATAC sequencing time course in intestinal organoid regeneration. (B, C) Uniform Manifold Approximation and Projection (UMAP) of the single cell ATAC sequencing data. Cells are coloured by experimental time (B) and assigned cell type (C). (D) UMAP embedding coloured by inferred YAP1 activity. Arrows are manually drawn according to predicted gene expression progressing from reprogramming cells to absorptive (dashed) or secretory lineage (continuous). (E) Line plot showing average YAP1 activity per time point in reprogramming-to-absorptive (dashed: reprogramming, regenerative, TA / stem cell progenitors, enterocyte progenitors) and reprogramming-to-secretory trajectories (continuous: reprogramming, regenerative, secretory progenitors, exocrine). Light blue box highlights timepoints of patterning (36-48h). (F) Bar plot showing number of peaks gaining (red) or losing (blue) accessibility (absolute log fold change > 0.81) compared to reprogramming cells. (G) Heatmap of peaks changing accessibility across cell types. Peaks are row scaled from 0 to 1 to highlight relative changes of accessibilities. Peaks are clustered (k-means clustering) in 8 groups based on their accessibility profile across cell types. (H) Scatter plot of YAP1 activity and average accessibility in peak cluster 5. Points are coloured according to cell type annotation. For robust estimates, single cells are grouped in meta-cells (average of 30 single cells), respecting cell type and experimental time point boundaries. Cell type centroids and error bars are calculated as median +/− mad. Connecting segments guide the lineage progression according to developmental time and predicted gene expression. (I) Pie chart of secretory progenitor peaks (groups 5-7-4) showing number of peaks uniquely accessible in secretory progenitor cells (groups 7-4, light blue), and number of peaks with accessibility in regenerative cells (group 5). (J) Violin plots of YAP1 activity in regenerative cells at 36-48 hours. Regenerative cells are stratified in secretory primed (highest accessibility in peak group 5, magenta), absorptive primed (highest accessibility in peak group 8 and low accessibility in peak group 3, green), and not primed (all others). (K) Heatmap of motif enrichment analysis in peak groups of interest. Groups of motifs are visually separated by white lines based on their enrichment profile across groups of peaks. Representative motifs for each group are shown. For full results see (fig. S3E). (L) Dot plot of predicted average expression of FOXA transcription factors across cell types. (M) Representative ATAC and ChIP sequencing signal in regenerative and secretory progenitor cells around DLL1 locus. (N) Bar plot of number of FOXA1 bound peaks in peak groups of interest identified from single cell ATAC sequencing. Peaks are defined as bound when log2 (IP / input) > 1.00. n = 2 replicates. (O) MIPs of DAPI and FOXA1 in early organoids, prior to patterning (< 8 cell-stage), in DMSO, YAP1 perturbations. N = 3 experiments, n(DMSO) = 456, n(YAP1i) = 972, n(YAP1a) = 1134 organoids. (P) Boxplot of FOXA1 organoid-level z-scored mean intensity in early organoids (< 8 cell-stage). (Q) MIPs of DAPI, FOXA1 and DLL1 in organoids at different time points after plating. Arrowheads at 36h indicate nuclear FOXA1 signal preceding DLL1 emergence. Scale bar, 10µm. (R) Line plot of z-scored FOXA1 mean intensity per cell stage. Blue shading indicates DLL1 emergence between 8-16 cell stage. N = 3 experiments, n = 960 organoids. (S) Line plot of z-scored DLL1 and FOXA1 mean intensities per organoid at 8 to 20-cell stage in DLL1 patterned organoids. Mean intensities +/− 90% confidence interval. N = 3 experiments, n = 195 organoids. (T) MIPs of wild-type and FOXA1-KO organoids at 48h. Scale bar, 10µm. (U) Boxplot (top) of FOXA1^+^ fraction in FOXA1-KO organoid line (FOXA1-KO organoids were identified by the absence of FOXA1 signal in polyclonal line). N = 3 experiments, n(wild-type, WT) = 263, n(FOXA1-KO) = 331 organoids. Boxplot (bottom) of DLL1^+^ cell fraction for WT organoids and FOXA1-KO organoids. Statistical significance is indicated with *** (p < 0.001).

These data suggest that YAP1 activity drives a transition from mature intestinal cells to a progenitor-like state, reopening stem cell-like chromatin regions. From this regenerative state, cells diverge into two primary fates: secretory progenitor (light blue, Fig. 3C), or stem-like transit amplifying (TA, yellow), characterised by reinforced accessibility at stem-like regions, ultimately maturing into absorptive progenitors (pink). This process underpins the lineage bifurcation into secretory and enterocyte progenitors. To connect YAP1 activity to chromatin dynamics, we inferred a single-cell YAP1 activity score (*30*) based on the predicted expression of YAP1 target genes derived from gene body accessibility (*60*). Cells committing to secretory fate retained higher YAP1 activity compared to absorptive lineages, aligning with our model of YAP1 pre-patterning DLL1 expression (Fig. 3D,E). Quantifying chromatin changes during fate acquisition revealed a stark asymmetry: while TA/stem cell and absorptive progenitors predominantly lost chromatin accessibility, secretory progenitors displayed both losses and gains of accessible regions (Fig. 3F). This asymmetry in chromatin accessibility suggests a default mode of intestinal differentiation, whereby Notch-mediated fates (e.g. absorptive) stem from loss of chromatin accessibility, while DLL1-driven secretory differentiation requires both losses and gains of accessibility.

To further understand the priming of chromatin accessibility mediated by YAP1 we clustered differentially accessible peaks into eight groups based on cell type-specific accessibility (Fig. 3G). Notably, 17% of secretory progenitor-specific regions gained accessibility in regenerative cells (group 5), thereby providing accessible regions linking YAP1-priming at the chromatin level to secretory fate. To further investigate YAP1’s role in priming cells toward the secretory fate, we sub-clustered regenerative cells into secretory pre-patterned cells (with the highest accessibility in group 5) and absorptive pre-patterned cells (with the highest accessibility in group 8). Secretory pre-patterned cells exhibited higher YAP1 activity than absorptive pre-patterned cells, reinforcing the link between YAP1 activity and lineage specification (Fig. 3J).

To identify putative transcription factors mediating these changes in chromatin accessibility we did motif analysis in accessible peaks and identified transcription factors (TFs) associated with specific groups and lineages (fig. S3E). Clustering transcription factors by motif similarity, revealed distinct enrichment patterns (Fig. 3K, and fig. S3F). HNF4-like motifs were enriched in groups 8 and 1, corresponding to the absorptive lineage. AP1-like motifs (FOS-JUN) were enriched in groups 2 and 3, suggesting roles in general reprogramming processes. Fo secretory lineage-associated peaks, ATOH1-like motifs were enriched in groups 7, 4, and 6, representing mature and progenitor secretory cells. Strikingly, group 5, which is associated with the pre-patterned secretory state, lacked enrichment in the ATOH1 motif, but uniquely displayed significant enrichment in FOXA motifs. This suggested FOXA TFs as key mediators of the transition from the YAP1-primed state to the secretory lineage at the chromatin level.

Among FOXA factors, we identified FOXA1 as a key candidate due to its specific predicted expression in secretory progenitor cells, compared to endocrine cells (Fig. 3L, and fig. S3G). Chromatin immunoprecipitation coupled with sequencing (ChIP-seq) further confirmed the specific binding of FOXA1 to secretory progenitor loci (groups 5, 7, and 4), importantly including the DLL1 locus and ATOH1 (Fig. 3M,N, and fig. S3H,I). To further validate that YAP1 pre-patterns the secretory fate through FOXA1 expression, we perturbed YAP1 activity in organoids prior to Delta-Notch patterning and imaged them (< 8 cell-stage). Consistent with our hypothesis, FOXA1 mean intensity was reduced upon YAP1 inhibition, but increased in YAP1 activation, further supporting a molecular link between YAP1 and FOXA1 (Fig. 3O,P). Altogether, our data support a model in which YAP1 activity primes the chromatin by a broad increase in accessibility followed by a chromatin state favourable for FOXA1 binding in cells that retain YAP1 activity.

To then investigate how FOXA1 regulates secretory identity, we stained and imaged FOXA1 and DLL1 simultaneously at various time points. FOXA1 was first detected at the 4-8 cell stage, prior to the emergence of DLL1 cells (Fig. 3Q,R). Consistent with the hypothesis that FOXA1 is critical for DLL1 acquisition, FOXA1 positive (FOXA1^+^) cells exhibit significantly higher DLL1 mean intensity at 8-20 cell stage (Fig. 3S) and FOXA1 signal was markedly increased in patterned organoids (fig. S3J,K). To directly test whether FOXA1 is necessary for DLL1 patterning, we generated a FOXA1 knockout organoid line (fig. S3L). FOXA1-deficient organoids showed a complete failure to establish DLL1 patterning, confirming the essential role of FOXA1 in the emergence of DLL1^+^ secretory cells (Fig. 3T,U).

Remarkably, although FOXA TFs are established players in embryonic development (*61*) and secretory differentiation within the gut (*62*), our study reveals their pivotal role in intestinal regeneration. Together, these findings demonstrate that YAP1-induced chromatin accessibility establishes a permissive environment for FOXA1 binding, enabling it to decode YAP1 heterogeneity directing the specification to secretory cell fates.

### FOXA1-mediated bistability decodes YAP1 pre-patterning

Having established FOXA1 as a key regulator linking YAP1 to DLL1 (Fig. 4A), we hypothesised that it contributes to the regulatory interactions needed to implement DLL1^+^ bistability. We then tested whether our model could accurately predict FOXA1 patterning under different experimental conditions (Fig. 4D). More specifically we incorporated FOXA1 into our model, where it not only mediates YAP1 regulation of DLL1 but also simultaneously integrates lateral signals from neighbouring DLL1^+^ cells (SI Theory section 4, 5).

**Figure 4:**
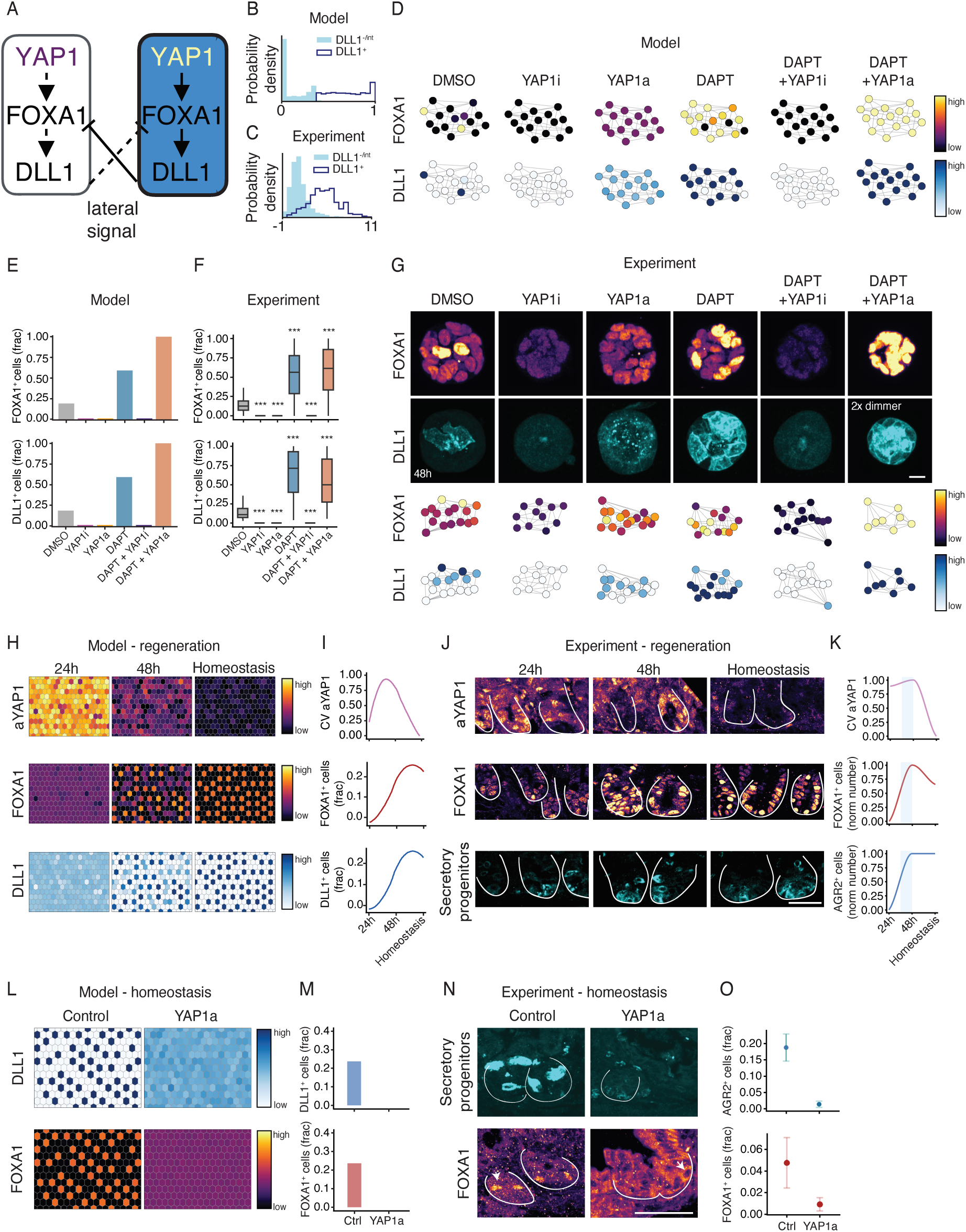
FOXA1-mediated bistability decodes YAP1 pre-patterning and confers patterning memory (A) Schematic of the YAP1-prepatterned DLL1 bistability model including FOXA1 as an intermediate variable. (B, C) Predicted (B) and experimental (C) wildtype distribution of FOXA1 levels of DLL1^−^/DLL1^int^ (light blue) and DLL1^+^ cells. (D) Network plots of model predictions for FOXA1 (top) and DLL1 (bottom) for all experimentally sampled perturbations. (E) Bar plot of predicted fraction of FOXA1^+^ and DLL1^+^ cells per organoid for each condition. (F) Boxplot of experimentally measured fraction of FOXA1^+^ and DLL1^+^ cells per organoid for each condition. N = 3 experiments, n(DMSO) = 351, n(YAP1i) = 1,352, n(YAP1a) = 1,689, n(DAPT) = 838, n(DAPT + YAP1i) = 1,367, n(DAPT + YAP1a) = 636 organoids. (G) MIPs of FOXA1 and DLL1 signal in organoids for various perturbation experiments (top). MIP of DLL1 staining of DAPT + YAP1 activator is shown 2x dimmer for better display of single-cell signal. Scale bar, 10µm. Network plots of 3D segmented, 2D projected experimentally measured FOXA1 levels (top) and DLL1 classification (bottom). (H) Model predictions for *in vivo* regenerative time-course experiments. The YAP1-levels evolved similarly to experiment (uniform high to heterogeneously downregulated to uniform low, see SI Theory section 6), and resulting FOXA1 and DLL1 patterns (mean positive fractions) are predicted, using the model parameters constrained from organoid experiments. (I) Time-evolution of mean fraction of positive FOXA1 and DLL1 cells and coefficient of variation calculated from positive/negative cell classification for aYAP1. (J) MIPs of aYAP1, FOXA1 and AGR2 (marker for secretory progenitors) at 24h, 48h post 10Gy irradiation and at homeostasis (sample prior to irradiation). (K) Time-evolution of number of positive FOXA1 and DLL1 cells and coefficient of variation calculated from positive/negative cell classification for aYAP1. Y axes are min-max scaled from 0 to 1. Smoothed curves are estimated with a loess function. N = 3 mice, n = 9-20 crypts per timepoint. Unsmoothed data in (fig. S4D). (L) Model prediction for *in vivo* YAP1 activation, obtained by rapid increase of YAP1 input levels starting from a homeostatic patterning state, which leads to homogeneous decrease of both FOXA1 and DLL1. (M) Bar plot of mean fraction of FOXA1 and DLL1 positive cells. (N) Z-planes of AGR2 (secretory progenitor marker) and FOXA1 in Control (Vehicle) and YAP1 activator treated samples. Scale bar, 50µm. (O) Mean fraction of AGR2 and FOXA1 positive cells +/− SEM in Control and YAP1 activator. N = 3 mice per condition, mean and SEM are drawn across 10 crypts per condition for FOXA1, and 14-17 crypts per condition for AGR2. Statistical significance is indicated with *** (p < 0.001).

The integration of inhibitory lateral signalling likely involves Notch activation and the downstream DLL1-repressor HES1 (*63*). Predicted HES1 binding sites at the FOXA1 locus show reduced accessibility in secretory progenitor cells compared to non-FOXA1 expressing cells (e.g. enterocyte progenitors), consistent with the hypothesis that HES1 directly represses FOXA1 (Fig. 4A, fig. S4A-C). The integration of YAP1 activity and lateral signal input creates a network architecture where FOXA1 and DLL1 levels within the same cell are tightly correlated, as evidenced by the minimal overlap in FOXA1 distributions between DLL1^+^ and DLL1^−^ cells (Fig. 4B). These predictions were confirmed experimentally, where FOXA1 and DLL1 levels showed strong positive correlations at the single-cell level (Fig. 4C, 3S). The model also predicts distinct FOXA1 expressions under YAP1 perturbation. Specifically, FOXA1 levels are predicted to be low under YAP1 inhibition and intermediate under YAP1 activation (Fig. 4D,E). Due to mutual inhibitory neighbouring signals under YAP1 activation, FOXA1 is not expected to reach high levels, a result that is consistent with experimental observations (Fig. 4F,G). To test whether neighbouring DLL1 feeds back onto FOXA1, we considered a condition of perturbed Notch signalling (DAPT). In the absence of lateral signalling, both the fraction of DLL1^+^ cells and FOXA1^+^ cells are predicted to strongly increase (Fig. 4D,E), which is indeed confirmed by experiments (Fig. 4F,G). These findings validate the regulatory network in which YAP1 activates FOXA1, while lateral signals from DLL1^+^ neighbours inhibit FOXA1 production. Overall, the model elucidates a landscape of DLL1 states as a function of YAP1 activity and lateral signalling at the single-cell level (Fig. 2H). The architecture of this landscape predicts non-trivial outcomes under combined perturbations of YAP1 and lateral inhibition signalling.

First, while Notch perturbation alone (DAPT) increases DLL1 production (as expected, Fig. 2N,P) (*57*), the model predicts that simultaneous inhibition of YAP1 prevents DLL1 patterning. This occurs because, in YAP1^low^ cells, the DLL1^−^ state becomes stable regardless of the lateral signal received (Fig. 2I, Fig. 4D,E). Remarkably, this prediction was experimentally validated resulting in a complete loss of DLL1^+^ cell emergence (Fig. 4F,G), confirming the essential role of YAP1 pre-patterning Delta-Notch lateral inhibition.

Second, although YAP1 activation disrupts DLL1 patterning by driving cells into an intermediate DLL1 state, in which cells collectively block each other from progressing to the secretory fate, the model further predicts that this effect can be counteracted by simultaneous Notch inhibition (Fig. 4D,E). Under these conditions, the loss of mutual inhibition allows cells to specify into DLL1^+^ fate, resulting in increased fractions of both FOXA1^+^ and DLL1^+^ cells. Indeed, this prediction was also confirmed experimentally: in tissues with homogeneously active YAP1, cells were able to specify into the DLL1^+^ state when Notch signalling was inhibited (i.e. absence of lateral signal) (Fig. 4F,G). These findings further corroborate our model whereby YAP1 pre-patterns the secretory fate at the single-cell level through increased chromatin accessibility. On the tissue scale, YAP1 heterogeneity and lateral signalling are decoded by FOXA1, driving bistability and establishing robust Delta-Notch patterning. This integration of intrinsic YAP1-driven chromatin accessibility changes with extrinsic cell-cell communication ensures patterning during intestinal regeneration.

### Patterning memory from bistability during *in vivo* regeneration

Having constrained our model using *in vitro* organoid data, we extended its application to predict the dynamics of *in vivo* intestinal regeneration. During *in vivo* regeneration, YAP1 activity exhibits a clear temporal pattern: it is strongly upregulated at 24h, followed by density-dependent heterogeneous downregulation at 48h, and is inactive in homeostasis (Fig. 1A,D,I). Using a similar temporal trajectory of YAP1 activity as input for our model (see SI Theory section 6 for more details), we simulated the dynamics of FOXA1 and DLL1 on a 2D lattice. The model predicts that FOXA1 and DLL1 levels are tightly correlated, with FOXA1^+^ and DLL1^+^ cells emerging at around 48h. These cells remain stably patterned even after YAP1 becomes inactive, ensuring tissue organization during homeostasis (Fig. 4H,I). Interestingly, these predictions align closely with experiments: FOXA1 is strongly upregulated 48h after the injury, concurrent with the emergence of secretory progenitors (Fig. 4J,K). Moreover, a direct consequence of the bistability via FOXA1 in our model is that cells can “remember” the influence of prior YAP1 activity. This memory effect enables cells to maintain their DLL1^+^ state as they transition from a YAP1^high^ to a YAP1^low^ state (Fig. 2H-J).

To further validate the bistability model and exclude alternative mechanisms where cells might become insensitive to YAP1 via complex biochemical processes, we tested the effect of YAP1 activation during homeostasis. Consistent with the prediction of our model (Fig. 4L,M), homogeneous YAP1 activation in *in vivo* homeostasis (YAP1 activator for 2.5 days) (*46*) disrupted patterning, resulting in a loss of secretory progenitor cells (Fig. 4N,O). This confirms that YAP1 remains a critical control regulatory factor, not only during regeneration but also in maintaining tissue homeostasis (Fig. 4N,O). Overall, these findings demonstrate that our heterogeneity-driven lateral inhibition model, constrained entirely by *in vitro* data, accurately predicts the dynamics of secretory progenitor cell patterning during *in vivo* intestinal regeneration and homeostasis. FOXA1-mediated bistability provides a robust mechanism for cells to integrate transient heterogenous YAP1 signals with long-term lineage specification, ensuring stable tissue patterning and functionality across regenerative and homeostatic states.

## Discussion

In this study, we demonstrated that YAP1 heterogeneity critically depends on the tissue density of the regenerating intestinal epithelium, acting as a permissive regulator of tissue patterning once an intermediate density threshold is reached.

At the chromatin level, YAP1 pre-patterns the secretory fate by broadly increasing accessibility first, and, as its activity diminishes, creating a chromatin state that becomes favourable for FOXA1 binding. This permissive chromatin environment allows FOXA1-mediated transcriptional regulation, enabling the integration of cell-intrinsic YAP1 states with neighbourhood derived DLL1 signals. FOXA1 then drives tissue patterning through a feedback loop with DLL1, establishing DLL1 bistability, which allows both high and low DLL1 states to coexist for the same level of inhibitory lateral signals at YAP1^low^ states.

This fate bistability of DLL1 is crucial for establishing patterns from a transient YAP1 heterogeneity. Because FOXA1 acts downstream of YAP1, YAP1 heterogeneity functions as a control parameter for patterning, critical at the time of pattern formation but dispensable during homeostasis (*64–66*). Tissue density and integrity emerge as key regulators in this process, defining a “window of opportunity” during which YAP1 heterogeneity is permissive for transitioning from the regenerative state to stable tissue homeostasis. Once density-dependent YAP1 heterogeneity resolves, the system shifts from transient dynamics to the stable maintenance of patterned states, enabling long-term tissue integrity and homeostasis.

Similar lateral inhibition-based bistabilities have been proposed in other systems, such as *Drosophila* bristle patterning (*18*). In that context, bistability operates at intermediate lateral signals, to interpret large-scale pre-patterns across many cells (*18*). In contrast, our findings reveal that bistability at low lateral signals enables the interpretation of short-scale pre-patterning at the single-cell level. On a temporal scale, bistability allows cells to retain a memory of prior YAP1 activity during the regenerative process, maintaining DLL1 patterning even as YAP1 activity diminishes during homeostasis (*67*, *68*).

This process exemplifies how tissue-level properties, such as tissue density, influence both single-cell states and the strength of cell-cell interactions to drive collective behaviours like pattern emergence. Moreover, our findings highlight how regulated heterogeneity of cellular states within a tissue serves as a critical control parameter for cell differentiation and tissue patterning.

This mechanism may represent a general principle governing tissue regeneration, where transient cellular heterogeneity and chromatin pre-patterning enable tissues to sense integrity and functionality and transition back to homeostasis. Failure to properly resolve YAP1 heterogeneity or misinterpretation of tissue density cues could disrupt this balance, leading to unchecked growth or impaired patterning, potentially contributing to pathological conditions such as fibrosis or cancer.

## Acknowledgements

We thank the Liberali group, M. Verbeek, M. Kahnwald and C. Soneson for fruitful discussions; E. Tagliavini and F. Moos for IT support; J. Lüthi for Fractal support; H. Köhler for FACS sorting; T.-O. Buchholz for help on cell segmentation; B. Treutlein and A. Xavier da Silveira dos Santos for help with the library preparation of the single cell ATAC sequencing; M. Lütolf, M. Nikolaev and O. Mitrofanova for providing material for the microwell setup and initial training; J. Tchorz and V. Orsini for providing intestinal tissue sections of YAP1 activator experiment in homeostasis; C. Lambert and N. Accart for help with sample preparation; Claudia Flandoli for graphic design of schemes. We thank C.P. Heisenberg, M. Lütolf, C. Tsiairis, J. Zaugg, G. D’Angelo, L. Giorgetti and the Liberali group for reading the manuscript. C.S was supported by the European Molecular Biology Organization (Postdoctoral Fellowship ALTF 660-2020) and Human Frontier Science Program (HFSP) LT000746/2021-L. D.B.B. was supported by the NOMIS foundation as a NOMIS fellow, and by the Austrian Academy of Sciences through an APART-MINT Fellowship. This work received funding from ERC to E.H. (grant number 851288) and to P.L. (grant number 758617) and from SNSF (POOP3_157531).

## Author contributions

P.L. conceived and supervised the study, P.L., C.S., and S.B. designed the experiments and interpreted the data, D.B.B. and E.H. developed the mathematical model, C.S. and S.B. performed and analysed the organoid imaging experiments. S.B. and M.B.S. designed the single cell image analysis, developed the python package ‘ez-zarr’. C.B. and N.R. performed the ATAC sequencing experiment. U.K. and G.C. prepared the libraries and sequenced ATAC samples. S.B. and M.B.S. analysed ATAC sequencing experiments. O.E.D. and B.S. performed *in vivo* irradiation experiments. C.S. and S.B. analysed the *in vivo* data. S.B. and V.K. collected samples for ChIP sequencing. J.S. and M.B. performed ChIP sequencing. S.B. and M.B.S. analysed the ChIP dataset. L.C.M. generated and validated the FOXA1 knockout line. S.S. designed and cloned the CTGF reporter and Q.Y. generated the organoid line of the CTGF reporter. C.S. performed light-sheet experiments with CTGF line, S.B. analysed light-sheet data. P.L., E.H., D.B.B., S.B. and C.S. wrote the manuscript.

## Data availability

All sequencing data have been depositeda at the Gene Expression Omnibus (GEO) under accession number GSE283662.

---

**Figure S1:**
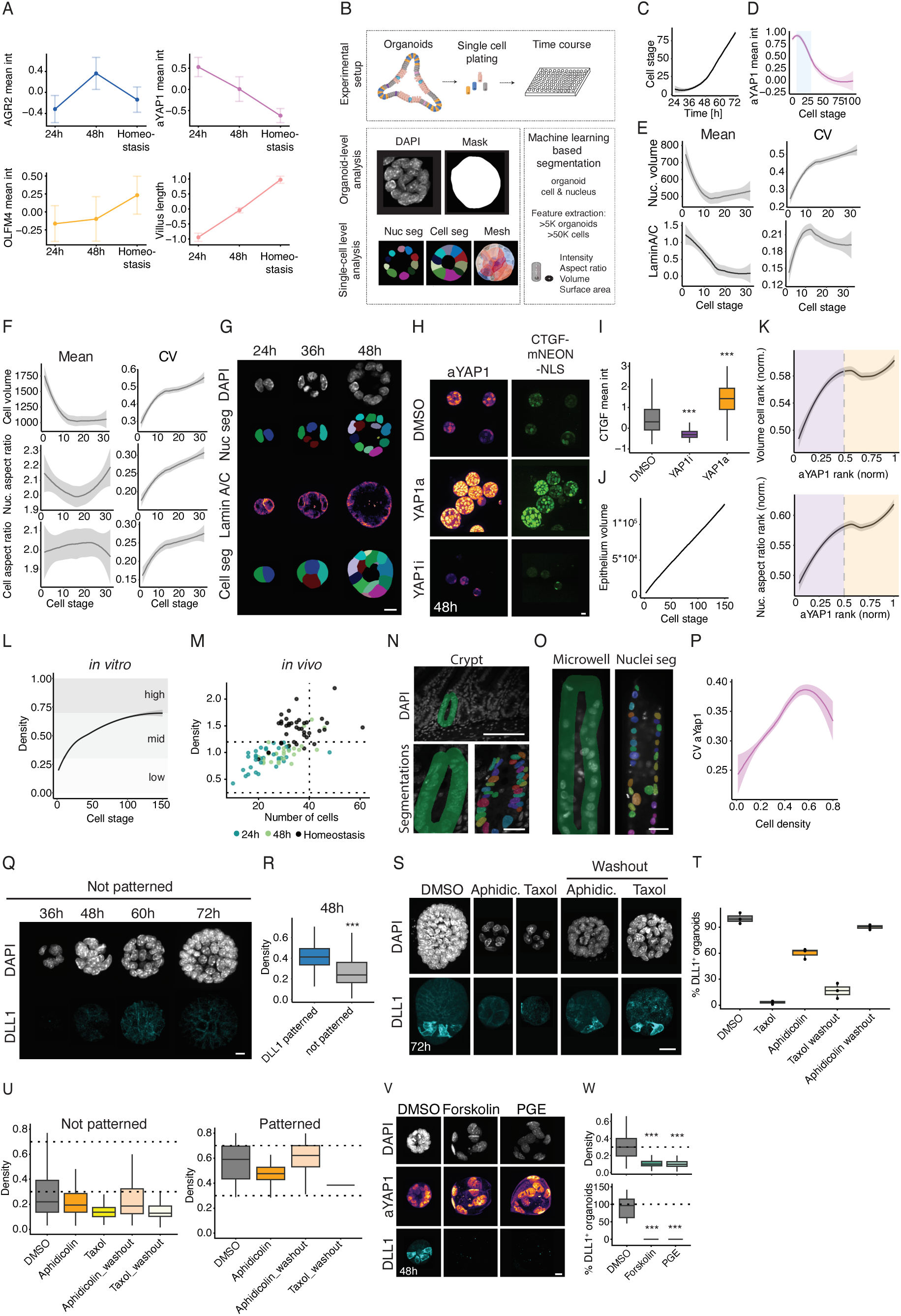
Single-cell and tissue-level feature changes during patterning. (A) Data used to generate smoothened line plots for (Fig.1A): Normalized mean intensities of cell type markers, YAP1 activity and villus length. N = 3 mice, n = 3,118 cells (between 170 and 300 cells per time point per staining). n = 9-20 crypts per timepoint. For villi length each time point counts 25 to 30 villi. Error bars show SEM of independent crypts. (B) Outline of multi-dimensional organoid workflow: 1) Experimental setup shows that organoids are singularised via FACS sorting and plated in Matrigel domes into 96-well plates and fixed at different time points after seeding. 2) Organoid-level analysis entails the use of RDCnet trained organoid segmentation and feature extraction from the region of interest. 3) Single-nucleus and -cell segmentation was performed using RDCnet and Cellpose, nuclei and cell masks were linked, and extraction of intensity and shape features was performed. Mesh of cell segmentation was generated with Paraview. (C) Line plot of cell stage as a function of time in DLL1 patterned organoids. Mean intensities +/− 90% confidence interval. N = 3 experiments, n = 5,210 organoids. (D) Line plot of aYAP1 z-scored mean intensity as a function of cell stage. Blue rectangle indicates DLL1 emergence. N = 3, n = 5,091 organoids. (E, F) Line plot of nuclear volume and LaminA/C z-scored mean intensity (E), cell volume in um^3^, nuclear and cell aspect ratio as a function of cell stage (F), with the mean in the left column and coefficient of variation in the right column. Mean intensities +/− 90% confidence interval. N = 2 experiments, n(mean) = 960, n(CV) = 880 organoids for LaminA/C; N = 3 experiments, n(mean) = 960, n(CV) = 951 organoids for shape features. Scale bar, 10µm. (G) Z-Planes of DAPI, LaminA/C, nuclear and cell segmentation at different timepoints. (H) MIPs of aYAP1, CTGF-mNEON-NLS signal are shown for DMSO control, YAP1 activator and inhibitor at 48h. Scale bar, 10µm. (I) Boxplot of z-scored CTGF mean intensity in YAP1 perturbation conditions from one representative experiment, n(DMSO) = 440, n(YAP1i) = 213, n(YAP1a) = 347 organoids. (J) Line plot of epithelium volume in um^3^ as a function of cell stage. Mean intensities +/− 90% confidence interval. Plot from one representative experiment, n = 2,130 organoids. (K) Organoids with cell stages from 8-20 were individually ranked for cell volume or nuclear aspect ratio against YAP1 activity and line plot shows correlation of intra-organoid single-cell features and aYAP1 ranks. Mean intensities +/− 90% confidence interval. N = 3 experiments, n = 3,731 organoids. (L) Line plot of tissue density, measured as number of nuclei per 100um^2^, as a function of cell stage from 24h-72h organoids. Gray boxes indicate the density bins: low for values up to 0.3, intermediate for values from 0.3-0.7, high from values above 0.7 density. Mean intensities +/− 90% confidence interval. N = 3 experiments, n = 5,065 organoids. (M) Scatter plot of density as a function of number of cells in *in vivo* crypts. Dots are colour coded by days. N = 3 mice per time point. n(24h) = 42, n(48h) = 28, n(homeostasis) = 39 crypts. (N) Z-plane of *in vivo* tissue section labelled with DAPI; green mask indicates manually segmented crypt. Scale bar, 100µm. Lower column: crypt and single-nuclei segmentation. Scale bar, 20µm. (O) Z-plane of DAPI, nuclei signal in microwells (left) and nuclear segmentation (right). Scale bar, 20µm. (P) Line plot of CV of aYAP1 as a function of cell density. Mean intensities +/− 90% confidence interval. N = 3 experiments, n = 5,065 organoids. (Q) MIPs of DAPI and DLL1 in not patterned organoids (no DLL1^+^ cell) at different timepoints of regeneration. Scale bar, 10µm. (R) Boxplot of tissue density (number of nuclei per 100µm^2^) for DLL1 patterned and not patterned organoids at 48h. N = 3 experiments, n = 1,067organoids. (S) MIPs of DAPI and DLL1 of DMSO control, Aphidicolin (0.25µM) and Taxol (5nM) treated organoids from 24h until 72h. Washout conditions of Aphidicolin and Taxol are shown on the right, with treatment window from 24h-48h and washout from 48h-72h. Scale bar, 20µm. (T) Boxplots of percentage of DLL1^+^ organoids (organoids with at least one DLL1^+^ cell) in conditions mentioned in (S). Data from one representative experiment. n = 3 wells per condition. (U) Boxplot of tissue density (number of nuclei per 100µm^2^) for not patterned and DLL1 patterned organoids at 72h. Line at 0.3 indicates transition from low density to intermediate density bin and line at 0.7 indicates transition from intermediate to high density bin. N, n same as in (T). (V) MIPs of DAPI and aYAP1 in DMSO control, Forskolin [10µM], PGE [0.5µM] at 48h. Scale bar, 10µm. (W) Boxplot of tissue density (number of nuclei per 100µm^2^) for (V). N = 2 experiments, n(DMSO) = 782, n(Forskolin) = 482, n(PGE) = 375 organoids. Statistical significance is indicated with *** (p < 0.001).

**Figure S2:**
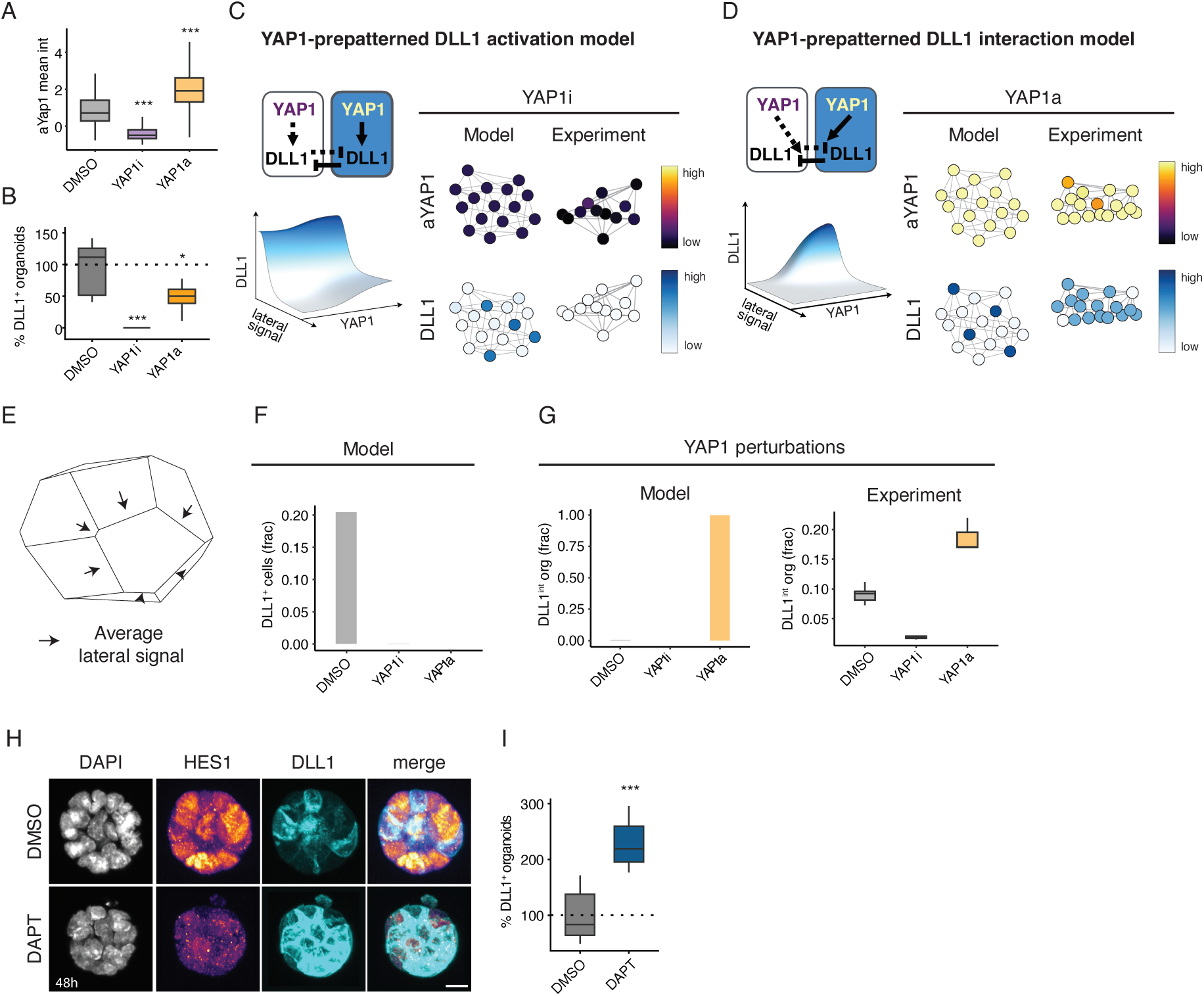
Alternative models for tissue patterning and experimental validation of working model. (A) Boxplot of aYAP1 z-scored mean intensity in YAP1 perturbations at 48h. N = 3 experiments, n(DMSO) = 166, n(YAP1i) = 915, n(YAP1a) = 1,590 organoids. (B) Percentage of DLL1^+^ organoids in DMSO and YAP1 perturbations at 48h. N = 3 experiments, n(DMSO) = 18, n(YAP1i) = 9, n(YAP1a) = 7 wells. (C) Schematic of the YAP1-prepatterned DLL1 activation model. 3D plot of DLL1 production as a function of lateral signal and YAP1 activity (left bottom). Model predictions are compared to the experiment for the YAP1 inhibition, where experimental network plots are generated through projection of spherical coordinates of the 3D cell nucleus position onto a 2D plane, with colours indicating YAP1 activity (top) and DLL1 classification (bottom). Experimental network plots are the same ones as shown in Fig. 2. (D) Schematic of the YAP1-prepatterned DLL1 interaction model, with corresponding DLL1 attractor surface. Model predictions are compared to the experiment for the YAP1 activation. Experimental network plots are the same ones as shown in Figure 2. (E) Scheme of lateral signal that one cell receives. For more detailed information, see SI Theory section 1. (F) Bar plot of average DLL1^+^ cell fraction in DMSO and YAP1 perturbation experiments for the YAP1-prepatterned DLL1-bistabiltiy model (Fig. 2H,Q). (G) Fraction of DLL1^int^ (DLL1 intermediate) organoids, defined as organoids that are enriched in DLL1 intermediate cell fraction (organoids that have a fraction of more than 0.15 of DLL1 intermediate cells) for model (bar plot) and experiment (boxplot). N, n same as (fig. S1A). (H) MIPs of DAPI, HES1 (Notch target), DLL1 and a merge of HES1 and DLL1 of DMSO and DAPT treated organoids and fixed at 48h. Scale bar, 10µm. (I) Bar plot of percentage of DLL1^+^ organoids in DMSO control compared to DAPT at 48h. N = 3 experiments, n(DMSO) = 351, n(DAPT) = 838 organoids. Statistical significance is indicated with * (p < 0.05), *** (p < 0.001).

**Figure S3:**
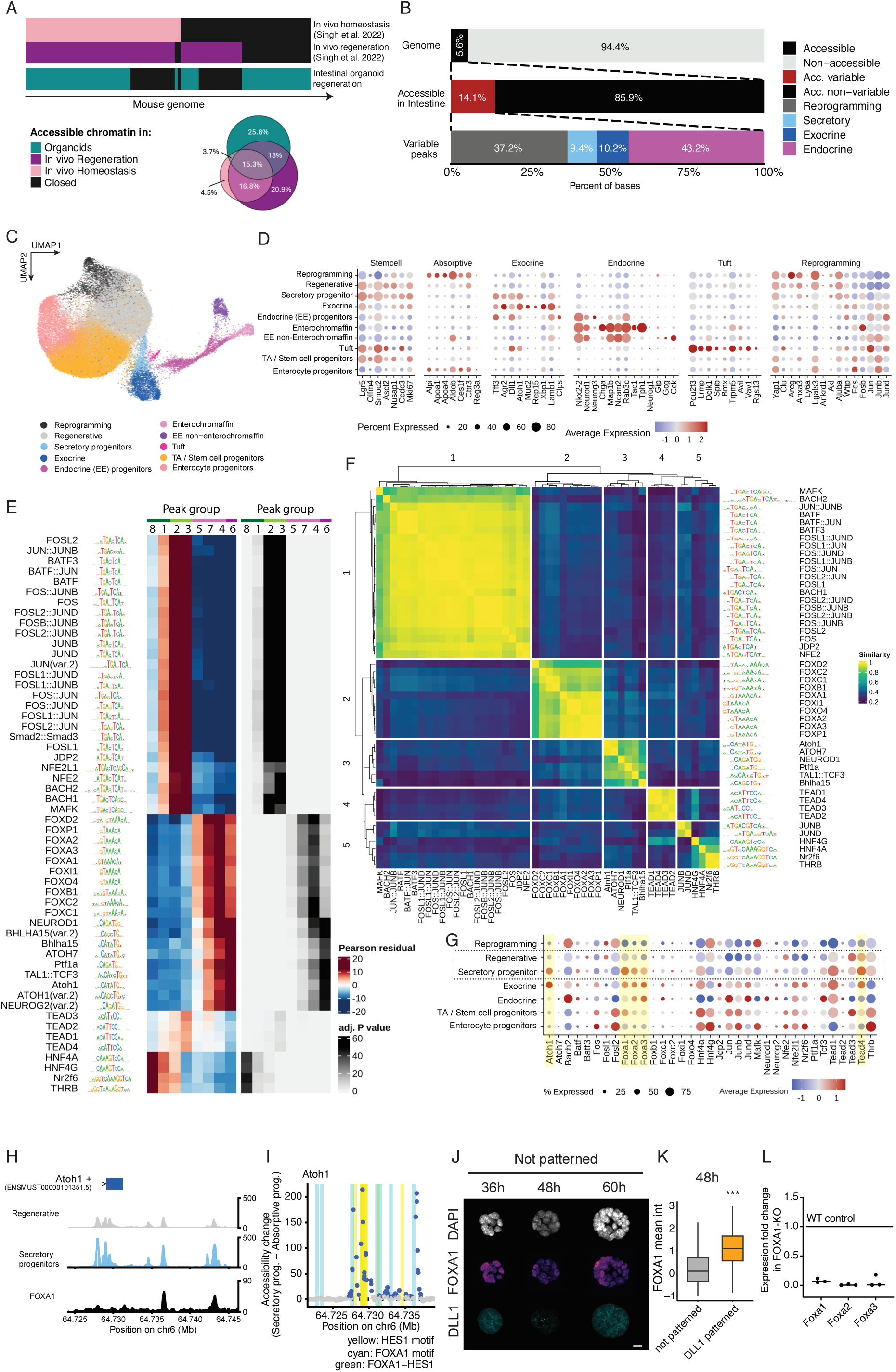
Characterisation of chromatin accessibility changes during patterning and experimental validation of FOXA1 candidate. (A) Bar plots measuring base by base overlapping of genome accessibility in intestinal regeneration *in vivo* (pink and dark magenta) and in organoids (dark cyan). Percentage of overlaps are summarised in Venn diagrams. (B) Bar plots of genome accessibility in mouse intestinal organoid regeneration, showing percentage of total accessible chromatin (5%, top), and variable regions (14%, middle). Bottom row shows how the variable peaks are distributed according to lineages. (C) UMAP embedding coloured by full cell type annotation, including endocrine sub-types. (D) Dot plot of predicted expression for known intestinal markers. (E) Heatmap of motif enrichment analysis in peak groups of interest. (F) Heatmap of similarity in transcription factor motifs. (G) Dot plot of predicted expression across cell types for identified transcription factors. (H) Representative ATAC and ChIP sequencing signal in regenerative (grey) and secretory progenitor cells (blue) around Atoh1 region. (I) Accessibility changes at Atoh1 locus in secretory progenitors (blue). HES1 motif in yellow, FOXA1 motif in cyan, green indicates the overlap of both. (J) MIPs of DAPI, FOXA1, DLL1 of not patterned organoids (no detectable DLL1 signal) at different time points. Scale bar, 10µm. (K) Boxplot of FOXA1 organoid-level z-scored mean intensity at 48h. N = 3 experiments, n(Dll1 patterned) = 1,269, n(not patterned) = 3,618 organoids. (L) Plot shows the expression fold-change of Foxa1, Foxa2, Foxa3 genes in FOXA1-KO for three different monoclonal organoid lines (single dots) normalised to the expression level of housekeeping gene (Elongation factor 1 alpha). Fold-change of 1 indicates WT control and values below indicate decreased expression levels.

**Figure S4:**
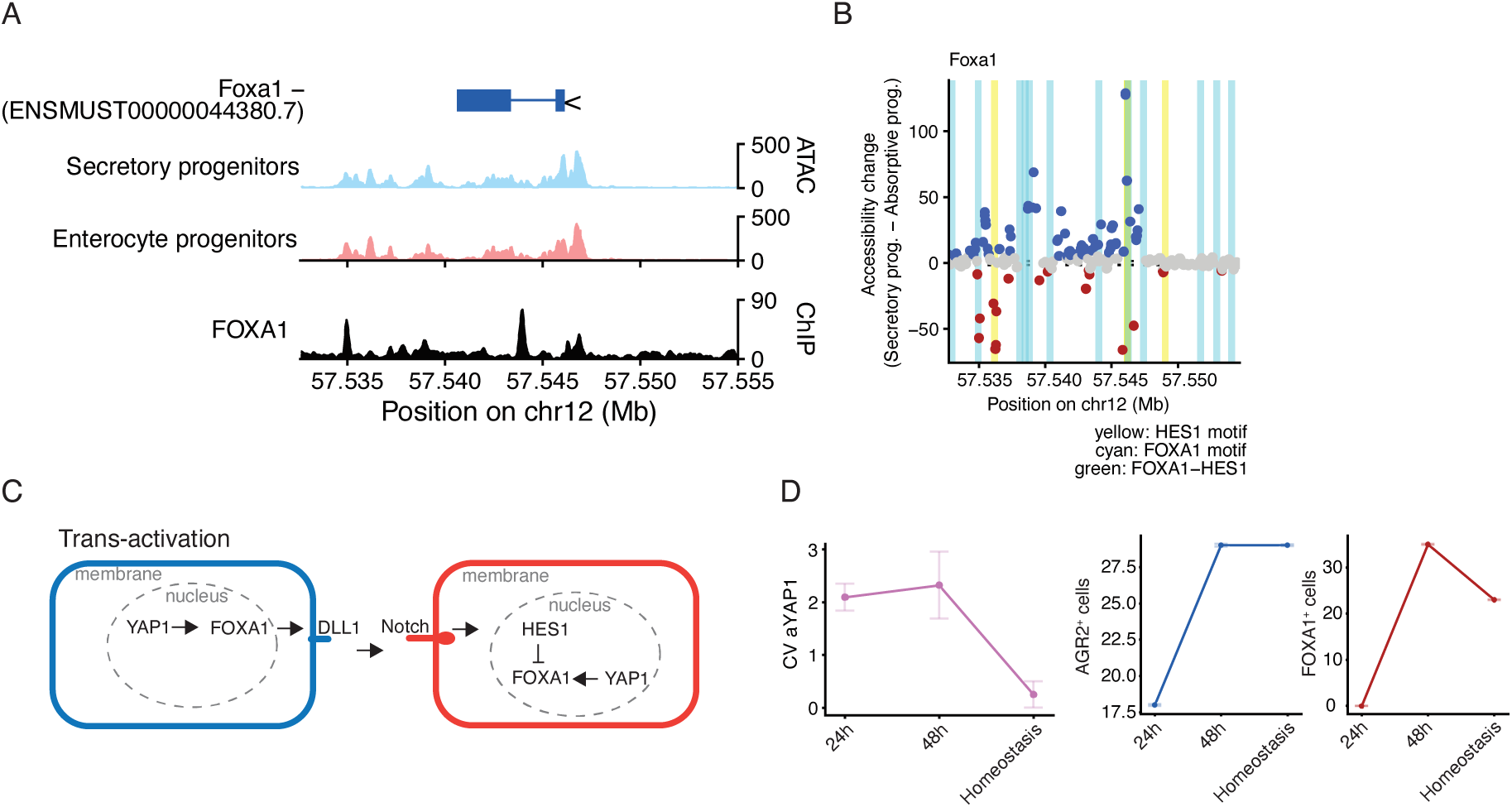
FOXA1-mediated integration of YAP1 heterogeneity and lateral inhibition. (A) Representative ATAC and ChIP sequencing signal in secretory (blue) and absorptive progenitor cells (red) around Foxa1 region. (B) Accessibility changes at Foxa1 locus in secretory progenitors (blue) or absorptive progenitors (red). HES1 motif in yellow, FOXA1 motif in cyan, green indicates the overlap of both. (C) Molecular model of lateral inhibition and feedback on FOXA1. (D) Data used to generate smoothened line plots for Fig. 4K. N, n same as in Fig. 4K.

## SI – Theory: Development of a mathematical model of heterogeneity-driven intestinal organoid patterning

### 1. Model setup

To determine how YAP1 heterogeneity and DLL1-mediated lateral inhibition regulate tissue patterning, we developed a minimal model of intestinal organoid patterning. To this end, we focused on signaling at the 16-cell stage during which pattering is initiated. Based on experimental findings that both YAP1 and DLL1 are critical for patterning, our goal was to identify the minimal regulatory network that links them. Such a network can be written as a general dynamical system for the change of DLL1 concentration 𝐷_𝑖_ in each cell *i*:

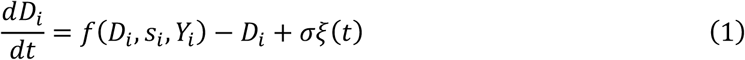

where 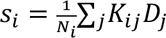 is the lateral inhibition signal sent by neighboring cells, 𝑁_𝑖_ is the number of nearest neighbors of cell *i*, and 𝐾_𝑖j_is the adjacency matrix determining the connectivity of the tissue. This implementation builds on previous work on modeling lateral inhibition signaling^1–3^ in which DLL1-Notch interactions are implemented through a phenomenological neighborhood signal (see section on the model including Notch below). The second term in Eq. (1) models the degradation dynamics of DLL1. Throughout, we used non-dimensionalised units such that the degradation constant of DLL1 is unity, to set the time unit. For more details, see the parameter overview (section 8). 𝜉(𝑡) is a Gaussian white noise process modelling the stochastic nature of cell signaling, with 〈𝜉(𝑡)〉 = 0 and 〈𝜉(𝑡)𝜉(𝑡′)〉 = 𝛿(𝑡 − 𝑡^′^) and amplitude 𝜎. Our goal was to identify and characterize the dynamical system 𝑓(𝐷_𝑖_, 𝑠_𝑖_, 𝑌_𝑖_) that captures the experimental observations, and to make new predictions for perturbation phenotypes and quantitative correlations in the experiments.

To simulate the model, we specified two inputs based on experimental data: the cell connectivity matrix 𝐾_𝑖j_and the YAP1 heterogeneity distribution 𝑌_𝑖_. Our goal was to predict the resulting pattern of DLL1 expression in the tissue.

Firstly, 𝐾_𝑖j_is determined by the spherical connectivity of the tissue. To this end, we placed points (cell nuclei) equistantly on a sphere, and added a random noise with amplitude *A*. We then performed spherical Voronoi tessellation using *scipy*^4^ to identify the neighborhood matrix. This gave access to the distribution of the number of nearest neighbors for each cell. To match the statistics to our experimental data, we similarly performed spherical Voronoi tessellation on 3D segmentation of the cell nucleus positions of intestinal organoids. We fitted the noise amplitude *A* for these distributions to match quantitatively (Fig. S1). This provided a procedure to generate organoid neighborhood maps with the same statistics as those in experiment.

Secondly, we took YAP1 levels to be heterogeneously distributed among different cells. Based on our experimental finding that downstream YAP1 targets exhibit dynamics that are slow compared to the growth of the organoid (Fig. 1E,F – main text), we first took YAP1 levels to be constant in time, to better understand their influence on DLL1 patterning in the model, before implementing slow (although still heterogeneous) decay in YAP1 activity. Based on observations that YAP1 activity is uncorrelated among neighboring cells, we then randomly sampled YAP1 activities 𝑌_𝑖_ for each cell *i* from the experimentally measured YAP1 distribution (Fig. S1). To make model parameters interpretable, we used normalized YAP1 values throughout, meaning we used the normalised experimental YAP1 distribution such that YAP1 activities are scored between 0 (min activity) and 1 (max activity).

Together with the initial condition of no DLL1 expression at the beginning of the patterning process, 𝐷_𝑖_(𝑡 = 0) = 0, this fully specified the setup for the simulations. We then used this model to predict the expected DLL1 pattern for different choices of the dynamical system *f*. This allowed us to systematically constrain *f* and thereby identify the regulatory network coupling YAP1 heterogeneity, DLL1 production and lateral inhibition signaling.

**Figure S1.**
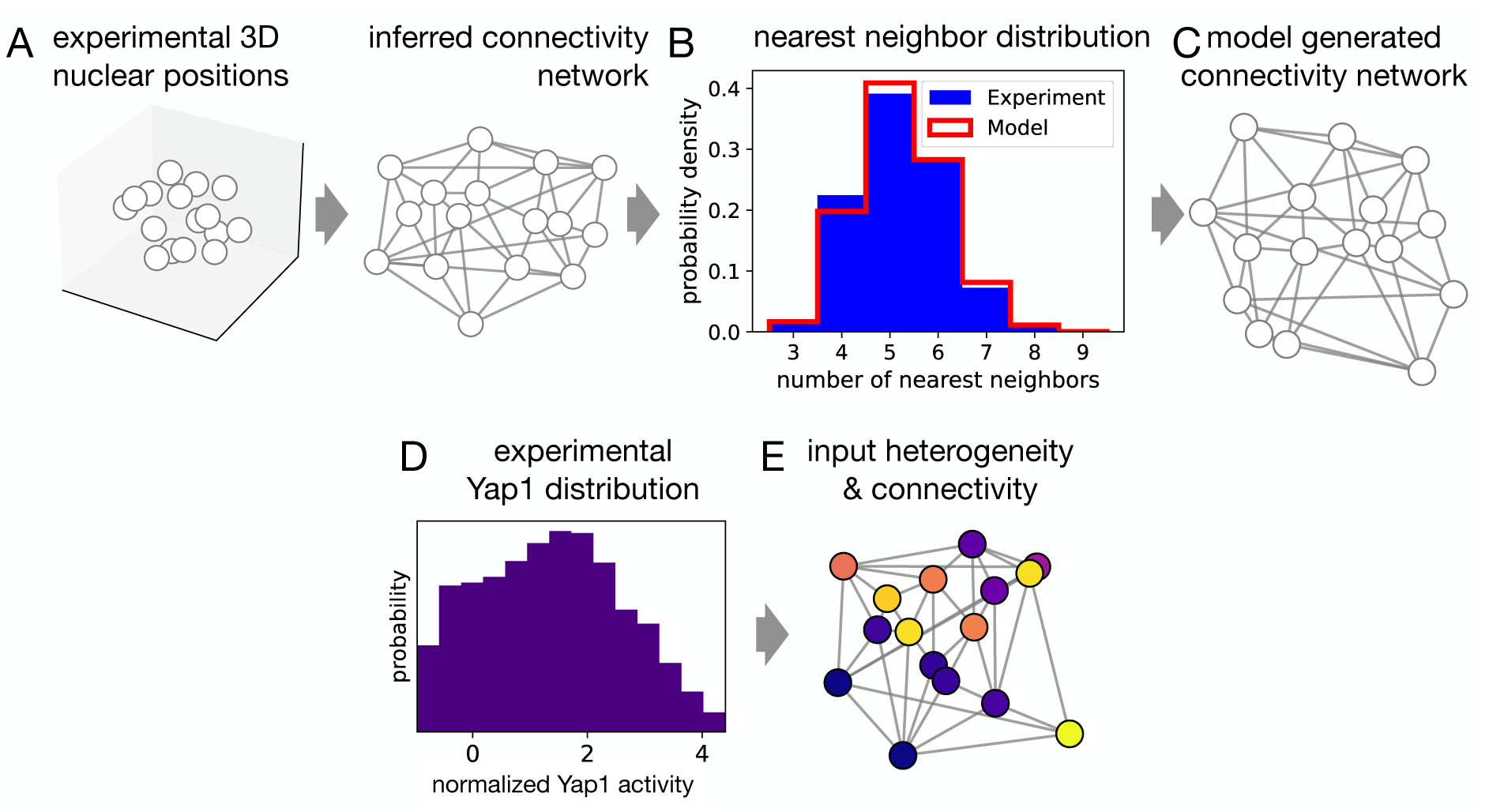
**(A)** Experimental connectivity network is inferred using a spherical Voronoi tessellation of experimental 3D nuclear positions. **(B)** Distribution of the number of nearest neighbors per cell in experimental (blue) and model generated (red) connectivity networks. **(C)** Model generated connectivity network with nearest neighbor distribution matching the experiment. **(D)** Normalized experimental distribution of YAP1 activities in wildtype organoids. To determine the YAP1 distribution during patterning, only patterned organoids (i.e. at least one DLL1+ cell) are included, with cell numbers between 8-20 cells. **(E)** Model generated connectivity network with YAP1 heterogeneity generated by random sampling from the experimental distribution in (D).

### 2. Ruling out pure lateral inhibition signaling

Using our model, we first simulated the expected dynamics of pure lateral inhibition, i.e. no coupling to YAP1 heterogeneity. In this case, the dynamical system is given by^1^

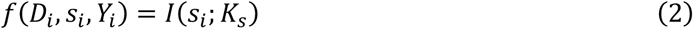

where the first term is a production term of DLL1 that is inhibited by the signal 𝑠_𝑖_ and the second term is a degradation term. Throughout, we used activation and inhibition functions modelled as Hill functions; which have been shown in multiple works to reproduce well the key aspects of lateral inhibition^1,3^, i.e.

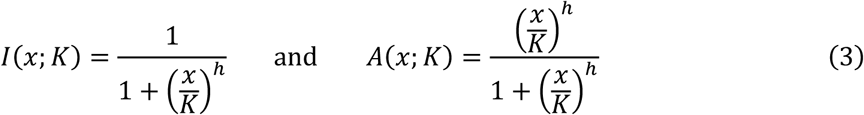

where *h* are the Hill exponents setting the steepness of the curve, and *K* is the Hill threshold parameter setting the location of the steepest response (at *x*=*K*).

Thus, for pure lateral inhibition we could visualize the DLL1 production term as a function of the signal received by each cell, which is a monotonically decreasing curve as a function of signal (Fig. S2A). This already demonstrates that, by construction, the DLL1 production of each cell only depends on the received lateral inhibition signal, and would be independent of YAP1 heterogeneity, which is in contrast to the experimental observation that DLL1 patterning is critically affected by YAP1 perturbations. Furthermore, the pure lateral inhibition model predicts an average of 4 DLL1+ cells per 16-cell organoid, i.e. a DLL1+ fraction of 0.25 (Fig. S2B, C). This prediction is independent of parameters, as it only depends on the connectivity of the tissue, which we have constrained experimentally. It contrasts with the expected fraction of 1/3 ≈ 0.33 DLL1+ cells in a 2D hexagonal lattice. This is because in a 2D hexagonal system, all cells have 6 nearest neighbours, which is in contrast to the nearest neighbor distribution in our spherical organoids with a high fraction of cells with 5 nearest neighbors (Fig. S1B). However, this prediction exceeds the experimentally measured fraction of 0.2 ± 0.02 (SEM). Taken together, these results suggest that a pure lateral inhibition model is unable to capture the experiments.

**Figure S2.**
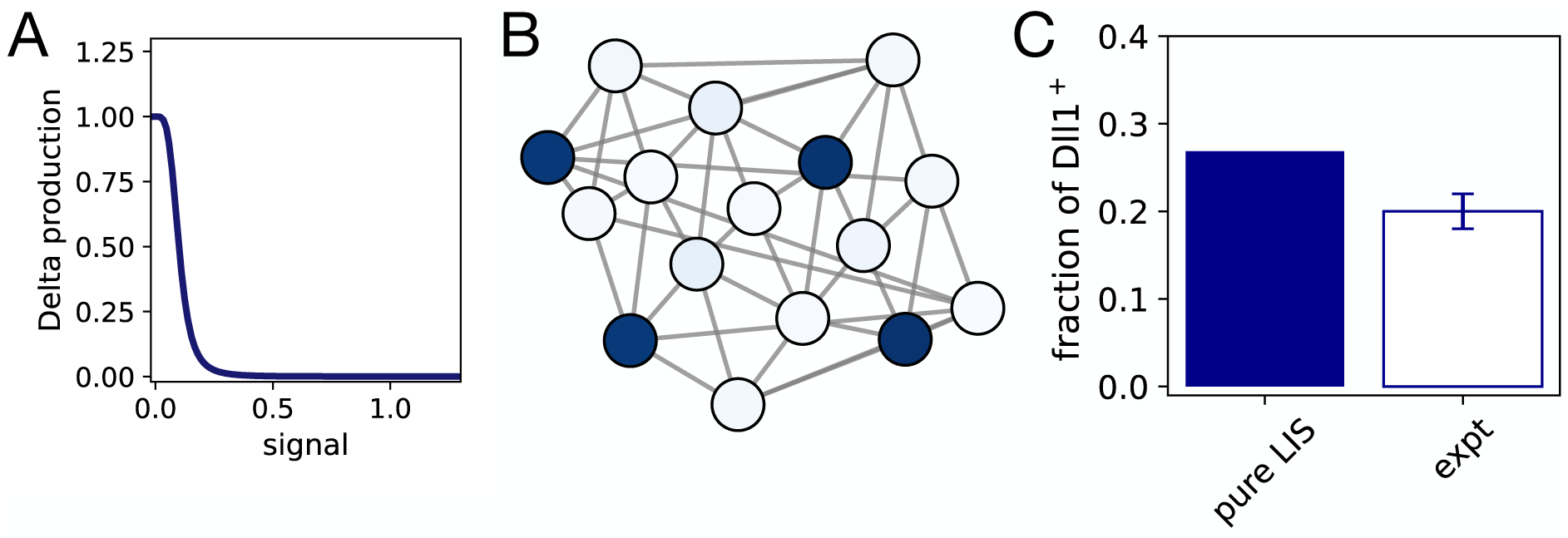
**(A)** DLL1 production as a function of lateral signal for this model. **(B)** DLL1 intensity on model generated network. **(C)** Pure lateral inhibition signaling predicts DLL1+ cell fraction of 0.25, compared to 0.2 ± 0.02 (SEM) observed experimentally.

### 3. Identifying a regulatory network constrained by perturbation phenotypes

We next systematically screened possible model candidates coupling YAP1 heterogeneity and lateral inhibition signaling. Our goal was to identify a model that is able to capture the following experimental observations: (1) The proportion of DLL1+ cells matches that in experiment, i.e. 0.2 ± 0.02. (2) Yap+ cells are more likely to be DLL1+. (3) There are no DLL1+ cells for homogeneously low YAP1 (YAP1 inhibition). (4) There are no DLL1+ cells for homogeneously high YAP1 (YAP1 activation).

To this end, we constructed proposed dynamical systems 𝑓(𝐷_𝑖_, 𝑠_𝑖_, 𝑌_𝑖_) by combining activation and inhition functions (Eq. 3) of the key variables 𝐷_𝑖_, 𝑠_𝑖_, 𝑌_𝑖_. We systematically increased the complexity of the model by first considering combinations of two regulatory functions, before considering combinations of three functions. Rather than explicitly going through all possible combinations, we here present the predicted dynamics of the most promising model candidates, specifically those that combine the functions 𝐼(𝑠_𝑖_) to model lateral inhibition and 𝐴(𝑌_𝑖_), motivated by the observed positive correlation between DLL1 and Yap. Each of these models can be characterised using the curve of DLL1 production as a function of the lateral signal 𝑠_𝑖_, plotted for different values of YAP1 𝑌_𝑖_. Thus, in these models, each cell has a different DLL1 vs. lateral signal curve which is determined by its YAP1 level. For each model, we performed computational parameter screens of the key parameters that determine these curves (see Fig. S4).

We first considered multiple possible implementations of a scenario in which YAP1 positively regulates DLL1 (which has been proposed in several other experimental scenarios, see refs.^5,6^). To this end, we combine the functions 𝐼(𝑠_𝑖_) and 𝐴(𝑌_𝑖_) in different ways. We first consider the two possible combinations of just these two functions, in which YAP1 regulates DLL1 either cell-autonomously, or through interactions.

#### Ruling out simple models: pre-patterned DLL1 production or interactions

First, we considered a model in which YAP1 regulates a cell-autonomous production of DLL1:

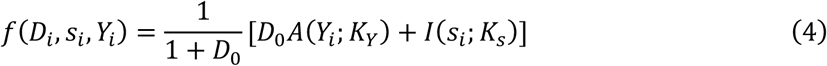

We termed this model the **Yap-prepatterned DLL1 production model**, since in this model, YAP1 generates a baseline activation of DLL1, on top of which additional DLL1 production can occur depending on the lateral signal. The additional factor of 1/(1 + 𝐷_0_) is simply for normalisation of DLL1 levels to a maximum of 1, which does not affect the results. We visualised the steady-state DLL1 levels of the model as a function of the YAP1 level 𝑌_𝑖_ and the received lateral signal 𝑠_𝑖_, defined by

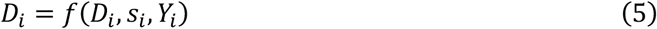

This showed that as a function of increasing Yap, all cells have a higher background-level of DLL1, independent of the lateral signal, but the DLL1+ state (𝐷_𝑖_ = 1) is only reached for cells that receive low lateral signal (Fig. S3A). For parameters 𝐾_𝑠_ < 𝐷_0_, this model captures the YAP1 activator experiment through an interesting effect: because YAP1 prepatterns DLL1, each cell receives sufficient signals from the neighbors to block any further DLL1 production through lateral signal; thus patterning is prevented (Fig. S4). However, this model fails because it predicts patterning in the YAP1 inhibitor experiment. Specifically, while for low Yap, there is no prepatterning (for low Yap, 𝐴(𝑌_𝑖_; 𝐾_F_) = 0), but the second term in Eq. (5) leads to DLL1 patterning through lateral inhibition interactions, which is not perturbed by YAP1 inhibition. Importantly, these findings are independent of parameters: for none of the tested combinations of (𝐾_𝑠_, 𝐾_F_), all three conditions (wildtype, YAP1 inhibition, YAP1 activation) are captured (Fig. S4A). Note that in those regions of parameter space in which both YAP1 inhibition and YAP1 activation are captured, there is also no patterning in the wildtype scenario, which means these combinations are also inconsistent with experiment.

Second, we considered a model in which YAP1 regulates the DLL1 production term that is controlled through lateral inhibition, i.e. it modulates how cells respond to external signal received through interactions with neighboring cells:

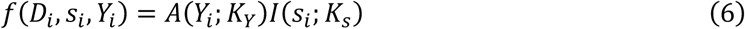

We termed this model the **Yap-prepatterned DLL1 interaction model** (Fig. S3B). Here, only Yap+ cells are able to become DLL1+: for low Yap, 𝐴(𝑌_𝑖_; 𝐾_F_) = 0, and thus no DLL1 production occurs, irrespective of the received lateral signal. This model does not capture the experiment since it predicts a 25% fraction of DLL1+ cells in a YAP1 activation experiment (Fig. S4). This is because for high Yap, 𝐴(𝑌_𝑖_; 𝐾_F_) = 1, and thus the model simply recovers the pure lateral inhibition case (Fig. S2). Again, we find similar behavior across parameter combinations: for none of the tested combinations of (𝐾_𝑠_, 𝐾_F_), all three conditions (wildtype, YAP1 inhibition, YAP1 activation) are captured (Fig. S4B).

#### Model with combined Yap-prepatterned DLL1 production and interactions

We next explored a model that combines these two options:

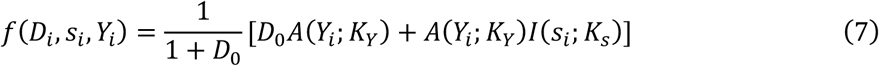

which we termed the **Yap-prepatterned DLL1 production and interaction model** since it combines the previous two models (Fig. S3C). This combination exhibits the two key features that allowed the previous models to capture the perturbations: first, like the Yap-prepatterned DLL1 production model, it exhibits a DLL1 intermediate state at high Yap, leading to blocking of patterning in the YAP1 activation experiment (Fig. S3C). Second, like the Yap-prepatterned DLL1 interaction model, steady-state DLL1 levels vanish for low YAP1 cells independent of the lateral signal, leading to absence of patterning in the YAP1 inhibition experiment (Fig. S3C). Thus, this model successfully predicted the absence of patterning in both YAP1 inhibitor and activator experiments, while capturing the correct proportion of DLL1+ cells in the wildtype (Fig. S4C).

However, this model also predicted that if the YAP1 level of a cell is downregulated, it can no longer be DLL1 positive. This is fundamentally inconsistent with the observed time-progression in the experiment, in which YAP1 is heterogeneously downregulated in time, leading to YAP1 heterogeneity being only transient, with uniformly low YAP1 activity at late times. This leads to patterning during the heterogeneous phase, but DLL1+ cells are lost at later stages when YAP1 activity is low. This led us to hypothesize that in addition to the DLL1-stable state for Yap-low cells, a second DLL1+ state should exist for these cells, separated by a barrier. This suggested bistability as a function of the received lateral signal, which is the simplest mechanism allowing an inductive signal to be first critical and then dispensible^7^. Interestingly, such bistability has been previously proposed in the context of DLL1-Notch signaling in other systems^3^, making it a promising model candidate.

**Figure S3.**
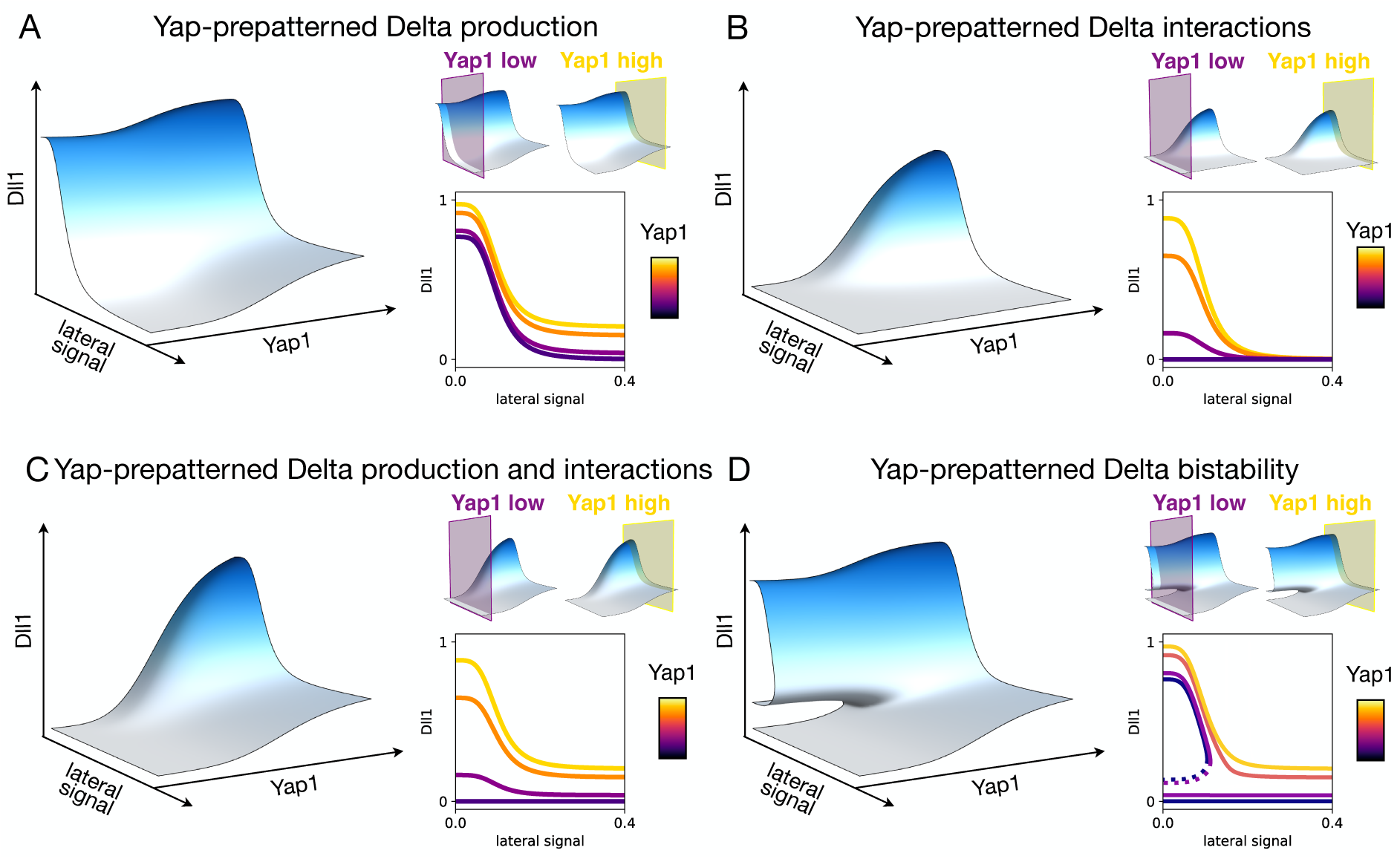
Analysis of stable manifolds for each model candidate. For each model, first the surface of stable DLL1 states as a function of lateral signal and YAP1 is shown. Cuts through the surface at low and high YAP1 are indicated, with the DLL1 vs lateral signal curve for different YAP1 values (𝑌_𝑖_ = 0.1, 0.4, 0.7, 1) shown below.

#### Yap-prepatterned DLL1 bistability model

A model in which YAP1 levels determine a cell-intrinsic bistability of DLL1 is implemented by making the interaction term depend not directly on YAP1 levels 𝑌_𝑖_ as in the Yap-prepatterned DLL1 production and interaction model (Eq. 7), but on DLL1 itself:

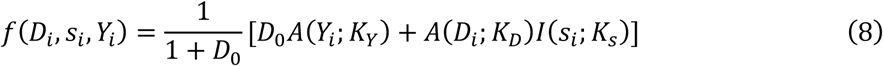

which we termed the **Yap-prepatterned DLL1 bistability model** (Fig. S3D). In contrast to the previous models, in which the signal-response curves were always monostable, low YAP1 cells now exhibit stable states for both DLL1+ and DLL1-, with the preferred state depending on signaling history. This bistability is controlled by the YAP1 activity of the cell, which gives the cellular landscapes the following structure (Fig. S3D): Yap-low cells are bistable at low lateral signal, meaning they can remain in a low-DLL1 state; at high lateral signal, there is only the low DLL1 state available. Yap-high cells have a single stable DLL1-high state at low signal, and an intermediate DLL1 state at high signal due to the prepatterning term. Thus, in the YAP1 inhibitor, cells are stuck in the low DLL1 state. In the YAP1 activator, all cells exhibit intermediate DLL1 and thus are blocked from becoming DLL1+ by the lateral signal of the neighbors. In numerical simulations, this model again is able to predict the YAP1 inhibition and activation experiments, while also capturing the experimentally measured DLL1+ fraction in wildtype (Fig. S4D). Throughout these models, we include a moderate level of intermediate noise in the chemical kinetics equations (𝜎 = 0.02, see parameter table), which is low enough that spontaneous stochastic transitions across the barrier in the bistable regime are rare.

**Figure S4.**
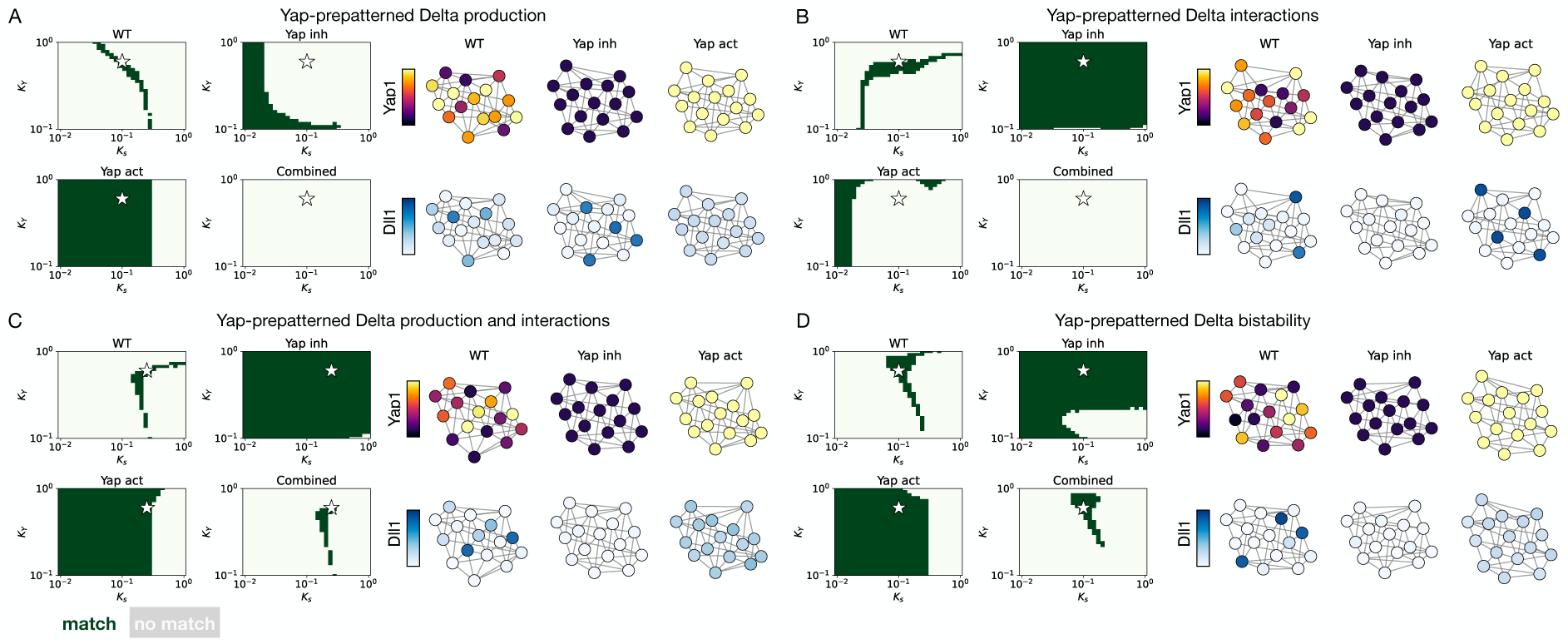
Analysis of parameter spaces of the four model candidates, varying 𝐾_𝑠_ and 𝐾_F_. Each panel shows the region in parameter space (marked in dark green) in which the model captures each of the three conditions: for the wildtype, the condition for match is a DLL1+ fraction of 0.2 ± 0.02; for YAP1 inhibitor and activator a DLL1+ fraction of 0. The combined plot shows the parameter range in which all three conditions are matched. The star indicates the parameter combination used for generating network plots: (A) 𝐾_𝑠_ = 0.1, 𝐾_F_ = 0.6, (B) 𝐾_𝑠_ = 0.1, 𝐾_F_ = 0.6, (C) 𝐾_𝑠_ = 0.25, 𝐾_F_ = 0.6, (D) 𝐾_𝑠_ = 0.1, 𝐾_F_ = 0.6. In all panels, we use a noise amplitude 𝜎 = 0.02 and Hill exponents ℎ =4. In panel D, the additional parameter 𝐾_𝐷_ = 0.2 is used.

### 4. Including FOXA1 in the model

To enable a mechanistic interpretation of the dynamical system identified by our model screen, we considered how the DLL1 cell-intrinsic bistability and the YAP1 prepatterning could be implemented mechanistically through interaction of molecular pathways.

From a theoretical perspective, a minimal implementation of bistability is a positive autoregulation, for instance if DLL1 could activate an intermediate species *X* which in turn positively regulates DLL1 production. However, in the canonical model of Delta-Notch signaling, Delta has no signaling pathway linking back to the nucleus, and therefore does not directly control gene regulation^8^. An alternative (mathematically equivalent) formulation is that the species *X* is trans-inhibited by the DLL1 of the neighboring cell. Mechanistically, this could function through trans-activation of Notch by DLL1 combined with cis-inhibition of *X* by Notch (see next section on an explicit model including Notch dynamics). The intermediate species *X* could furthermore provide a link from YAP1 to DLL1, if YAP1 activates *X* and *X* activates DLL1. As discussed in the main text, based on single-cell ATAC sequencing, we identified the transcription factor FOXA1 as a candidate for this intermediate species (Fig. 3 – main text), and therefore included its dynamics in our model.

A minimal extension of our Yap-prepatterned DLL1 bistability model (Eq. 8) which includes FOXA1 dynamics is given by the following set of coupled equations:

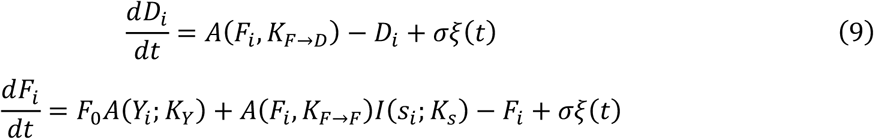

This model is equivalent to the identified dynamical system from the previous section, under the assumption that the dynamics of DLL1 production are fast compared to those of FOXA1: if we approximate the first equation to be in steady-state, i.e. 𝑑𝐷_𝑖_/𝑑𝑡 = 0, and linearize the Hill function, we obtain 𝐷_𝑖_ ≈ 𝐹_𝑖_. Substituting into the second equation recovers the Yap-prepatterned DLL1 bistability model (Eq. 8). Similar to the Yap-DLL1 models, we fitted the parameters of this model to simultaneously predict patterning of FOXA1 and DLL1, given an input distribution of YAP1 (Fig. S1).

### 5. Approximating molecular model with Notch interactions

At a molecular level, the signaling described by our model is implemented by trans-membrane interactions of DLL1 and Notch. Throughout our modeling approach, we neglected the dynamics of Notch expression as we assume these to be strongly anti-correlated to those of DLL1, and therefore used the inhibition by neighborhood signal as an effective implementation of DLL1-Notch dynamics throughout. This also reduced the number of parameters in the model.

To demonstrate how our model (Eq. 9) is implemented at the molecular level, we first wrote a complete model including DLL1-Notch trans-membrane interactions (Fig. S4C – main text). Specifically, we assumed that the inhibition of FOXA1 by DLL1 of neighboring cells is implemented the following way: DLL1 trans-activates Notch in our system (neglecting alternative or additional mechanisms such as cis-inhibition^2^ and cis-activation^9^), Notch activates HES1, and HES1 inhibits FOXA1. Adding these additional steps in our model formulated in Eq. 9 leads to the following set of equations:

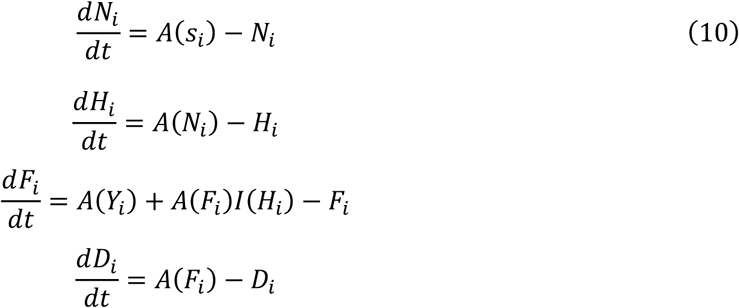

Where 𝑁_𝑖_ is Notch and 𝐻_𝑖_ is HES1 in cell *i*. As before, 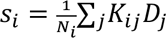 is the DLL1 level of neighboring cells. If Notch and HES1 are in steady-state (i.e., their dynamics evolves faster than those of FOXA1 and DLL1), i.e. 𝑑𝑁_𝑖_/𝑑𝑡 = 𝑑𝐻_𝑖_/𝑑𝑡 = 0, and linearizing the Hill functions, we obtain 𝐻_𝑖_ ≈ 𝑁_𝑖_ ≈ 𝑠_𝑖_, which consequently maps Eq. 10 to Eq. 9. Therefore, this more detailed molecular model including Notch and HES1 directly maps onto our simplified model, in which the neighbor DLL1 signal inhibits FOXA1 (Fig. S4C – main text).

### 6. Prediction of in vivo regeneration dynamics

To make predictions for the expected dynamics of FOXA1 and DLL1 emergence during regeneration *in vivo*, we generalized our model to larger tissues and to a dynamical YAP1 heterogeneity. As before, we will use a distribution of YAP1 states among cells as an input, and predict the expected output, namely FOXA1 and DLL1 patterns.

To this end, we simulated dynamics on a large 2D tissue with a hexagonal lattice of cells. We focus on the dynamics in the bulk of the tissue to avoid boundary effects. We then used a minimal implementation of the dynamics of YAP1 activity observed *in vivo*. Initially, cells are uniformly active in Yap, and we therefore set all cells to uniform maximal Y=1. In homeostasis, YAP1 is uniformly inactive, and the final state is therefore with all cells distributed around uniform Y=0. To interpolate between these two states, we generate YAP1 trajectories in single cells that heterogeneously reduce their YAP1 activity at randomly chosen rates (Fig. S5A). Given that the *in vivo* experiments, and YAP1 downregulation, occur on time scales of days, we use a time-scale for YAP1 dynamics that is much slower than the time-scale of equilibration of the dynamical system, such that the FOXA1 and DLL1 dynamics are in quasi-steady-state. This is therefore consistent with the implementation of YAP1 heterogeneity in the modelling results above. The model then predicts the expected output patterns of FOXA1 and DLL1 (Fig. S5B). To model the *in vivo* YAP1 activation experiment, we furthermore rapidly increase YAP1 in simulations after homeostasis was reached. In our model, such YAP1 activation brings all cells back to the intermediate state, thus destroying the DLL1 spatial pattern. The fact that the experiment is consistent with this prediction is nontrivial as it demonstrates that YAP1 continues to control DLL1 patterning even in homeostatic states, consistent with our model.

**Figure S5.**
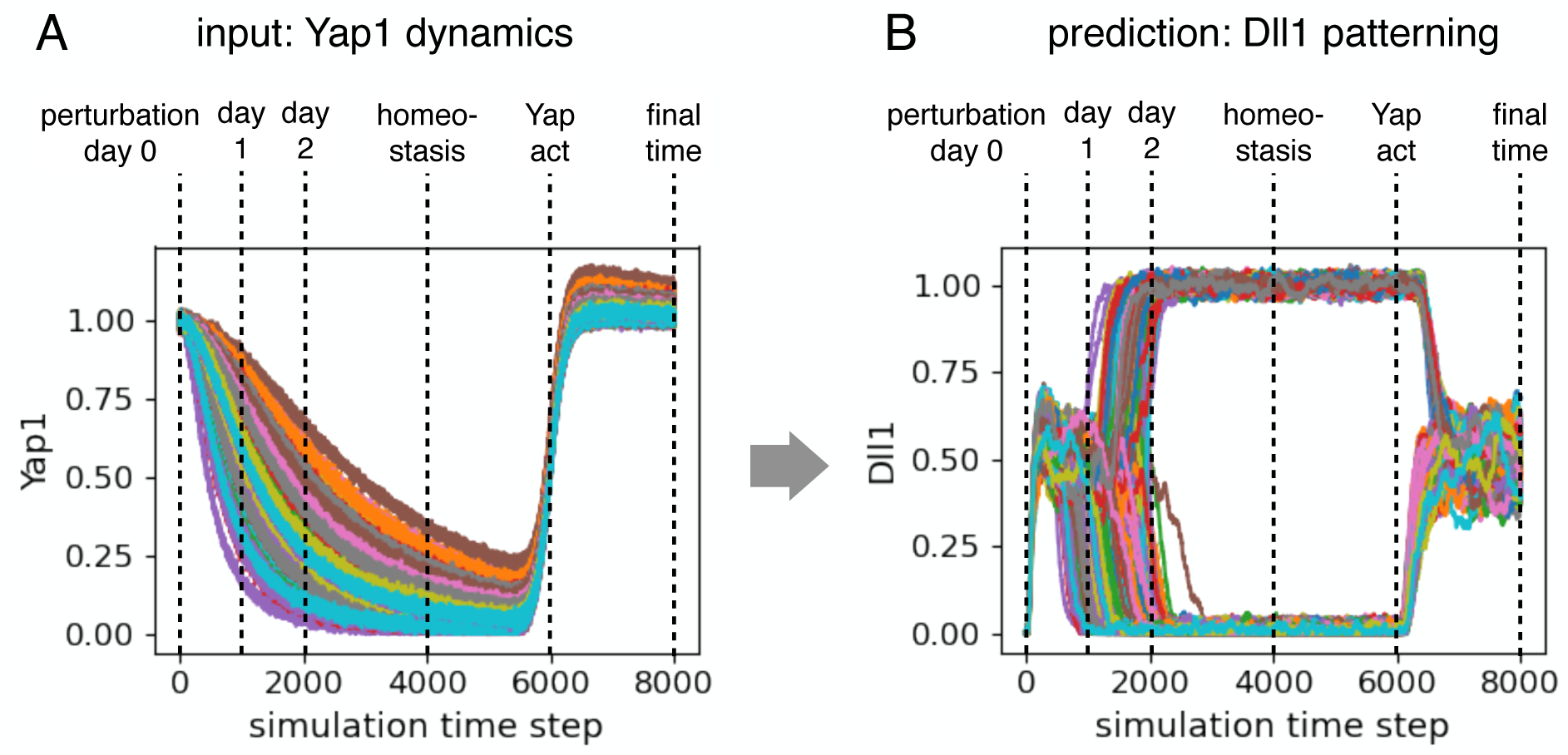
YAP1 and DLL1 dynamics in *in vivo* simulations. **(A)** YAP1 trajectories are generated as input, modeling initial heterogeneous downregulation to homeostasis. Beyond 6000 time-steps, rapid increase of YAP1 models the YAP1 activation experiment, in which homeostatic tissue is perturbed. **(B)** The model predicts the expected output DLL1 pattern as a function of time, as visualised in Fig. 4 in the main text.

### 7. Numerical implementation

The dynamical system (Eq. 1) is numerically integrated using custom-written python code using numpy [9]. The time derivative is approximated using Euler forward differences. Specifically, the following scheme for each cell *i* is used:

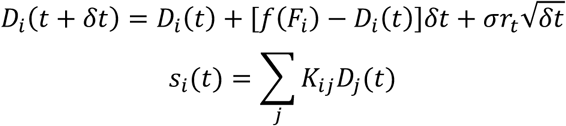

where 𝑟_𝑡_ is a Gaussian random number with mean zero and unit variance.

### 8. Parameter overview

The parameters of the final YAP1-FOXA1-DLL1 model are summarized below.

**Table S1.**
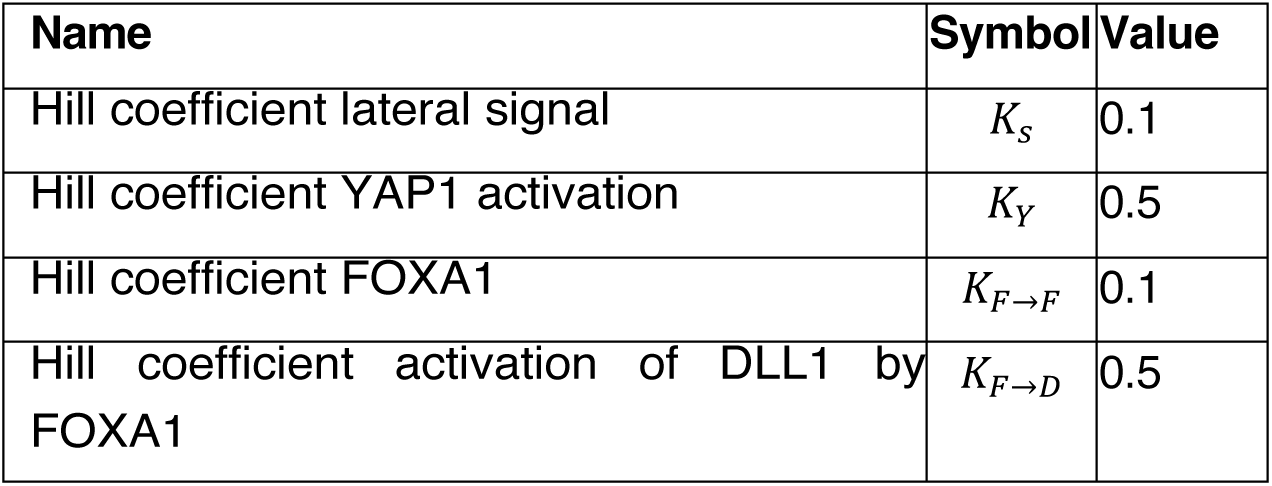
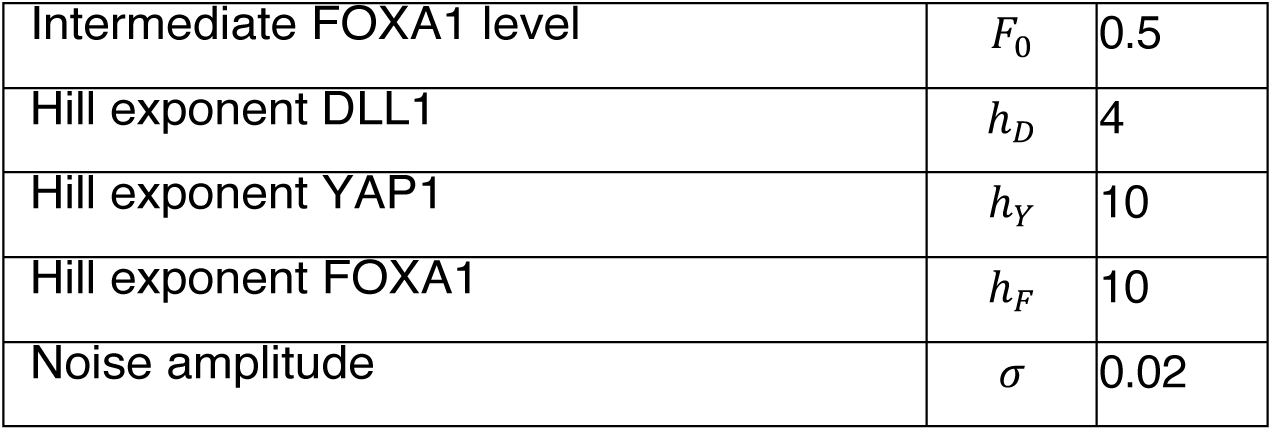
Overview of parameters used, in units with non-dimensionalised time and concentration coordinates.

## Materials and Methods

### Animal experiments and mouse intestinal organoid lines

For irradiation experiments, wild-type C57BL/6J mice were bred and maintained in individually ventilated cages at the Francis Crick Institute Biological Research Facility under specific pathogen-free conditions. Mouse experiments were performed according to the guidelines detailed in the Project Licence granted by the UK Home Office to Brigitta Stockinger (PP0858308). All other animal experiments were approved by the Basel Cantonal Veterinary Authorities and performed in accordance with the Guide for Care and Use of Laboratory Animals. Male outbred mice between 8 and 12 weeks old were used for experiments. Wild-type C57BL/6 mouse line was used (Charles River Laboratories and Jackson Labs).

### Mouse intestinal organoid maintenance culture

Organoids were generated from isolated crypts of mice, as previously described (*1*). In brief, the first part of the small intestine, the Duodenum, was cut open longitudinally, flushed with cold PBS and villi were removed using a cold glass slide. Tissue was cut into fragments and incubated in 2.5 mM EDTA/PBS at 4°C for 30min under shaking. Tissue was transferred into cold PBS and shaken vigorously, and supernatant was collected. Procedure was repeated two more times, to collect in total three fractions. Each fraction was centrifuged at 300xg for 5 min at 4°C. Pellets were collected in 5ml DMEM/F12/Hepes (STEMCELL), 1× Glutamax (Thermo Scientific), and 100μg/ml Penicillin-Streptomycin and centrifuged at 300xg for 5min at 4°C. After removing the supernatant, the pellet was resuspended in IntestiCult Organoid Growth Medium (STEMCELL Technologies) with 100 μg/ml Penicillin-Streptomycin, 10µM ROCK inhibitor. After that maintenance of organoid cultures was performed in IntestiCult Organoid Growth Medium (STEMCELL Technologies) with 100μg/ml Penicillin-Streptomycin.

### Generation of FOXA1-KO organoid line

For the CRISPR-Cas9 mediated knockout of FOXA1, 20bp long guide sequences were designed using the online CHOPCHOP tool (https://chopchop.cbu.uib.no) (table S1) and cloned into the lentiCRISPRv2 backbone (Addgene plasmid #52961) using the protocol described previously (*2*). Briefly, vectors were digested using BsmBI and used for ligation with the phosphorylated sgRNA oligos. Lentiviral particles were produced in HEK293-T LentiX cells (Takara Bio Cat: Z2180N Lot: AN60006S) using a 2nd generation lentiviral system (pMDG.2, Addgene plasmid #12259, and psPAX2, Addgene plasmid #12260) and the plasmid described above. Supernatant from producing cells was concentrated using Amicon Ultra-15, 100 kDa centrifugation columns (Merck Millipore, cat# UFC910024) and used for lentiviral infection as described previously (*3*). Generated organoids were passaged once before selecting with 1µg/ml Puromycin over the course of 2 passages. After selection, organoid culture was performed as described above.

**Table S1.**
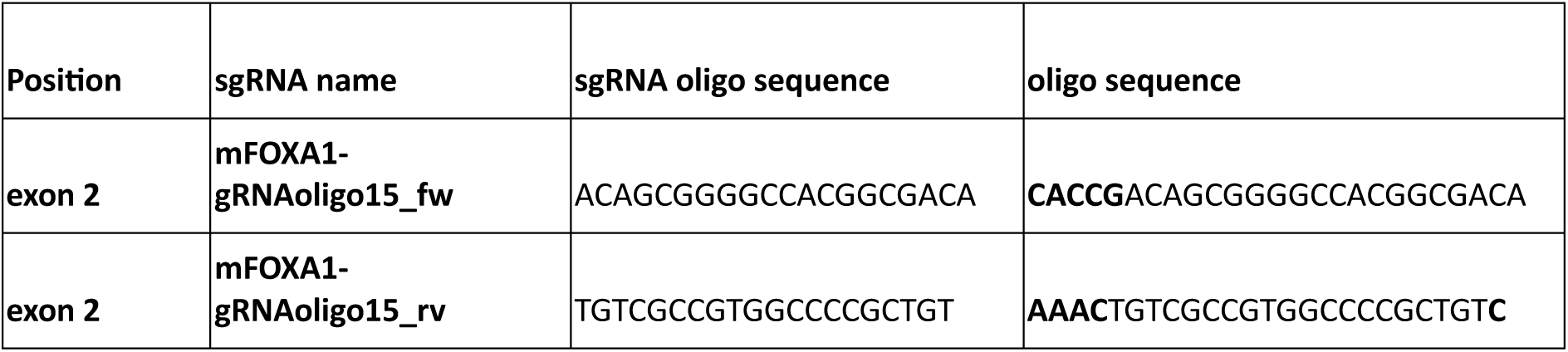
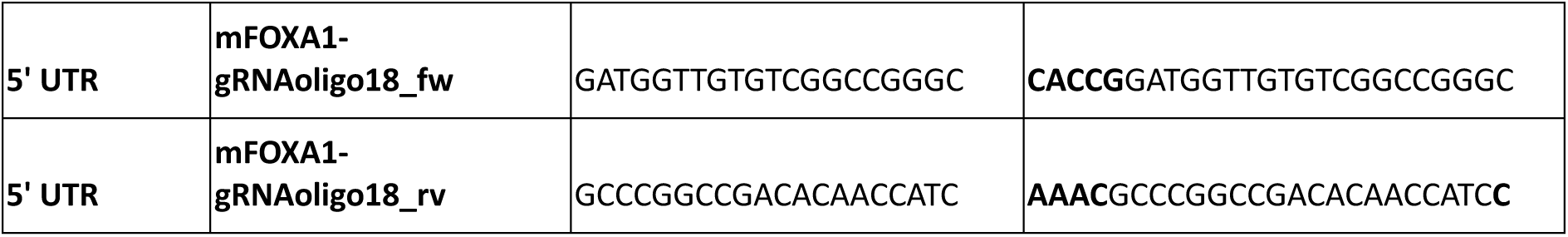

### Foxa1 qPCR analysis

RNA was extracted from 2 wells of a 24-well plate of organoids using the Qiagen RNAeasy mini kit (Qiagen cat# 74104) according to the manufacturer’s instructions. RNA was quantified using a Nanodrop One (ThermoFisher Scientific), and 2 µg of total RNA was reverse transcribed into cDNA using the PrimeScript RT Master Mix (Takara #RR036A) following the manufacturer’s instructions. RT-PCR was performed on a StepOne real-time PCR system (ThermoFisher Scientific) using Sybr Green (ThermoFisher Scientific #A25742) and the primers listed in table S2 (hexabiogen.com). The comparative CT (ddCT) method was used to determine the relative expression of the target gene compared to the housekeeping gene Elongation factor 1 alpha.

**Table S2.**
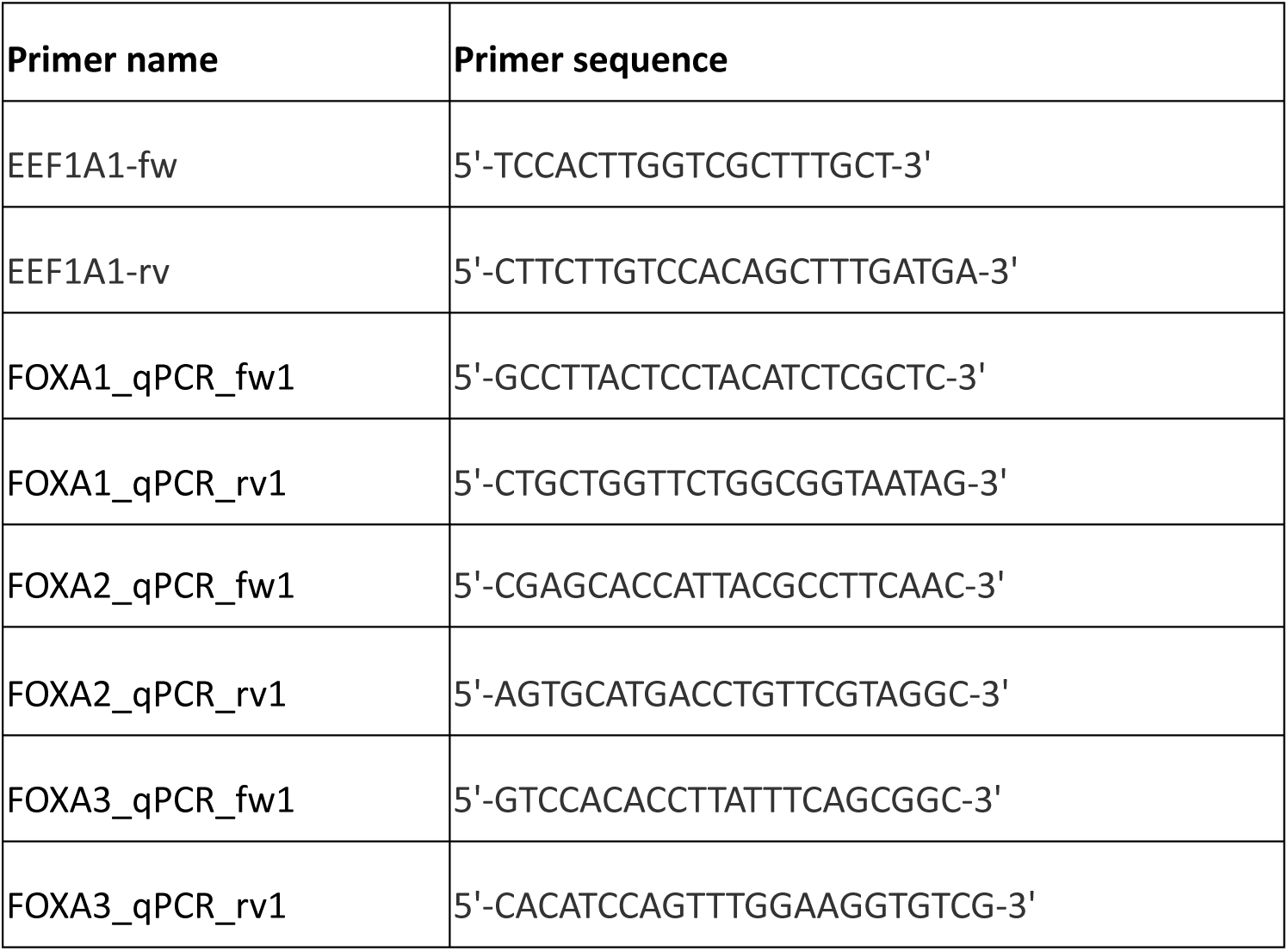

### Mouse intestinal organoid regeneration time course

Organoids were collected in cold DMEM/F-12 with 15 mM HEPES (Stem Cell Technologies) supplemented with 100 μg/ml Penicillin–Streptomycin, 1× Glutamax (Thermo Scientific) between 3-5 days after passaging and digested with TrypLE (Life Technologies, #12604013) for 20min at 37°C. Dissociated cells were collected and passed through a 30µm cell strainer (Sysmex). Single cells were sorted by FACS (Becton Dickinson Influx or Aria cell sorter or SONY, MA900 cell sorter). Forward and side scatter were used to remove cell doublets and debris. Sorted cells were collected in ENR medium composed of advanced DMEM/F-12 with 15mM HEPES (STEM CELL Technologies) supplemented with 100 μg/ml Penicillin-Streptomycin, 1×Glutamax (Thermo Fisher Scientific), 1×B27 (Thermo Fisher Scientific), 1xN2 (Thermo Fisher Scientific), 1mM N-acetylcysteine (Sigma), 500ng/ml R-Spondin (kind gift from Novartis), 100 ng/ml Noggin (PeproTech) and 100 ng/ml murine EGF (R&D Systems) plus Y-27632 (ROCK inhibitor, STEMCELL Technologies). After spinning down single cells at 900xg, the pellet was resuspended in 60% ENR medium + Y-27632 and 40% Matrigel (Corning). 5ul droplets with 3,000 cells were seeded per well of a 96-well plate. After 20min of solidification at 37°C, 100μl of medium was added per well. From 0h to 24h, Day0 medium was used: ENR was supplemented with 15-20% Wnt3a-conditioned medium (Wnt3a-CM), 3μM of CHIR99021 (GSK3B inhibitor, STEMCELL Technologies, cat# 72054) and 10μM Y-27632 (ROCK inhibitor, STEMCELL Technologies). From 24h-72h, Day1 medium was used: ENR was supplemented with 15-20% Wnt3a-conditioned medium (Wnt3a-CM) and 10μM Y-27632 (ROCK inhibitor, STEMCELL Technologies). From 72h till 120h only ENR medium was added. Wnt3a-conditioned medium was produced in-house by Wnt3a L-cells (kind gift from Novartis).

### Compound treatments

Single-cells were plated in 5ul droplets of 40% Matrigel (Corning) and 60% ENR + 10µM ROCK inhibitor, into a 96-well plate. For YAP1 perturbation experiments, 5µM LATS inhibitor (*4*) was added from 0h onwards and 5µM Verteporfin (SIGMA-ALDRICH, cat# SML0534) was added from 36h and DMSO (SIGMA-ALDRICH, cat# 276855) was used as a control. For Notch inhibition experiments, 10µM DAPT (Stemgent cat# 04-0041) was used from 0h onwards. To induce lumen inflation and nuclear stretch, 0.5µM Prostaglandin E2 (PGE) (5) or 10µM Forskolin (6) was added from 24h onwards. To inhibit cell division and induce a lower density state, 5-10nM Taxol (Paclitaxel, Selleck Chemicals, cat# S1150) or 250nM Aphidicolin (from Nigrospora sphaerica, ≥98%, SIGMA-ALDRICH, cat# A0781) was used from 24h-72h. For the washout condition, organoids were treated from 24h-48h and exchanged with Day1 medium from 48h-72h.

### Fixed sample preparation and antibody staining

One 96-well plate per time point was prepared and the entire plate was fixed as follows: Plate was centrifuged at 900xg for 10min at 10°C; 4% PFA (Electron Microscopy Sciences) in PBS was added and organoids were fixed for 45min at room temperature. Plate was washed three times with PBS and sealed with Aluminium foil for storage at 4°C or further processed for antibody staining. Here, organoids were permeabilized with 0.5% Triton X-100 (SIGMA-ALDRICH) for 1h and blocked for 1h with 3% Normal Donkey Serum (SIGMA-ALDRICH) in PBS containing 0.1% Triton X-100 at room temperature. Primary antibody stainings were performed overnight at 4°C shaking and secondary antibody staining was performed for 2h at room temperature shaking and sample was washed three times with PBS. For AGR2 antibody stainings, nanobodies were used as following: after permeabilisation and blocking procedure, sample was incubated in multiplexing blocker solution for 1h at room temperature; nanobody pre-complex of AGR2 antibody and anti-Rabbit IgG sdAb - FluoTag-X4 in PBS was incubated for 1h in the dark at room temperature; after incubation period, pre-complex was diluted in blocking solution without multiplexing blocker solution and added to the sample for a 2h incubation period at room temperature. Samples were washed three times in PBS. Concentrations of antibodies were used according to table S3. Nuclei staining was performed using 100μg/ml DAPI (4’,6-Diamidino-2-Phenylindole, Invitrogen) along with the secondary antibody staining.

**Table S3.**
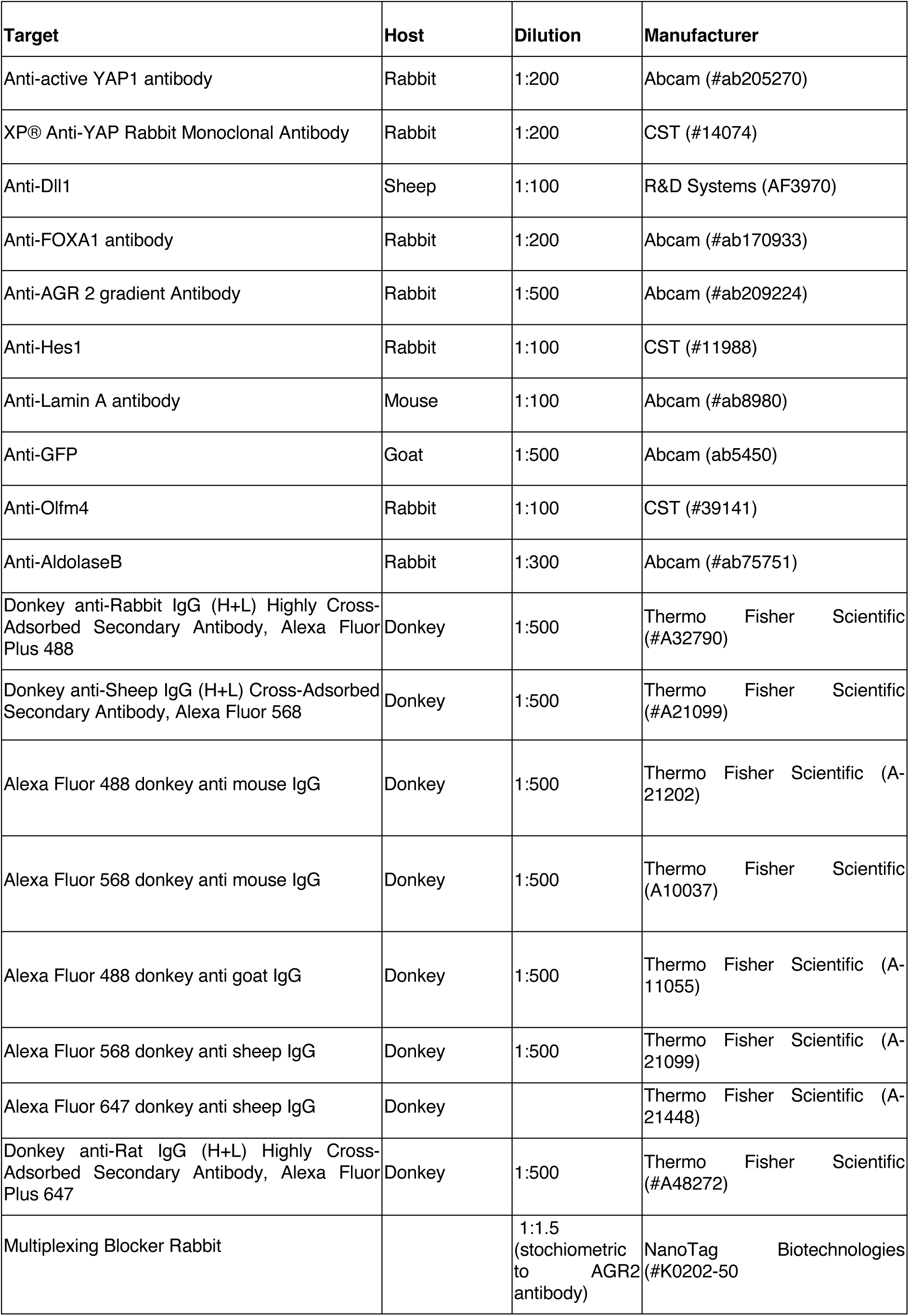
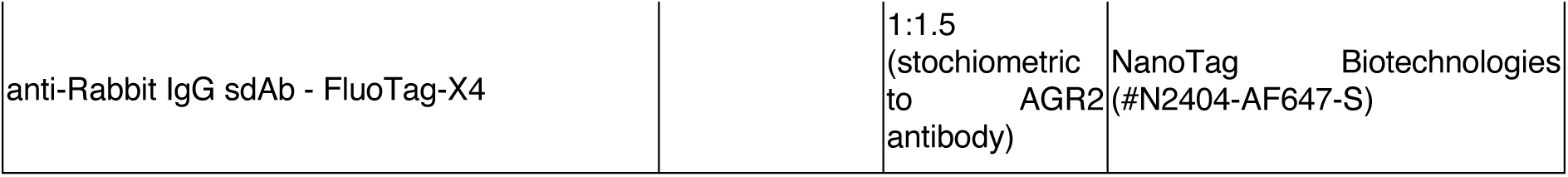

### High-throughput imaging of mouse intestinal organoid time course

High-throughput imaging was performed on a Yokogawa (CellVoyager 7000S), an automated spinning disk microscope equipped with CSU-W1 Confocal Scanner Unit (Yokogawa) with a 50µm pinhole disk unit, 2 sCMOS cameras (Andor Neo5.5, 2,560 × 2,160 pixels) and a 100mW 405nm Cube Laser, 200mW Laser 488nm Saphire Laser, 200nW Saphire Laser 561nm, 100mW 640nm Cube Laser (all from Coherent) using a 40x/0.95 UPLSAPOx2 (Olympus) Objective, or on a Yokogawa (CellVoyager 8000) equipped with CSU-W1 Confocal Scanner Unit (Yokogawa) with a 50µm pinhole disk unit, 2 sCMOS cameras (Hamamatsu Orca Flash 4 version 3, 2,048 × 2,048 pixels), and 100mW 405nm laser, 150mW Laser 488nm, 200mW Laser 561nm, 100mW 640nm Laser (all Lasers are Obis lasers from Coherent) using a 20x/1.0 water immersion Objective (Yokogawa). For imaging, either a fixed area was imaged or an intelligent imaging approach (Search First module of Wako software) was used, where the entire well was scanned with 4x and organoids were segmented and in the next step only fields that contained organoids were imaged with 40x. Z-stacks were acquired at 1-2µm z-step and ranged from 26µm for small organoids to 80µm for big organoids.

### Irradiation experiments

7-8 weeks old C57BL/6J male mice were lethally irradiated with 10 Gy (*7*, *8*) from a GSR D1 ^137^Caesium source irradiator (2 x 5 Gy doses with 3h rest between exposures). Mice were injected i.v. with 2.5 x 10^6^ bone marrow cells from C57BL/6J WT male mice in a volume of 0.2ml 24h after irradiation. At the indicated time points, the small intestine was divided in 3 sections of equal length and fixed in periodate-lysine-4% paraformaldehyde (PLP) buffer (*9*) overnight at 4°C. Tissues were washed twice with PBS and incubated in 30% sucrose overnight at 4°C. Each section was then rolled in a proximal to distal fashion, embedded in OCT and stored at −80 °C.

### Sectioning, immunostaining and imaging of OCT embedded *in vivo* tissue

OCT block was cut into 20µm thick section using Histology Cryostat CryoStar NX70 and Ultra Microtome Blades (Thermo Scientific, #30-538-35). Sections were placed onto Superfrost; Excell; Microscope Slides (Epredia, #10149870) and processed for antibody staining. First, sections were carefully washed 3x with PBS and incubated in permeabilisation and blocking solution (3% Normal Donkey Serum (SIGMA-ALDRICH) in PBS containing 0.5% Triton X-100) for 1h at room temperature. Antibody staining was performed as mentioned above (*section: Fixed sample preparation and antibody staining*). After antibody staining, sections were closed with Mounting medium (Ibidi, #50001) and a cover slip (Corning® 24×50 mm Rectangular #1½ Cover Glass, #CLS2980245-1000EA). Imaging was performed on an upright microscope Axio Imager M2 (Zeiss) equipped with CSU-W1 Confocal Scanner Unit (Yokogawa) with a 50um pinhole disk unit, a sCMOS cameras (PCO.edge 4.2M, PCO), an MS-2000 motorized stage (Applied Scientific Instrumentation) and controlled with VisiView 6.0 imaging software (Visitron Systems GmbH). Illumination was achieved with 405, 488, 561 and 639 lasers and VS-Homogenizer (Visitron Systems GmbH). A Fluar 5x/0.25 Air objective was used for acquiring overviews of the entire swiss roll and a 40x/1.3Oil objective (EC Plan Neofluar, Zeiss) was used to acquire ROIs as tiles. Z-stacks of around 20-40µm were acquired at a z-step of 1µm.

### Deparaffinization and staining of YAP1 activator treated ***in vivo*** tissue sections

Paraffin sections of YAP1 or vehicle treated samples were a kind gift of the Tchorz group (*4*). Details on animal maintenance, and *in vivo* treatments can be found here (*4*). In brief, C57BL/6 male mice (12-weeks-old) were treated with NIBR-LTSi (100mg/kg) or vehicle (0.5% methylcellulose (MC), 0.5% Tween 80) for 2.5 days and animals were sacrificed 4h after treatment, tissue was fixed in formalin and embedding as Swiss rolls in paraffin. Paraffin block was cut in 5µm sections. Paraffin sections were heated for 45min at 60°C. Slides were cooled down for 10min and incubated 2x for 3min in 100% EtOH, for 2min in 95% EtOH, for 3min in 75% EtOH, and for 5min in ddH2O. Sodium citrate buffer was added and heated for 50min at 90°C. After cooling down the slides, slides were transferred in ddH2O, and antibody staining was performed as above (OCT sections).

### Live sample preparation and light sheet imaging

Samples for light sheet imaging were prepared as mentioned previously (*section: Mouse intestinal organoid regeneration time course*). Samples were mounted as previously described (*10*, *11*). In brief, a custom designed and 3D printed imaging chamber was used. A 25 µm thin FEP foil (Katco) was plasma treated and then placed into the imaging chamber and glued with biocompatible silicone to the bottom of the chamber (Silpuran 4200; Wacker) and cured overnight at room temperature. Chamber was washed with EtOH and ddH2O for 3x times. Chamber was UV sterilized for 20min and pre-warmed before 5 µl droplets of single-cell suspension in 60% Matrigel (Corning) was added. After 15min, droplets solidified and Day0 medium was added. The LS1-Live dual illumination and inverted detection microscope from Viventis Microscopy Sàrl was used to perform light sheet imaging. At 24h post plating, organoids at the 2- or 4-cell stage were selected. A volume of 150 µm and Z-spacing of 2µm was acquired. Medium was refreshed every 24h according to timecourse medium changes (*section: Mouse intestinal organoid regeneration time course*).

### Design and construction of CTGF-mNeonGreen-NLS transcriptional reporter

The Yap1 transcriptional target Ctgf (Ccn2) was selected as a proxy for Yap1 activity. Using a combination of the Eukaryotic Promoter Database (Swiss Institute of Bioinformatics, https://epd.expasy.org/epd/) as well as the ChIP-Atlas (https://chip-atlas.org/) a relevant Ccn2 promoter sequence was defined taking the following DNA binding transcription factors into account: Tead1, Tead2, Tead3, Tead4, Smad2, Smad3 and Smad4. For the Search Motif Tool (EPD) a cut-off p-value of 1E-03 was used. Employing the ChIP-Atlas peak browser, peaks were assessed without a cell type class selection. A MACS2 significance threshold of Q value < 1E-05 was considered. Eventually a 1074bp long sequence was selected mapping to chr10:24,594,651-24,595,725(GRCm38/mm10).

The above-described sequence was ordered as gBlock (IDT). Due to high sequence complexity 4 repetitive bases have been deleted (see sequence below). To generate a stable reporter-line the PiggyBac transposon system was used. To that end a pPB-CAG-DEST-pA-pgk-hph vector (kind gift from J. Betschinger, Novartis) was subcloned to introduce a single MluI cloning site. To do so, the initial backbone was linearized with XbaI (cat# R0145T, NEB) and BbsI (cat#R0539S, NEB) and a gBlock (IDT) was used to introduce the MluI cut site via Gibson assembly (cat# E2611L, NEB). Using MluI (cat# R3198S, NEB) the expression vector was linearised, and Gibson assembly was used to incorporate the mNeonGreen-NLS coding sequence, amplified via PCR from pLV 4xSTAR-mNeonGreen-NLS-blast (Addgene #136258). In a subsequent cloning step this construct was cut with AccI (cat# R0161S) and XcmI (cat# R0533S) to remove the CAG promoter. A gBlock was used to retain parts of the mNeonGreen protein coding sequence as well as to introduce the above described Ctgf promoter sequence.

The Qiaquick gel extraction kit (cat# 28706, Qiagen) was used to purify PCR and digestion products from agarose gels. Mix & Go E. Coli transformation kit (cat# T3001, Zymo Research) treated DH5alpha cells (cat# 18265-017, Invitrogen) were used for transformation and plasmid amplification. Bacterial selection was performed using Ampicillin (100µ/ml). Plasmid purification was performed using QIAprep Spin Miniprep Kit (cat# 27106, Qiagen) and NucleoBond Xtra Midi kit (cat# 740410.50, Macherey-Nagel).

#### Ctgf promoter sequence

CCCAACTGAACCCCTACTGGCCTCCTTATACCTCTTTCTTCTCCCACTATATTCCCTGACACTTAGCTTCTGAACACAGCCATTTGGTCTGAACTCATAAACTTATTTTTCTAGAAAAGCCATGCCCAGTCATTCCCCTTGCCTCCCTGGACCCTGAAGACAAGTTCTTACATAAAGAGTGCTGAAAATCTCCCTGGGAACCTACATCCTTGGCTTTCATATCTTTCAGCCATCAAAATGGTTATCTCCAGTGACCAAAGATCAAATGCCTGTATTTCAGATACAAAAGTTGCACATAGGAATTCTGGGAGGAGAGGAGGCATTTCAAATGGCTATAAGCACCCTTCTCCTCTCAGTAGAAGAACACCAAGAGACTACAGCCCCGTAAAGAAAAAAAAAAATCCAAAACAAAGAAAAATATTTTTTTTAATTTCTAGGGGCCCATGGTATTTGCCTCTTGAGCTATTTGAGTCTTGAGAAGTTTTTATGTCAGTAGCCAGAACTGGCAAAGAGATTTTTAAGAAGAAAAGATCAGAGAAATAATCGTTTATTTCTAAGTTATATTTCATCAGGAGGGGTGAGAAGATGATATGGAGAAAGTTTTACTTCTTGGTGTTGTGCTGGAAACACAGCGCCTTTTTTTTCCTGGCGAGCTAAAGTGTGCCAGCTTTTTCAGACGGAGGAATGTGGAGTGTCAAGGGGTCAGGATCAATCCGGTGTGAGTTGATGAGGCAGGAAGGTGGGGAGGAATGTGAGGAATGTCCCTGTTTGTGTAGGACTCCATTCAGTTCTTTGGCGAGCCGGCTCCCGGGAGCGTATAAAAGCCAGCGCCGCCCGCCTAGTCTCACACAGCTCTTCTCTCCAAGAAGACTCAGCCAGATCCACTCCAGCTCCGACCCCAGGAGACCGACCTCCTCCAGACGGCAGCAGCCCCAGCCCAGCCGACAACCCCAGACGCCACCGCCTGGAGCGTCCAGACACCAACCTCCGCCCCTGTCCGAATCCAGGCTCCGGCCGCGCCTCTCGTCGCCTCTGCACCCTGCTGTGCATCCTCCTACCGCGTCCCGATC

### Generation of CTGF-mNEONgreen-NLS organoid line

CTGF-mNEONgreen-NLS reporter line was generated via electroporation as reported before (*10*). Briefly, organoids were digested into single cells and resuspended in BTX electroporation solution (cat# 732-1285). 7 ug of reporter constructs and 3 ug of pBase constructs were added to 1x 10^6^ single cells with a total volume of 100µl and placed on ice. Cell-plasmid mixture was added to cuvette, and the cuvette’s impedance was confirmed within the range of 0.030 to 0.05 before electroporation. Following electroporation, 400µl of BTX electroporation solution was applied to dilute and transfer the electroporation mixture, which was then incubated at room temperature for 30min and centrifuged at 300xg for 3 minutes at 4°C. After centrifugation, the supernatant was removed, and the remaining cells with 50µl of buffer was mixed with Matrigel (Corning) and seeded on 24-well plates. 500µl of ENR medium with WNT3a conditional medium, ROCKi and CHIR was added to each well. After 5-day culture at 37°C in a CO2 incubator, organoids were again digested to single cells and sorted for positive fluorescent signal. Following another round of 5-day culture, 30-50 single organoids were hand-picked and seeded in individual wells for expansion. The fluorescent signal of the reporter was verified via live imaging. 5 organoid lines with positive signal were selected and further stained for cell markers to check the cellular composition of the organoid tissue. The organoid line with the appropriate fluorescent signal and cellular composition were selected for experiments.

### Microwell experiments

Microwells were prepared as previously described (*12*). In brief, PDMS stamps were plasma treated and Collagen type I (Koken) - Matrigel mix (20%/80%) was added and inverted onto glass-bottom MatTek dish on top of two spacers. After 30min of solidification at 37°C, the stamp was removed and the single-cell suspension was added. After 5min, cells were seeded in the microwells, and gentle washing was performed to get rid of the remaining cells. A Collagen-Matrigel mix (20%/80%) was added onto a cover slip and inverted on top of the microwells and incubated for 15min. ISC expansion medium and Thiazovivin (2.5 μM; Stemgent) was added. ISC expansion medium: DMEM/F-12 with 15mM HEPES (STEM CELL Technologies) supplemented with 100 μg/ml Penicillin-Streptomycin, 1×Glutamax (Thermo Fisher Scientific), 1×B27 (Thermo Fisher Scientific), 1xN2 (Thermo Fisher Scientific), 1mM N-acetylcysteine (Sigma), 500ng/ml R-Spondin (kind gift from Novartis), 100 ng/ml Noggin (PeproTech) and 50 ng/ml murine EGF (R&D Systems) plus 3μM GSK-3β inhibitor CHIR99021 (Stem Cell Technologies, cat# 72052) and 1mM valproic Acid (Sigma #P4543).

### Image analysis of high-content organoid data

#### Raw-image processing, 2D and 3D segmentation, feature extraction

Raw-images were converted to the Open Microscopy Environment format (OME-Zarr) (13) using the Fractal platform (https://fractal-analytics-platform.github.io/). Where needed, the python package ‘ez-zarr’ (https://fmicompbio.github.io/ez_zarr/) was used for post-processing of OME-Zarr (e.g. feature extraction). Whole-well MIPs of the DAPI-channel were segmented with a custom model using RDCNet (version v0.4) (14). For nuclei segmentations, a RDCnet model (16) was trained from scratch based on 3D data of DAPI stained organoids from 24h-72h of organoid regeneration time courses. For cell segmentations, 2D and 3D Cellpose (15) models were re-trained based on 3D data of DAPI stained nuclei and E-cadherin stained cell membranes of organoids from 24h-72h regeneration time courses. Nuclei and cell segmentations were linked through maximum overlap. Feature extraction of single-nucleus, cell, cytoplasm and organoid-level information was performed using the scikit-image package (17) and custom-written code in Python. Extracted features were casted in a SingleCellExperiment object for further post-processing.

#### Quality control and normalization

Organoid filtering was performed by removing single-cells and debri based on area measurements as well as cut-off organoids. Mis-segmented organoids were filtered out by calculating the Euler number and only retaining objects with Euler number equals to one. Mis-segmentations of single-cells were filtered by identifying empty masks (low DAPI mean intensity) and too small cells (smaller than 1/10 of the median size observed). To correct for signal decay along the z-axis due to reduced penetration in the sample, for each intensity feature we summarised measurements along z-axis into 7 to 8 bins (averaging 1,200 to 9,000 single cell measurements per bin) and fit a loess model of intensity feature over z-axis bins (using the r function loess(), with span = 0.5, family = “symmetric”, and control = loess.control(surface = “direct”). Finally, we subtract the fitted values from the raw extracted values. For z-score value calculation, data were standardized against the average (mu) and standard deviation (sigma) of the entire distribution or in case of perturbation experiments, data were standardized against the mu and sigma of DMSO or WT controls.

#### Single cell classification and calculation of cell-to-cell variability

For classification into FOXA1^−/int/+^ or Dll1^−/int/+^ k-means clustering was performed. Organoid-level mean intensities were calculated by averaging the z-scored mean intensities of all nuclei/cells per organoid. Coefficient of variation was calculated as standard deviation of all nuclei/cells per organoid divided by the average of all nuclei/cells. A pseudo-count of 0.01 was introduced in the calculation of the mean to avoid numerical problems during division. For line plots of organoid-level information, data were smoothed using the loess function in R; for single-cell data, the data frame was first subsampled and then loess function was used to produce the smoothed line plots. For rank plots, single-cells or -nuclei were ranked by intensities, size or shape within the same organoid and loess was used to produce a smoothed line plot. Correlation plots were produced by correlating the single-cell aYAP1 and DLL1 z-scored mean intensities per organoid and then averaging the per-organoid level information. Linear models were used to perform the smoothing.

### scATAC sequencing time course

Wild-type organoids, isolated from a 10 weeks old male C57BL/6 mouse, were collected in cold advanced DMEM/F-12 with 15 mM HEPES (Stem Cell Technologies, cat# 36254), supplemented with 100 μg/mL Penicillin-Streptomycin and 1× Glutamax (Thermo Scientific, cat# 35050061). The organoids were pelleted at 600xg for 5 minutes at 4°C and digested into single cells at 37°C in a water bath for 20min using TrypLE (Life Technologies, #12604013) containing 10μM ROCK inhibitor (Y-27632). After digestion, single cells were pelleted at 600xg for 5min at 4°C, resuspended in PBS supplemented with 10μM ROCK inhibitor, and passed through a 30μm cell strainer. Single cells were sorted using a MA900 cell sorter (Sony) equipped with a 100µm nozzle. Live single cells were gated based on DRAQ7 staining, size, and granularity. Sorted cells were collected in 1.5mL tubes containing 200μL ENR medium supplemented with 10μM ROCK inhibitor, then pelleted at 900xg for 5 minutes. The cell pellets were resuspended in ENR medium consisting of advanced DMEM/F-12 with 15mM HEPES (Stem Cell Technologies, cat# 36254), supplemented with 100 μg/mL Penicillin-Streptomycin, 1× Glutamax (Thermo Scientific, cat# 35050061), 1× B27 (Thermo Scientific, cat# A1895602), 1× N2 (Thermo Scientific, cat# 17502001), 1 mM N-acetylcysteine (Sigma, cat# A7250), 500 ng/mL R-Spondin (Novartis), 100 ng/mL Noggin (PeproTech, cat# 250-38), and 100 ng/mL murine epidermal growth factor (EGF; R&D Systems). The medium was mixed with Matrigel (Corning) at a 1:1 ratio. For each well of a six-well plate, five droplets of 50μL, each containing 30,000 single cells (i.e. 0.6k cell/µL), were seeded and cultured for 24h in ENR medium supplemented with 25% WNT3A-conditioned medium (WNT3A-CM), 10μM ROCK inhibitor (Stem Cell Technologies, cat# 72304), and 3μM GSK-3β inhibitor CHIR99021 (Stem Cell Technologies, cat# 72052). From 24h to 72h, organoids were cultured in ENR medium supplemented with 25% WNT3A-CM and 10μM ROCK inhibitor, and from 72h onwards, ENR medium alone was used. Organoids were harvested at 24, 36, 48, 60, 72, and 84 hours post-seeding. Harvested organoids were digested into single cells and prepared for FACS sorting as described above. Sorted cells (50 – 100k) were collected in 1.5 mL tubes containing 200μl ENR medium supplemented with 10μM ROCK inhibitor. The cells were pelleted at 900xg for 5min at 4°C, and the pellet was resuspended in 600μL freezing medium (Stem Cell Technologies, cat# 210373) supplemented with 10μM ROCK inhibitor. Cell suspensions were frozen in cryo-boxes at −80°C and transferred to liquid nitrogen storage after 24h.

### Nuclei Isolation and scATAC-seq Library Preparation

The sorted and cryopreserved cells from each organoid time point were thawed in a 37°C water bath and washed three times with 0.8% BSA-PBS. After the third wash, cells were transferred into BSA-coated 0.2mL PCR tubes. Nuclei were then isolated using a low-input nuclei isolation protocol (Demonstrated protocol CG000169 • Rev D, 10x Genomics), with an 8-minute lysis time. Propidium iodide was used to identify the nuclei, and counting was performed using the MoxiGo II counter (Orflo Technologies). Nuclei were loaded to achieve a target recovery of 10,000 nuclei; if recovery was insufficient, all available nuclei were loaded instead. scATAC-seq libraries were generated using the Chromium Single Cell ATAC Library & Gel Bead Kit (PN-1000110) following the user guide (CG000168 • Rev A, 10x Genomics). Libraries were sequenced on the Illumina NovaSeq platform.

### scATACseq analysis

All analyses, unless stated otherwise, were performed within R (version 4.2.0) and Bioconductor (version 3.15) (*18*), using SeuratObject version 4.1.0, Seurat version 4.1.1, and Signac version 1.7.0.

#### Alignment and quantification

Fragment files with aligned read coordinates were generated for each sample (one file per each of the timepoints: 24h, 36h, 48h, 60h, 72h, 84h), using CellRanger ATAC (version 1.2.0) with the “count” subtool and reference “refdata-cellranger-atac-mm10-1.2.0”. For initial quantification, genomic features were obtained by combining promoters (transcription start sites (TSS) expanded to 1000 base pairs centered on the TSS) and enhancer from the cellranger annotation. All overlapping regions were collapsed, and any region overlapping a cellranger annotated blacklist region was removed. Alignments in these regions were then quantified with Signac::FeatureMatrix() from the fragment files.

#### Quality control, filtering, normalization and dimensionality reduction

Cells with fewer than 300 detected features were excluded, and features detected in fewer than 10 cells were removed, yielding 83,493 cells across 302,405 features. Quality control metrics were calculated using TSSenrichment() and NucleosomeSignal(). Cells with TSS enrichment ≤ 2, nucleosomal signal ≥ 5, ≤ 2,500 fragments, or ≤ 25% fragments in promoters (calculated using FRiP()) were excluded, resulting in 68,905 cells. Normaliszation was performed with RunTFIDF() (method = 1, scale.factor = 10,000), and variable features were selected using FindTopFeatures() (min.cutoff = “q50”), yielding 151,214 features. Dimensionality reduction was performed using RunSVD() (n = 50). The first component of the resulting Latent Semantic Index (LSI) was excluded from all subsequent analysis as it showed high correlation with sequencing depth (as shown by DepthCor()). Using the remaining LSI components as input, a total of 38 clusters were identified using FindNeighbors() and FindClusters() (resolution = 3).

#### Predicted RNA expression, initial cell type annotation and peak calling

Gene coordinates were extracted from the reference (genes gtf file in refdata-cellranger-atac-mm10-1.2.0) including type = “gene” and gene_type = “protein_coding”. Only standard chromosomes were retained, using GenomeInfoDb::keepStandardChromosomes(), with pruning.mode = “coarse”. Predicted gene expression was quantified with GeneActitity(), with extend.upstream = 2000, extend.downstream = 0, biotypes = NULL (as only protein coding genes were already pre-selected), and max.width = 5e+05. The RNA count matrix was normalized with NormalizeData(), normalize.method = “LogNormalize” and scale.factor = median(seurat$nCount_RNA) (the median total predicted RNA count per cell). Coarse-grain cell type annotation was performed by assigning the 38 clusters to cell types based on predicted RNA expression of known markers of intestinal lineages averaged over cells in each cluster (see Fig. S3D). We removed 1,929 cells (2 of the 38 clusters) showing high expression of stress markers (including *Hsp90aa1*, and *Hsp90b1*). Peaks were identified for each of these preliminary cell types with Signac::CallPeaks() using MACS2 (version 2.1.3.3) (*19*), with effective.genome.size = 1.87e9, extsize = 200, and shift = −100, and merged across clusters, yielding 252,049 peaks. A second fragment quantification was then performed using Signac::FeatureMatrix(), replacing the initial generic promoter/enhancer features by the sample specific peaks identified in our data.

#### Final cell type annotation and transfer labels to full dataset

The preliminary cell type annotation revealed that regenerative and TA / Stem cell progenitors were much more abundant in our data compared to other cell types. In order to reduce the weight of these frequent cell types, for example in the identification of variable features, we randomly subsampled these two cell types from about 15,000 and 36,000 cells respectively to 10,000 each, reducing the total number of cells to 36,988 (seuratSubsampled object). The normalization and latent semantic indexing were then repeated on seuratSubsampled (runTFIDF(), FindTopFeatures(), RnuSVD()), and a 2D embedding was generated using RunUMAP with dims = 2:30, umap.method = ‘uwot’, n.neighbors = 30L, min.dist = 0.3 and spread = 1. Cell types were re-annotated by clustering seuratSubsampled using FindNeighbors() and FindClusters() with resolution = 2.0, resulting in 28 clusters, that were assigned to cell types using marker genes as described, resulting in final cell type assignment of the subsampled cells (see Fig.S3C,D). We transferred the cell type annotation from the seuratSubsampled to the full dataset, projecting all cells into the LSI and UMAP embeddings identified from the subsampled cells. Cell types of non-sampled cells were assigned using the cell type of the nearest subsampled cell in LSI space.

#### Differential Accessibility and Motif Enrichment Analysis

Log2 fold changes of accessibility between each of Reprogramming, Regenerative, Secretory progenitors, Exocrine, TA / Stem cell progenitors, Enterocyte progenitors and reprogramming cells were computed similarly as in Signac:::FoldChange.ChromatinAssay but using a pseudocount of 0.2. Peaks with absolute log2 fold change > 1.75 in any contrast were retained, yielding 10,938 differentially accessible peaks (Fig. 3G). We calculated the average accessibility for each cell type by taking the mean of log2 normalized accessibility values over cells, and then calculated relative accessibility for each peak by linearly scaling its celltype averages into the interval from zero to one. Based on these relative accessibilities, peaks were clustered in 8 groups using the kmeans function in R. Motif enrichment analysis was performed with calcBinnedMotifEnrR() from the monaLisa package (*20*) in the 8 identified groups and JASPAR2020 as motif database. We sorted motifs by significance (“negLog10Padj”) and selected the top 50 plus TEAD motifs, which we manually included for their known role as YAP1 co-factors. The identified motifs were then clustered with motifSimilarity(), and 2–3 representative motifs were shown (Fig.3K).

### ChIPseq sample dissociation

Lgr5::DTR-eGFP organoids from two technical replicates (same organoid culture, 4 passages apart) were grown in ENR medium supplemented with 1:2000 DMSO and collected 4-days after mechanical splitting. Organoids were collected in 5ml DMEM/F-12 with 15mM HEPES (Stem Cell Technologies, cat# 36254), supplemented with 100 μg/mL Penicillin-Streptomycin, 1× Glutamax (Thermo Scientific, cat# 35050061), and pelleted at 600xg for 5 min at 4°C. After discarding supernatant, organoids were resuspended in 2ml TrypLE (Life Technologies, #12604013) and 10µM ROCK inhibitor and incubated at room temperature for 10min; every 5min the mix was resuspended pipetting for 10x, to ensure single cell dissociation. To quench TrypLE reaction, 6ml DMEM/F-12 with 15mM HEPES (Stem Cell Technologies, cat# 36254), supplemented with 100 μg/mL Penicillin-Streptomycin, 1× Glutamax (Thermo Scientific, cat# 35050061) was added to the suspension. After pelleting single-cell suspension at 600xg for 5min at 4°C cell pellet was resuspended in 1ml PBS and filtered through FACS tube (35µm, #352235 Corning). Cells were counted and 10M cells per sample were resuspended in 10ml PBS before proceeding with ChIP library preparation.

### ChIPseq library preparation and sequencing

ChIP was done as described in Kaaij et al. *Cell* 2019 with minor modifications (*21*). A total of 10⁷ cells were crosslinked for 8min at room temperature in 10mL PBS supplemented with 1% formaldehyde (Sigma, F8775). The reaction was quenched by adding glycine to a final concentration of 0.125mM. Cells were pelleted by centrifugation at 500×g for 3min at 4°C, washed once with PBS, pelleted again, flash-frozen in liquid nitrogen, and stored until further processing. 30µL of a 1:1 mixture of Protein A/G Dynabeads (Invitrogen, 10002D and 10004D) was pre-coupled with 3µL of FOXA1 antibody (ab170933) for 1 hour at room temperature, followed by three washes with 1× ChIP buffer and stored on ice until use. The cells were lysed in 10mL of buffer A (50mM HEPES pH 8.0, 140mM NaCl, 1mM EDTA, 10% glycerol, 0.5% NP-40, 0.25% Triton X-100) on ice for 10min. After centrifugation, the pellet was resuspended in 5mL of buffer B (10mM Tris pH 8.0, 1mM EDTA, 0.5mM EGTA, 200mM NaCl) and incubated on ice for 5min. Nuclei were pelleted by centrifugation at 500×g for 3min at 4°C, resuspended in 150µL of buffer C (50mM Tris pH 8.0, 5mM EDTA, 1% SDS, 100mM NaCl), and incubated on ice for 10min. TE buffer (1.3mL) was then added, and chromatin was sheared in 15mL tubes using a Bioruptor Pico device for two sessions of 10 cycles (30 seconds ON / 30 seconds OFF). After shearing, 160µL of 10× ChIP buffer (0.1% SDS, 10% Triton X-100, 12mM EDTA, 167mM Tris-HCl pH 8.0, 1.67M NaCl) was added. The chromatin was transferred to fresh tubes and centrifuged at 13,000×g for 10min at 4°C. Per sample, 30µL of 1:1 pre-coupled beads were added to the chromatin, and the mixture was incubated with rotation for 4 hours at 4°C. The ChIP samples were washed four times with LSB (10mM Tris-HCl pH 8.0, 1mM EDTA pH 8.0, 140mM NaCl, 1% Triton X-100, 0.1% SDS, 0.1% sodium deoxycholate), twice with HSB (same as LSB but with 500mM NaCl), twice with LiSB (10mM Tris-HCl pH 8.0, 1mM EDTA pH 8.0, 250mM LiCl, 0.5% NP-40, 0.5% sodium deoxycholate), and once with TE+ buffer (10mM Tris-HCl pH 8.0, 1mM EDTA, 50mM NaCl). During the final wash, beads were transferred to fresh tubes, and the wash buffer was completely removed. DNA was eluted twice from the beads using 75µL of elution buffer (10mM Tris-HCl pH 8.0, 1mM EDTA pH 8.0, 150mM NaCl, 1% SDS) by incubation at 65°C with shaking. The eluates were pooled, and 2µL of RNase A (20 µg/µL) was added, followed by incubation for 1 hour at 37°C. Input samples were adjusted to a total volume of 150µL with elution buffer and processed in the same manner as ChIP samples. Proteinase K (2µL, 20mg/mL) was added, and samples were incubated for 2 hours at 55°C and then for 6 hours at 65°C. Afterward, 9µL of 5M NaCl, 30µL of AMPure XP beads, and 190µL of isopropanol were added. Samples were vigorously mixed, incubated at room temperature for 10min, and the beads were collected on a magnetic rack. The beads were washed twice with 80% ethanol, and DNA was eluted in 30µL of 10mM Tris-HCl pH 8.0 for 5min at 37°C. For library preparation, 25µL of ChIP DNA or 50ng of input DNA was used with the NEBNext Ultra II Library Prep Kit for Illumina (NEB), following the manufacturer’s protocol but with reactions scaled down to half-volume. Libraries were sequenced on the NovaSeq 6000 platform (Illumina) with paired end reads (2x 56 bp).

### ChIPseq analysis

All analyses, unless stated otherwise, were performed within R (version 4.3.2) and Bioconductor (version 3.18) (*18*). ChIP-seq read pairs were aligned to the GRCm38/mm10 mouse genome assembly using the qAlign function from the QuasR package (*22*) with default parameters. BigWig files with alignment fragment coverage along the genome were generated from the bam files using bedtools version 2.30 (*23*), by first extracting a bed file with fragment coordinates (bedtools bamtobed), calculating genome coverage in a bedGraph file (bedtools genomecov) and then converting it to BigWig format using bedGraphToBigWig from KentTools version 442 (*24*). To calculate ChIP enrichments in ATAC peaks, we first counted the number of ChIP fragments r overlapping a peak using qCount with shift = “halfinsert” (overlaps are determined using the midpoint of each ChIP fragment). The raw counts r where then normalized to log2-counts per million lcpm using lcpm = log2(r / N * 1e6 + 1), where N is the total number of alignments in a sample. As we observed non-linearities between replicates, we separated lcpm values into IP (lcpm_ip) and input samples (lcpm_input) and separately normalized them using the normalizeCyclicLoess function from the limma package with default parameters (*25*), producing lcpm_ip_loess and lcpm_input_loess. IP-enrichments were calculated as the difference lcpm_ip_loess – lcpm_input_loess, which corresponds to log2(IP/input).

### Statistical analysis

To compare two groups, Mann-Whitney U test was used and to compare multiple groups, Kruskal-Wallis and Dunn’s multiple comparisons test was used.

## References

1. S. Kondo, T. Miura, Reaction-diffusion model as a framework for understanding biological pattern formation. Science 329, 1616–1620 (2010).

2. J. B. A. Green, J. Sharpe, Positional information and reaction-diffusion: two big ideas in developmental biology combine. Development 142, 1203–1211 (2015).

3. E. Nacu, E. M. Tanaka, Limb regeneration: a new development? Annu Rev Cell Dev Biol 27, 409–440 (2011).

4. J. C. Rink, Stem Cells, Patterning and Regeneration in Planarians: Self-Organization at the Organismal Scale. Methods Mol Biol 1774, 57–172 (2018).

5. K. W. Rogers, A. F. Schier, Morphogen gradients: from generation to interpretation. Annu Rev Cell Dev Biol 27, 377–407 (2011).

6. A. M. Turing, The chemical basis of morphogenesis. Philos. Trans. R. Soc. Lond. 237, 37–72 (1952).

7. L. Wolpert, Positional information and the spatial pattern of cellular differentiation. J. Theor. Biol. 25, 1–47 (1969).

8. B. C. Goodwin, M. H. Cohen, A phase-shift model for the spatial and temporal organization of developing systems. J Theor Biol 25, 49–107 (1969).

9. L. Wolpert, Positional information revisited. Development 107 Suppl, 3–12 (1989).

10. J. Briscoe, J. Ericson, The specification of neuronal identity by graded Sonic Hedgehog signalling. Semin Cell Dev Biol 10, 353–362 (1999).

11. R. Kraut, M. Levine, Spatial regulation of the gap gene giant during Drosophila development. Development 111, 601–609 (1991).

12. L. Panman, R. Zeller, Patterning the limb before and after SHH signalling. Journal of Anatomy 202, 3–12 (2003).

13. Seeing Is Believing: The Bicoid Morphogen Gradient Matures. Cell 116, 143–152 (2004).

14. J. Jaeger, J. Reinitz, On the dynamic nature of positional information. BioEssays 28, 1102–1111 (2006).

15. S. Artavanis-Tsakonas, M. D. Rand, R. J. Lake, Notch signaling: cell fate control and signal integration in development. Science 284, 770–776 (1999).

16. J. R. Collier, N. A. Monk, P. K. Maini, J. H. Lewis, Pattern formation by lateral inhibition with feedback: a mathematical model of delta-notch intercellular signalling. Journal of theoretical biology 183 (1996).

17. D. Sprinzak, A. Lakhanpal, L. LeBon, L. A. Santat, M. E. Fontes, G. A. Anderson, J. Garcia-Ojalvo, M. B. Elowitz, Cis-interactions between Notch and Delta generate mutually exclusive signalling states. Nature 465, 86–90 (2010).

18. F. Corson, L. Couturier, H. Rouault, K. Mazouni, F. Schweisguth, Self-organized Notch dynamics generate stereotyped sensory organ patterns in Drosophila. Science (New York, N.Y.) 356 (2017).

19. P. W. Sternberg, Falling off the knife edge. Curr Biol 3, 763–765 (1993).

20. S. Artavanis-Tsakonas, K. Matsuno, M. E. Fortini, Notch signaling. Science 268, 225–232 (1995).

21. A. Chitnis, D. Henrique, J. Lewis, D. Ish-Horowicz, C. Kintner, Primary neurogenesis in Xenopus embryos regulated by a homologue of the Drosophila neurogenic gene Delta. Nature 375, 761–766 (1995).

22. A. Gregorieff, Y. Liu, M. R. Inanlou, Y. Khomchuk, J. L. Wrana, Yap-dependent reprogramming of Lgr5(+) stem cells drives intestinal regeneration and cancer. Nature 526, 715–718 (2015).

23. L. Aloia, M. A. McKie, G. Vernaz, L. Cordero-Espinoza, N. Aleksieva, J. van den Ameele, F. Antonica, B. Font-Cunill, A. Raven, R. Aiese Cigliano, G. Belenguer, R. L. Mort, A. H. Brand, M. Zernicka-Goetz, S. J. Forbes, E. A. Miska, M. Huch, Epigenetic remodelling licences adult cholangiocytes for organoid formation and liver regeneration. Nat Cell Biol 21, 1321–1333 (2019).

24. P. W. Tetteh, O. Basak, H. F. Farin, K. Wiebrands, K. Kretzschmar, H. Begthel, M. van den Born, J. Korving, F. de Sauvage, J. H. van Es, A. van Oudenaarden, H. Clevers, Replacement of lost Lgr5-positive stem cells through plasticity of their enterocyte-lineage daughters. Cell Stem Cell 18, 203–213 (2016).

25. S. Yui, L. Azzolin, M. Maimets, M. T. Pedersen, R. P. Fordham, S. L. Hansen, H. L. Larsen, J. Guiu, M. R. P. Alves, C. F. Rundsten, J. V. Johansen, Y. Li, C. D. Madsen, T. Nakamura, M. Watanabe, O. H. Nielsen, P. J. Schweiger, S. Piccolo, K. B. Jensen, YAP/TAZ-dependent reprogramming of colonic epithelium links ECM remodeling to tissue regeneration. Cell Stem Cell 22, 35–49.e7 (2018).

26. I. Lukonin, D. Serra, L. Challet Meylan, K. Volkmann, J. Baaten, R. Zhao, S. Meeusen, K. Colman, F. Maurer, M. B. Stadler, J. Jenkins, P. Liberali, Phenotypic landscape of intestinal organoid regeneration. Nature 586, 275–280 (2020).

27. S. Viragova, D. Li, O. D. Klein, Activation of fetal-like molecular programs during regeneration in the intestine and beyond. Cell Stem Cell 31, 949–960 (2024).

28. K. C. Oost, M. Kahnwald, S. Barbiero, G. de Medeiros, S. Suppinger, V. Kalck, L. C. Meylan, S. A. Smallwood, M. B. Stadler, P. Liberali, Dynamics and plasticity of stem cells in the regenerating human colonic epithelium, bioRxiv (2023)p. 2023.12.18.572103.

29. R. Priya, S. Allanki, A. Gentile, S. Mansingh, V. Uribe, H.-M. Maischein, D. Y. R. Stainier, Tension heterogeneity directs form and fate to pattern the myocardial wall. Nature 588, 130–134 (2020).

30. D. Serra, U. Mayr, A. Boni, I. Lukonin, M. Rempfler, L. Challet Meylan, M. B. Stadler, P. Strnad, P. Papasaikas, D. Vischi, A. Waldt, G. Roma, P. Liberali, Self-organization and symmetry breaking in intestinal organoid development. Nature 569, 66–72 (2019).

31. N. Q. Balaban, J. Merrin, R. Chait, L. Kowalik, S. Leibler, Bacterial persistence as a phenotypic switch. Science 305, 1622–1625 (2004).

32. P. S. Stewart, M. J. Franklin, Physiological heterogeneity in biofilms. Nat Rev Microbiol 6, 199–210 (2008).

33. B. Snijder, L. Pelkmans, Origins of regulated cell-to-cell variability. Nat Rev Mol Cell Biol 12, 119–125 (2011).

34. K. Keren, Z. Pincus, G. M. Allen, E. L. Barnhart, G. Marriott, A. Mogilner, J. A. Theriot, Mechanism of shape determination in motile cells. Nature 453, 475–480 (2008).

35. M. Zinner, I. Lukonin, P. Liberali, Design principles of tissue organisation: How single cells coordinate across scales. Curr Opin Cell Biol 67, 37–45 (2020).

36. D. A. Turner, P. Baillie-Johnson, A. M. Arias, Organoids and the genetically encoded self-assembly of embryonic stem cells. BioEssays 38, 181–191 (2016).

37. A. Schauer, D. Pinheiro, R. Hauschild, C.-P. Heisenberg, Zebrafish embryonic explants undergo genetically encoded self-assembly. doi: 10.7554/eLife.55190 (2020).

38. Feedback Control of Gene Expression Variability in the Caenorhabditis elegans Wnt Pathway. Cell 155, 869–880 (2013).

39. A. Raj, A. van Oudenaarden, Nature, Nurture, or Chance: Stochastic Gene Expression and Its Consequences. Cell 135, 216–226 (2008).

40. A. Eldar, M. B. Elowitz, Functional roles for noise in genetic circuits. Nature 467, 167–173 (2010).

41. C. Chazaud, Y. Yamanaka, T. Pawson, J. Rossant, Early lineage segregation between epiblast and primitive endoderm in mouse blastocysts through the Grb2-MAPK pathway. Dev Cell 10, 615–624 (2006).

42. C. Schröter, K. S. Stapornwongkul, V. Trivedi, Local cellular interactions during the self-organization of stem cells. Curr Opin Cell Biol 85, 102261 (2023).

43. D. Fabrèges, B. Corominas-Murtra, P. Moghe, A. Kickuth, T. Ichikawa, C. Iwatani, T. Tsukiyama, N. Daniel, J. Gering, A. Stokkermans, A. Wolny, A. Kreshuk, V. Duranthon, V. Uhlmann, E. Hannezo, T. Hiiragi, Temporal variability and cell mechanics control robustness in mammalian embryogenesis. Science 386, eadh1145 (2024).

44. A. Ayyaz, S. Kumar, B. Sangiorgi, B. Ghoshal, J. Gosio, S. Ouladan, M. Fink, S. Barutcu, D. Trcka, J. Shen, K. Chan, J. L. Wrana, A. Gregorieff, Single-cell transcriptomes of the regenerating intestine reveal a revival stem cell. Nature 569, 121–125 (2019).

45. S. Das, S. M. Parigi, X. Luo, J. Fransson, B. C. Kern, A. Okhovat, O. E. Diaz, C. Sorini, P. Czarnewski, A. T. Webb, R. A. Morales, S. Lebon, G. Monasterio, F. Castillo, K. P. Tripathi, N. He, P. Pelczar, N. Schaltenberg, M. De la Fuente, F. López-Köstner, S. Nylén, H. L. Larsen, R. Kuiper, P. Antonson, M. A. Hermoso, S. Huber, M. Biton, S. Scharaw, J.-Å. Gustafsson, P. Katajisto, E. J. Villablanca, Liver X receptor unlinks intestinal regeneration and tumorigenesis. Nature, doi: 10.1038/s41586-024-08247-6 (2024).

46. K. Namoto, C. Baader, V. Orsini, A. Landshammer, E. Breuer, K. T. Dinh, R. Ungricht, M. Pikiolek, S. Laurent, B. Lu, A. Aebi, K. Schönberger, E. Vangrevelinghe, O. Evrova, T. Sun, S. Annunziato, J. Lachal, E. Redmond, L. Wang, K. Wetzel, P. Capodieci, J. Turner, G. Schutzius, V. Unterreiner, M. Trunzer, N. Buschmann, D. Behnke, R. Machauer, C. Scheufler, C. N. Parker, M. Ferro, A. Grevot, A. Beyerbach, W.-Y. Lu, S. J. Forbes, J. Wagner, T. Bouwmeester, J. Liu, B. Sohal, S. Sahambi, L. E. Greenbaum, F. Lohmann, P. Hoppe, F. Cong, A. W. Sailer, H. Ruffner, R. Glatthar, B. Humar, P.-A. Clavien, M. T. Dill, E. George, J. Maibaum, P. Liberali, J. S. Tchorz, NIBR-LTSi is a selective LATS kinase inhibitor activating YAP signaling and expanding tissue stem cells in vitro and in vivo. Cell Stem Cell 31, 554–569.e17 (2024).

47. J. Moore, D. Basurto-Lozada, S. Besson, J. Bogovic, J. Bragantini, E. M. Brown, J.-M. Burel, X. Casas Moreno, G. de Medeiros, E. E. Diel, D. Gault, S. S. Ghosh, I. Gold, Y. O. Halchenko, M. Hartley, D. Horsfall, M. S. Keller, M. Kittisopikul, G. Kovacs, A. Küpcü Yoldaş, K. Kyoda, A. le Tournoulx de la Villegeorges, T. Li, P. Liberali, D. Lindner, M. Linkert, J. Lüthi, J. Maitin-Shepard, T. Manz, L. Marconato, M. McCormick, M. Lange, K. Mohamed, W. Moore, N. Norlin, W. Ouyang, B. Özdemir, G. Palla, C. Pape, L. Pelkmans, T. Pietzsch, S. Preibisch, M. Prete, N. Rzepka, S. Samee, N. Schaub, H. Sidky, A. C. Solak, D. R. Stirling, J. Striebel, C. Tischer, D. Toloudis, I. Virshup, P. Walczysko, A. M. Watson, E. Weisbart, F. Wong, K. A. Yamauchi, O. Bayraktar, B. A. Cimini, N. Gehlenborg, M. Haniffa, N. Hotaling, S. Onami, L. A. Royer, S. Saalfeld, O. Stegle, F. J. Theis, J. R. Swedlow, OME-Zarr: a cloud-optimized bioimaging file format with international community support. Histochem Cell Biol 160, 223–251 (2023).

48. Fractal. https://fractal-analytics-platform.github.io/.

49. J. Swift, I. L. Ivanovska, A. Buxboim, T. Harada, P. C. D. P. Dingal, J. Pinter, J. D. Pajerowski, K. R. Spinler, J.-W. Shin, M. Tewari, F. Rehfeldt, D. W. Speicher, D. E. Discher, Nuclear lamin-A scales with tissue stiffness and enhances matrix-directed differentiation. Science 341, 1240104 (2013).

50. N. Gjorevski, M. Nikolaev, T. E. Brown, O. Mitrofanova, N. Brandenberg, F. W. DelRio, F. M. Yavitt, P. Liberali, K. S. Anseth, M. P. Lutolf, Tissue geometry drives deterministic organoid patterning. Science 375, eaaw9021 (2022).

51. N. P. Tallapragada, H. M. Cambra, T. Wald, S. Keough Jalbert, D. M. Abraham, O. D. Klein, A. M. Klein, Inflation-collapse dynamics drive patterning and morphogenesis in intestinal organoids. Cell Stem Cell 28, 1516–1532.e14 (2021).

52. Q. Yang, S.-L. Xue, C. J. Chan, M. Rempfler, D. Vischi, F. Maurer-Gutierrez, T. Hiiragi, E. Hannezo, P. Liberali, Cell fate coordinates mechano-osmotic forces in intestinal crypt formation. Nat Cell Biol 23, 733–744 (2021).

53. E. Hannezo, C.-P. Heisenberg, Mechanochemical Feedback Loops in Development and Disease. Cell 178, 12–25 (2019).

54. M. Aragona, T. Panciera, A. Manfrin, S. Giulitti, F. Michielin, N. Elvassore, S. Dupont, S. Piccolo, A mechanical checkpoint controls multicellular growth through YAP/TAZ regulation by actin-processing factors. Cell 154, 1047–1059 (2013).

55. A. Jiménez, J. Cotterell, A. Munteanu, J. Sharpe, A spectrum of modularity in multi-functional gene circuits. Molecular Systems Biology, doi: 10.15252/msb.20167347 (2017).

56. M. Cohen, M. Georgiou, N. L. Stevenson, M. Miodownik, B. Baum, Dynamic filopodia transmit intermittent Delta-Notch signaling to drive pattern refinement during lateral inhibition. Dev Cell 19, 78–89 (2010).

57. A. Geling, H. Steiner, M. Willem, L. Bally-Cuif, C. Haass, A gamma-secretase inhibitor blocks Notch signaling in vivo and causes a severe neurogenic phenotype in zebrafish. EMBO Rep. 3, 688–694 (2002).

58. F. Zanconato, M. Forcato, G. Battilana, L. Azzolin, E. Quaranta, B. Bodega, A. Rosato, S. Bicciato, M. Cordenonsi, S. Piccolo, Genome-wide association between YAP/TAZ/TEAD and AP-1 at enhancers drives oncogenic growth. Nat. Cell Biol. 17, 1218–1227 (2015).

59. P. N. P. Singh, S. Madha, A. B. Leiter, R. A. Shivdasani, Cell and chromatin transitions in intestinal stem cell regeneration. Genes Dev. 36, 684–698 (2022).

60. T. Stuart, A. Srivastava, S. Madad, C. A. Lareau, R. Satija, Single-cell chromatin state analysis with Signac. Nat. Methods 18, 1333–1341 (2021).

61. I. Burtscher, H. Lickert, Foxa2 regulates polarity and epithelialization in the endoderm germ layer of the mouse embryo. Development 136, 1029–1038 (2009).

62. D. Z. Ye, K. H. Kaestner, Foxa1 and Foxa2 control the differentiation of goblet and enteroendocrine L- and D-cells in mice. Gastroenterology 137, 2052–2062 (2009).

63. R. Kageyama, T. Ohtsuka. The Hes gene family: repressors and oscillators that orchestrate embryogenesis. Development 134, 1243–1251 (2007)

64. J. Cai, N. Zhang, Y. Zheng, R. F. de Wilde, A. Maitra, D. Pan, The Hippo signaling pathway restricts the oncogenic potential of an intestinal regeneration program. Genes & development 24 (2010).

65. V. S. W. Li, H. Clevers, Intestinal Regeneration: YAP—Tumor Suppressor and Oncoprotein? Current Biology 23, R110–R112 (2013).

66. E. R. Barry, T. Morikawa, B. L. Butler, K. Shrestha, R. de la Rosa, K. S. Yan, C. S. Fuchs, S. T. Magness, R. Smits, S. Ogino, C. J. Kuo, F. D. Camargo, Restriction of intestinal stem cell expansion and the regenerative response by YAP. Nature 493, 106–110 (2012).

67. J. Zhao, M. L. Perkins, M. Norstad, H. G. Garcia, A bistable autoregulatory module in the developing embryo commits cells to binary expression fates. Curr Biol 33, 2851–2864.e11 (2023).

68. W. Xiong, J. E. Ferrell Jr, A positive-feedback-based bistable “memory module” that governs a cell fate decision. Nature 426, 460–465 (2003).

## Supplementary references for SI Theory

1. Collier JR, Monk NAM, Maini PK, Lewis JH. Pattern formation by lateral inhibition with feedback: A mathematical model of delta-notch intercellular signalling. Journal of Theoretical Biology. 1996;183(4):429–446. doi:10.1006/jtbi.1996.0233.

2. Sprinzak D, Lakhanpal A, Lebon L, Santat LA, Fontes ME, Anderson GA, Garcia-Ojalvo J, Elowitz MB. Cis-interactions between Notch and Delta generate mutually exclusive signalling states. Nature. 2010;465(7294):86–90. doi:10.1038/nature08959.

3. Corson F, Couturier L, Rouault H, Mazouni K, Schweisguth F. Self-organized Notch dynamics generate stereotyped sensory organ patterns in Drosophila. Science. 2017;356(6337). doi:10.1126/science.aai7407.

4. Virtanen P, Gommers R, Oliphant TE, et al. SciPy 1.0: fundamental algorithms for scientific computing in Python. Nature Methods. 2020;17(3):261–272. doi:10.1038/s41592-019-0686-2.

5. Totaro A, Castellan M, Battilana G, Zanconato F, Azzolin L, Giulitti S, Cordenonsi M, Piccolo S. YAP/TAZ link cell mechanics to Notch signalling to control epidermal stem cell fate. Nat Commun. 2017;8(1):15206. doi:10.1038/ncomms15206.

6. Shue YT, Drainas AP, Li NY, Pearsall SM, Morgan D, Sinnott-Armstrong N, Hipkins SQ, Coles GL, Lim JS, Oro AE, Simpson KL, Dive C, Sage J. A conserved YAP/Notch/REST network controls the neuroendocrine cell fate in the lungs. Nat Commun. 2022;13(1):2690. doi:10.1038/s41467-022-30416-2.

7. Zhao J, Perkins ML, Norstad M, Garcia HG. A bistable autoregulatory module in the developing embryo commits cells to binary expression fates. Current Biology. 2023;33(14):2851–2864.e11. doi:10.1016/j.cub.2023.06.060.

8. Gozlan O, Sprinzak D. Notch signaling in development and homeostasis. Development. 2023;150(4):dev201138. doi:10.1242/dev.201138.

9. Nandagopal N, Santat LA, Elowitz MB. Cis-activation in the Notch signaling pathway. eLife. 2019;8:e37880. doi:10.7554/eLife.37880.

## Supplementary references for Material and Methods

1. T. Sato, R. G. Vries, H. J. Snippert, M. van de Wetering, N. Barker, D. E. Stange, J. H. van Es, A. Abo, P. Kujala, P. J. Peters, H. Clevers, Single Lgr5 stem cells build crypt-villus structures in vitro without a mesenchymal niche. Nature 459, 262–265 (2009).

2. N. E. Sanjana, O. Shalem, F. Zhang, Improved vectors and genome-wide libraries for CRISPR screening. Nat Methods 11, 783–784 (2014).

3. B.-K. Koo, D. E. Stange, T. Sato, W. Karthaus, H. F. Farin, M. Huch, J. H. van Es, H. Clevers, Controlled gene expression in primary Lgr5 organoid cultures. Nat Methods 9, 81–83 (2011).

4. K. Namoto, C. Baader, V. Orsini, A. Landshammer, E. Breuer, K. T. Dinh, R. Ungricht, M. Pikiolek, S. Laurent, B. Lu, A. Aebi, K. Schönberger, E. Vangrevelinghe, O. Evrova, T. Sun, S. Annunziato, J. Lachal, E. Redmond, L. Wang, K. Wetzel, P. Capodieci, J. Turner, G. Schutzius, V. Unterreiner, M. Trunzer, N. Buschmann, D. Behnke, R. Machauer, C. Scheufler, C. N. Parker, M. Ferro, A. Grevot, A. Beyerbach, W.-Y. Lu, S. J. Forbes, J. Wagner, T. Bouwmeester, J. Liu, B. Sohal, S. Sahambi, L. E. Greenbaum, F. Lohmann, P. Hoppe, F. Cong, A. W. Sailer, H. Ruffner, R. Glatthar, B. Humar, P.-A. Clavien, M. T. Dill, E. George, J. Maibaum, P. Liberali, J. S. Tchorz, NIBR-LTSi is a selective LATS kinase inhibitor activating YAP signaling and expanding tissue stem cells in vitro and in vivo. Cell Stem Cell 31, 554–569.e17 (2024).

5. Q. Yang, S.-L. Xue, C. J. Chan, M. Rempfler, D. Vischi, F. Maurer-Gutierrez, T. Hiiragi, E. Hannezo, P. Liberali, Cell fate coordinates mechano-osmotic forces in intestinal crypt formation. Nat Cell Biol 23, 733–744 (2021).

6. N. P. Tallapragada, H. M. Cambra, T. Wald, S. Keough Jalbert, D. M. Abraham, O. D. Klein, A. M. Klein, Inflation-collapse dynamics drive patterning and morphogenesis in intestinal organoids. Cell Stem Cell 28, 1516–1532.e14 (2021).

7. A. Ayyaz, S. Kumar, B. Sangiorgi, B. Ghoshal, J. Gosio, S. Ouladan, M. Fink, S. Barutcu, D. Trcka, J. Shen, K. Chan, J. L. Wrana, A. Gregorieff, Single-cell transcriptomes of the regenerating intestine reveal a revival stem cell. Nature 569, 121–125 (2019).

8. S. Das, S. M. Parigi, X. Luo, J. Fransson, B. C. Kern, A. Okhovat, O. E. Diaz, C. Sorini, P. Czarnewski, A. T. Webb, R. A. Morales, S. Lebon, G. Monasterio, F. Castillo, K. P. Tripathi, N. He, P. Pelczar, N. Schaltenberg, M. De la Fuente, F. López-Köstner, S. Nylén, H. L. Larsen, R. Kuiper, P. Antonson, M. A. Hermoso, S. Huber, M. Biton, S. Scharaw, J.-Å. Gustafsson, P. Katajisto, E. J. Villablanca, Liver X receptor unlinks intestinal regeneration and tumorigenesis. Nature, doi: 10.1038/s41586-024-08247-6 (2024).

9. I. W. McLean, P. K. Nakane, Periodate-lysine-paraformaldehyde fixative. A new fixation for immunoelectron microscopy. J Histochem Cytochem 22, 1077–1083 (1974).

10. D. Serra, U. Mayr, A. Boni, I. Lukonin, M. Rempfler, L. Challet Meylan, M. B. Stadler, P. Strnad, P. Papasaikas, D. Vischi, A. Waldt, G. Roma, P. Liberali, Self-organization and symmetry breaking in intestinal organoid development. Nature 569, 66–72 (2019).

11. G. de Medeiros, R. Ortiz, P. Strnad, A. Boni, F. Moos, N. Repina, L. Challet Meylan, F. Maurer, P. Liberali, Multiscale light-sheet organoid imaging framework. Nat Commun 13, 4864 (2022).

12. N. Gjorevski, M. Nikolaev, T. E. Brown, O. Mitrofanova, N. Brandenberg, F. W. DelRio, F. M. Yavitt, P. Liberali, K. S. Anseth, M. P. Lutolf, Tissue geometry drives deterministic organoid patterning. Science 375, eaaw9021 (2022).

13. J. Moore, D. Basurto-Lozada, S. Besson, J. Bogovic, J. Bragantini, E. M. Brown, J.-M. Burel, X. Casas Moreno, G. de Medeiros, E. E. Diel, D. Gault, S. S. Ghosh, I. Gold, Y. O. Halchenko, M. Hartley, D. Horsfall, M. S. Keller, M. Kittisopikul, G. Kovacs, A. Küpcü Yoldaş, K. Kyoda, A. le Tournoulx de la Villegeorges, T. Li, P. Liberali, D. Lindner, M. Linkert, J. Lüthi, J. Maitin-Shepard, T. Manz, L. Marconato, M. McCormick, M. Lange, K. Mohamed, W. Moore, N. Norlin, W. Ouyang, B. Özdemir, G. Palla, C. Pape, L. Pelkmans, T. Pietzsch, S. Preibisch, M. Prete, N. Rzepka, S. Samee, N. Schaub, H. Sidky, A. C. Solak, D. R. Stirling, J. Striebel, C. Tischer, D. Toloudis, I. Virshup, P. Walczysko, A. M. Watson, E. Weisbart, F. Wong, K. A. Yamauchi, O. Bayraktar, B. A. Cimini, N. Gehlenborg, M. Haniffa, N. Hotaling, S. Onami, L. A. Royer, S. Saalfeld, O. Stegle, F. J. Theis, J. R. Swedlow, OME-Zarr: a cloud-optimized bioimaging file format with international community support. Histochem. Cell Biol. 160, 223–251 (2023).

14. R. Ortiz, G. de Medeiros, A. H. F. M. Peters, P. Liberali, M. Rempfler, “RDCNet: Instance segmentation with a minimalist recurrent residual network” in Machine Learning in Medical Imaging (Springer International Publishing, Cham, 2020)*Lecture notes in computer science*, pp. 434–443.

15. C. Stringer, T. Wang, M. Michaelos, M. Pachitariu, Cellpose: a generalist algorithm for cellular segmentation. Nat. Methods 18, 100–106 (2021).

16. R. Ortiz, G. de Medeiros, A. H.F.M. Peters, P. Liberali, M. Rempfler. RDCNet: Instance segmentation with a minimalist recurrent residual network. arXiv. Oct 2020 arXiv:2010.00991

17. S. van der Walt, J. L. Schönberger, J. Nunez-Iglesias, F. Boulogne, J. D. Warner, N. Yager, E. Gouillart, T. Yu, scikit-image: image processing in Python. PeerJ 2, e453 (2014).

18. W. Huber, V. J. Carey, R. Gentleman, S. Anders, M. Carlson, B. S. Carvalho, H. C. Bravo, S. Davis, L. Gatto, T. Girke, R. Gottardo, F. Hahne, K. D. Hansen, R. A. Irizarry, M. Lawrence, M. I. Love, J. MacDonald, V. Obenchain, A. K. Oleś, H. Pagès, A. Reyes, P. Shannon, G. K. Smyth, D. Tenenbaum, L. Waldron, M. Morgan, Orchestrating high-throughput genomic analysis with Bioconductor. Nat. Methods 12, 115–121 (2015).

19. S. Heinz, C. Benner, N. Spann, E. Bertolino, Y. C. Lin, P. Laslo, J. X. Cheng, C. Murre, H. Singh, C. K. Glass, Simple combinations of lineage-determining transcription factors prime cis-regulatory elements required for macrophage and B cell identities. Mol. Cell 38, 576–589 (2010).

20. D. Machlab, L. Burger, C. Soneson, F. M. Rijli, D. Schübeler, M. B. Stadler, monaLisa: an R/Bioconductor package for identifying regulatory motifs. Bioinformatics 38, 2624–2625 (2022).

21. L. J. T. Kaaij, F. Mohn, R. H. van der Weide, E. de Wit, M. Bühler, The ChAHP complex counteracts chromatin looping at CTCF sites that emerged from SINE expansions in mouse. Cell 178, 1437–1451.e14 (2019).

22. D. Gaidatzis, A. Lerch, F. Hahne, M. B. Stadler, QuasR: quantification and annotation of short reads in R. Bioinformatics 31, 1130–1132 (2015).

23. A. R. Quinlan, I. M. Hall, BEDTools: a flexible suite of utilities for comparing genomic features. Bioinformatics 26, 841–842 (2010).

24. W. J. Kent, C. W. Sugnet, T. S. Furey, K. M. Roskin, T. H. Pringle, A. M. Zahler, D. Haussler, The human genome browser at UCSC. Genome Res. 12, 996–1006 (2002).

25. G. K. Smyth, “limma: Linear Models for Microarray Data” in Bioinformatics and Computational Biology Solutions Using R and Bioconductor (Springer-Verlag, New York, 2005), pp. 397–420.

